# Confronting population models with experimental microcosm data: from trajectory matching to state-space models

**DOI:** 10.1101/2021.09.13.460028

**Authors:** Benjamin Rosenbaum, Emanuel A. Fronhofer

## Abstract

Population and community ecology traditionally has a very strong theoretical foundation with well-known dynamical models, such as the logistic and its variations, and many modification of the classical Lotka-Volterra predator-prey and interspecific competition models. More and more, these classical models are being confronted with data via fitting to empirical time series for purposes of projections or for estimating model parameters of interest. However, using statistical models to fit theoretical models to data is far from trivial, especially for time series data where subsequent measurements are not independent. This raises the question of whether statistical inferences using pure observation error models, such as simple (non-)linear regressions, are biased, and whether more elaborate process error models or state-space models have to be used to address this complexity.

In order to help empiricists, especially researchers working with experimental laboratory populations in micro- and mesocosms, make informed decisions about the statistical formalism to use, we here compare different error structures one could use when fitting classical deterministic ODE models to empirical data. We consider a large range of biological scenarios and theoretical models, from single species to community dynamics and trophic interactions. In order to compare the performance of different error structure models, we use both realistically simulated data and empirical data from microcosms in a Bayesian framework.

We find that many model parameters can be estimated precisely with an appropriate choice of error structure using pure observation error or state-space models, if observation errors are not too high. However, Allee effect models are typically hard to identify and state-space models should be preferred when model complexity increases.

Our work shows that, at least in the context of low environmental stochasticity and high quality observations, deterministic models can be used to describe stochastic population dynamics that include process variability and observation error. We discuss when more complex state-space model formulations may be required for obtaining accurate parameter estimates. Finally, we provide a comprehensive tutorial for fitting these models in R.

**Open research:** Code for stochastic individual-based simulations is available from https://doi.org/10.5281/zenodo.5500442. A tutorial for fitting ODE models to time series data in R is presented in the Supplementary Information and is also available online https://github.com/benjamin-rosenbaum/fitting deterministic population models. Data (Fronhofer et al., 2020) will be provided via GitHub and Zenodo.

## Introduction

Studying biotic interactions and measuring their strength in order to understand how ecological systems work is at the heart of scientific ecology. Biotic interactions are at the centre of classical questions in population ecology, such as density regulation (e.g., Sibly et al., 2005), but also in community ecology, including modern coexistence theory (Chesson, 2000; Godwin et al., 2020) and beyond, such as host-parasite (epidemiological parameters) and trophic interactions. Importantly, interaction strengths also provide a bridge between ecology and evolution as biotic interactions directly or indirectly influence fitness. Biotic interactions are therefore components of eco-evolutionary dynamics and feedbacks (Yoshida et al., 2003; Hiltunen et al., 2014).

As a consequence, correct and unbiased estimations of biotic interaction strengths are of great importance, be it using times series from the field (e.g., Sibly et al., 2005) or from laboratory systems (Rosenbaum et al., 2019). The strength of ecology in this context is its important and solid body of theory and the availability of mechanistic models. As a consequence, researchers would like to fit these models to time series data to extract the relevant parameters (e.g., Godwin et al., 2020). This exercise is not always straightforward as models often consist of (coupled) differential equations (ODEs), such as classical models including logistic or other limited local population growth, Lotka-Volterra-type models for interspecific competition and consumer-resource dynamics, or SI-type epidemiological models.

Fitting such models to data is possible via multiple approaches, ranging from “naive” trajectory matching (e.g. Fronhofer and Altermatt, 2015) using nonlinear least-squares or Bayesian approaches (e.g. Nørgaard et al., 2021) to state-space models that allow to explicitly take into account observation and process errors (for a recent overview, see Auger-Méthé et al., 2021). Ecologists and evolutionary biologists faced with these choices may often wonder what the pros and cons of these different approaches are, especially given the varying degrees of complexity of these approaches and technical skills they may require.

Importantly, these choices are not mere technical details since they can impact scientific results and conclusions. For instance, Sibly et al. (2005) used a rather simple likelihood approach to fit the *θ*-logistic model to population census data. The authors concluded that a large array of taxa exhibit similarly shaped density-regulation functions which has important applied consequences for conservation and management. However, Clark et al. (2010) could show that these results are likely flawed due to likelihood ridges (parameter combinations of similar likelihood), a problem which could have been mitigated, for instance, using a Bayesian approach with informed priors.

Making a decision regarding the correct statistical formalism to use is not made easier by the fact that the question includes multiple dimensions of complexity, such as frequentist versus Bayesian approaches, discrete-time versus continuous-time models, stochastic versus deterministic models and, finally, multiple error structures.

Of course, we cannot treat all these dimensions comprehensively here. Regarding the first, we will rely on Bayesian inference because Bayesian models are capable of quantifying uncertainty of estimated parameters exactly, even for nonlinear problems, and they perform well with complex models (Clark et al., 2010). Nevertheless, de Valpine and Hastings (2002) compare state-space models to observation and process error models using a likelihood framework and show that state-space models outperform other options.

In terms of the second axis of complexity, we will focus on continuous-time models. Note that Clark and Bjørnstad (2004) study discrete-time models, including exponential growth and the Ricker model and provide a complementary view to the results we will present below.

Regrading the third axis, our study focuses on deterministic theoretical models, which makes our work most relevant when environmental stochasticity is low, such as for data collected in the context of laboratory experimental populations from microcosms or mesocosms. Applications include, for example, plant growth (Paine et al., 2012), Lotka-Volterra competition and predation (Mühlbauer et al., 2020), bacteria or protist predator-prey systems (Rosenbaum et al., 2019; DeLong and Lyon, 2020), host-pathogen interactions (Lunn et al., 2013), and even terrestrial mesocosm foodwebs (Wootton et al., 2022). Data from the field, which may be heavily impacted by environmental stochasticity (Shoemaker et al., 2020), are beyond the scope of our work, and are likely best analysed using stochastic models (see e.g., Barraquand and Gimenez, 2019, 2021). We also do not account for detection error (see Hefley et al., 2013) or model errors (see Xu et al., 2019).

We will here extensively study the fourth axis of complexity, the choice of error structure (Fig. 1). This choice seems especially important when one intends to fit dynamical models to time series data, since in time series, by default, subsequent measurements are not statistically independent. This implies that, besides observation error, process error may have to be taken into account. Nevertheless, researchers often choose a trajectory-matching approach and account for observation error only because this omission reduces model fitting to a comparatively simple nonlinear regression problem. By contrast, process error-only models are nonlinear autoregressive models, where each observation in time is predicted from the previous one. The most complete treatment of error structure involves using state-space models (Auger-Méthé et al., 2021) which take both sources of error into account. Such state-space models cannot be classified as standard regression problems, since they require the simultaneous estimation of latent states as parameters (estimated true population abundances).

**Figure 1:**
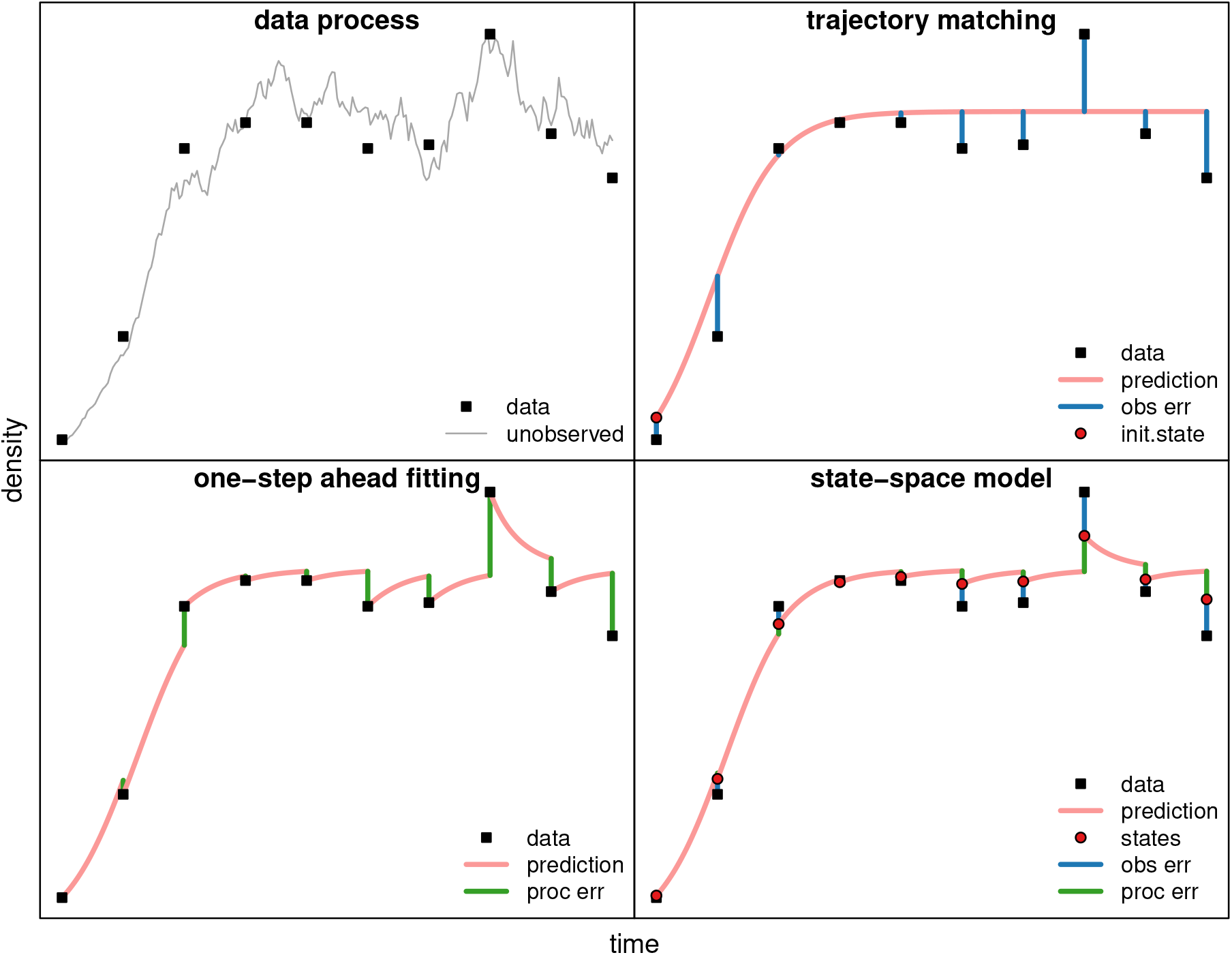
Possible statistical approaches for fitting a population model to a time series of population densities. The upper left panel depicts the underlying stochastic birth-death process (logistic growth) and the black squares show the sampled data with corresponding errors. The simplest statistical model assumes only observation errors and matches the calculated trajectory of the ODE directly to the data (top right). Assuming process error (“one step ahead fitting”; bottom left) takes into account the non-independence of the subsequent data points in the time series. Finally, the state-space model (bottom right) takes both sources of error (observation and process) into account.

In order to understand under which conditions observation error, process error or both have to be modelled when analysing population and community dynamics data with ODE models, we consider ecological scenarios of increasing complexity, ranging from single species dynamics up to predator-prey systems. Briefly, we use stochastic individual-based models to generate observations with known underlying processes and sampling regimes following a “virtual ecologist” approach (Zurell et al., 2010). Specifically, (1) we choose an ecological model with known parameters and (2) simulate trajectories of population abundances with a stochastic algorithm (Gillespie, 2007). Each “experiment” has a duration of 14 days and is replicated 10 times. Here, process error is introduced as demographic stochasticity via random birth and death events. We do not include any form of environmental stochasticity in the data-generating process. (3) We simulate realistic observations from each of the 10 replicates by sampling every 12–24 hours, each measurement includes observation error. (4) We subsequently fit the appropriate dynamical equations in R with Markov Chain Monte Carlo (MCMC) sampling using Stan (Stan Development Team, 2018). Here, we use three different statistical approaches by accounting for observation error only (OBS), process error only (PROC), or fitting state-space models accounting for both (SSM). For each ecological scenario as described below, we conducted 10,000 experiments, each, while varying the levels of introduced process and observation error. Finally, we complement our work with an analysis of real population dynamic data from microbial laboratory systems using published data (Fronhofer et al., 2020).

Overall, in our analysis, process error models fare least well and state-space models outperform simpler approaches for biological models as complex as predator-prey systems when data is irregular or incomplete. Additionally, we provide a comprehensive tutorial for fitting these models in R.

## Materials and methods

### Mathematical models

The following ecological models are used for generating the observed data. Example time series are depicted in Fig. S1, see Table S1 for an overview of the parameters.

### Single species dynamics: Logistic growth model

As the simplest single species density regulation model we take the Verhulst (1838) model, that is, the *r − α* formulation of the logistic growth equation (for a detailed discussion of the advantages of this formulation see Mallet, 2012)

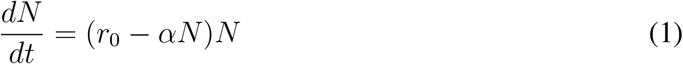

with *N* as the size of the focal population, *r*_0_ as the intrinsic rate of increase and *α* as the intraspecific competition coefficient. The equilibrium population size can be calculated as *K* =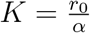

We extend this model to include an Allee effect by adding a density-dependent mortality term as described in Thieme (2003) which can be derived mechanistically, for example, for mate-finding Allee effects or satiating generalist predators:

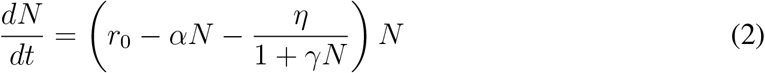

with *η* as the amount by which growth is reduced at *N* = 0 and *γ* controlling the consequences of the Allee effect for higher densities. Calculations of the two equilibrium population densities *A* and *K* (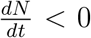 *<* 0 for *N* < *A* and for *N* > *K*) are given in the SupplementaryInformation.

### Single species dynamics: Beverton-Holt model

As a more mechanistic single species density-regulation function (Thieme, 2003; Fronhofer et al., 2020) we explore the continuous-time Beverton-Holt model which follows

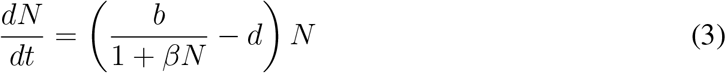

with *b* as the birth rate and *d* as the mortality rate. The intrinsic rate of increase can be calculated as *r*_0_ = *b − d*. The equilibrium population size is *K* =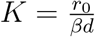

In analogy to Eq. 2 we can expand the Beverton-Holt model to include an Allee effect by adding a mortality terms (Thieme, 2003), which yields

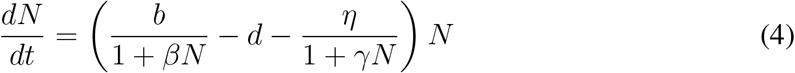

Again we refer to the Supplementary Information for the calculation of the equilibrium population densities *A* and *K*.

### Interspecific competition: *n*-species Lotka-Volterra competition model

To capture interspecific competition and the dynamics of a horizontal community, we will first use an expansion of the logistic model (Eq. 1):

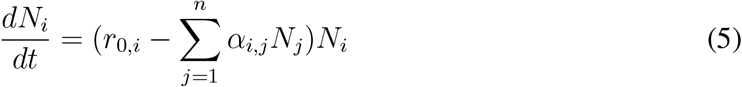

where *α*_*i,j*_ represent the inter- and intraspecific competition coefficients and form the community matrix. We investigate two separate two-species scenarios, which are different in their interspecific competition coefficients. In the first scenario, the system reaches a stable equilibrium (coexistence), while in the second one species is outcompeted by the other and goes extinct (competitive exclusion).

### Predator-prey interactions

For predator-prey interactions, we use the following general model

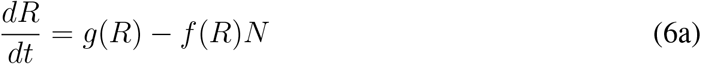

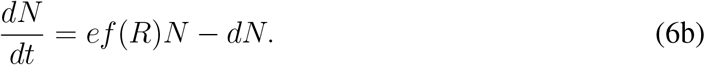

where *g*(*R*) is the growth of the resources which can follow any of the above introduced single species population growth functions (Eqs. 1 – 4). For simplicity we will assume that *g*(*R*) follows Eq. 1. *f* (*R*) is the consumer’s functional response which can be linear, saturating or sigmoid. *e* captures the conversion factor, which includes consumer-resource body-size ratio as well as assimilation efficiency. In our analyses we will focus on a Holling type II (Holling, 1959), that is, saturating functional response of the form 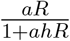with *a* as the predator’s search efficiency and *h* as the handling time. This combination of growth and functional response is also known as the Rosenzweig-MacArthur model and we investigate scenarios that feature a limit cycle (Rosenzweig and MacArthur, 1963).

### Individual-based simulations

In order to simulate the ecological models introduced above, we use an individual-based modelling approach that relies on a modified Gillespie algorithm. The main difference to a classical Gillespie algorithm (exact stochastic simulation algorithm; Gillespie, 2007) consists in calculating maximum rate constants which speeds up the simulation as updating occurs less frequently as detailed in Allen and Dytham (2009). More generally, our approach assumes that birth and death events happen stochastically. Via increasing or decreasing birth and death rates but keeping the resulting intrinsic rate of increase constant (*d* ∈ [0, 1], *b* = *r*_0_ + *d, r*_0_ = 0.1 [h^*−*1^]), we can simulate biologically relevant increases or decreases in demographic stochasticity and therefore process error *σ*_proc_. This approach has the advantage that we do not need to assume any scaling of demographic stochasticity with population size a priori.

The correct scaling emerges from our model. For predator-prey dynamics, we treat the resource’s intrinsic growth rate as above. The predator’s rate of change (i.e., conversion factor *e* and mortality rate *d*) is simultaneously varied. Stochasticity here increases the system’s frequency, but keeps equilibria 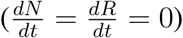 = 0) intact.

We initially verified computationally that a stochastic time series’ variation *σ*^2^ around its deterministic counterpart’s equilibrium *K* scales with 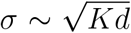 (Fig. S15). Hence we identify *σ*_proc_ with 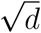 for fixed *K*. For example, with the logistic growth model we observed on average 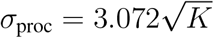 for *d* = 1.0.

Finally, we sample from the generated time series at varying time intervals to include observation error (Fig. 1). Concretely, here the first three days, sampling occurs every 12 hours, then every 24 hours until 14 days. We count abundances *N*_count_ in a fixed fraction *p* of the space monitored or the faction of volume sampled. Assuming a random distribution of individuals in space, this is described by a binomial sampling process *N*_count_ *∼* Binomial(*N, p*). This implies that the estimated abundance 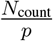 in the total volume has a mean of 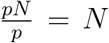 and a standard deviation of 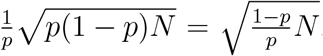. Hence we identify *σ*_obs_ with 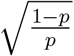 for fixed*N*. For example, any population at its carrying capacity *N* = *K* features an expected observation error 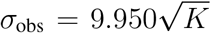 for *p* = 0.01, which is approximately three times higher than the process error as described above.

For each scenario, that is, each population or community model, we simulated 10,000 microcosm experiments. Each experiment was assigned a fixed process error and a fixed observation error (death rate *d* ∈ [0.0, 1.0] and sampling fraction *p* ∈ [0.01, 1.0], random draws from a uniform distribution, for *p* on logscale). In each experiment we simulated 10 time series replicates with identical parameterisation. In the two-species scenarios, 5 additional replicates for the two single-species dynamics were added. This additional single-species data, in conjunction with the 10 two-species time series, informs intra-specific model parameters like *r*_0,*i*_ and *α*_*i,i*_ (*i* = 1, 2).

### Empirical data example

In order to confront our statistical approach with empirical data, we complemented the above described simulations by using population times series data from a microbial laboratory system.

This dataset represents a empirical example that matches our single species model examples introduced above (Eqs. 1 and 3). Furthermore, it fulfils the prerequisites of low environmental stochasticity discussed in the introduction. Nevertheless, it provides room for testing whether and how the different statistical models can pick up non-linearities in density-regulation (Fronhofer et al., 2020).

More precisely, we used the data collected by Fronhofer et al. (2020) from microcosms of the freshwater protist *Tetrahymena thermophila*. These cultures were grown from low density in volumes of 20 mL. Data was collected using a computer-vision and video-analysis pipeline on sample volumes of 31 µL. For details see Fronhofer et al. (2020).

### Statistical approach

Here, we describe the statistical approaches for analyzing our simulated and empirical data. In summary, we combine a deterministic prediction model *U* (*t*) with a statistical model to fit it to an observed time series *Y*_*i*_ at times *t*_*i*_ (*i* = 1, …, *n*) and to estimate model parameters *θ*. Our three proposed statistical models vary in their treatment of observation and process error, that is, their variance structure. Mathematically, they differ in the way *U*_*i*_ = *U* (*t*_*i*_) is predicted from information at the previous time point *t*_*i−*1_: either from the previous prediction *U*_*i−*1_ (OBS), the previous observation *Y*_*i−*1_ (PROC), or a previous latent state variable *Z*_*i−*1_ (SSM). Each model is presented in a separate section below.

More specifically, let *Y*_*i*_ denote untransformed observations (counted abundances in sampling fraction *p*) of a single time series. Estimated abundances in the total volume are given by 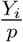. Each observation is univariate for single species systems (*Y*_*i*_ *∈* ℕ), or bivariate for two-species systems (*Y*_*i*_ *∈* ℕ^2^). Generally, we use *m* = 10 non-aggregated time series replicates denoted by *Y*_*ij*_ (*i* = 1, …, *n, j* = 1, …, *m*). In the following, we describe the model fitting of single time series for simplicity and briefly explain extensions to multiple replicates where needed. *U* (*t*) is a deterministic prediction, or process model, for the population time series in the total volume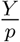. Here we use *U* (*t*) = *U* (*t*|*t*_1_, *U*_1_, *θ*), the numerical solution of the continuous-time ODE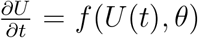= *f* (*U* (*t*), *θ*) with initial value *U* (*t*_1_) = *U*_1_ and model parameters *θ*. A discrete-time process model *U* (*t*) would work analogously.

We use a Bayesian approach for parameter estimation, but the models can generally be fitted with maximum likelihood estimation as well (e.g. Xu et al., 2019; DeLong and Lyon, 2020). Both methodologies require the evaluation of the likelihood function 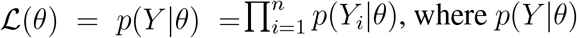 where *p*(*Y* |*θ*) denotes the probability density function of the observed data *Y* given the model parameters *θ*.

### Observation error model (OBS)

When ignoring process error, the whole trajectory

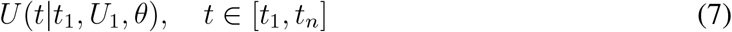

is computed, yielding predictions *U*_1_, …, *U*_*n*_. The initial abundance *U* (*t*_1_) = *U*_1_ is also a free parameter to be estimated. Eq. 7 is identical to

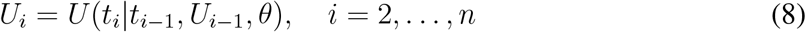

and predictions *U*_*i*_ are (iteratively) defined by initial state *U*_1_ and model parameters *θ* (Fig. 1b). The predictions *p · U*_*i*_ for the sampled fraction *p* are confronted with the data *Y*_*i*_ by evaluating the likelihood ℒ (*θ, U*_1_). For the likelihood, we chose a negative binomial distribution

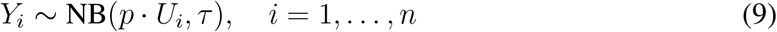

which has mean *µ* = *p* · *U*_*i*_ and variance 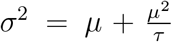 Parameter *τ* > 0 describes the *τ* amount of overdispersion which makes the negative binomial a flexible choice, both for error variances scaling with *µ* (large *τ*) or with *µ*^2^ (small *τ*). Also, it is suited for the integer- and potentially zero-valued observations *Y*_*i*_ (O’Hara and Kotze, 2010). For time series that did not include any zeros as our empirical data example, we found that a lognormal distribution works as well. By neglecting process error and assuming that the process is sufficiently described by a deterministic trajectory, parameter estimation reduces to a nonlinear regression problem, fitting *U* (*t*) (Eq. 7) to observations *Y*_*i*_ with independent residuals (Eq. 9).

In case of multiple time series replicates (observations *Y*_*ij*_), we fitted *m* trajectories *U* (*t*|*t*_1_, *U*_1*j*_, *θ*) using a single set of model parameters (e.g. *θ* = (*r*_0_, *K*), logistic growth scenario), but allowing individual initial values *U*_1*j*_ (*j* = 1, …, *m*) in one statistical model, i.e. using a joint likelihood function ℒ (*θ, τ, U*_11_, …, *U*_1*m*_).

### Process error model (PROC)

When observation error is ignored, we assume that the observations for the total volume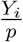 are sufficiently close to the true abundances. Predictions are generated one-step-ahead

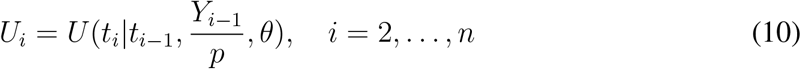

i.e. predicting *U*_*i*_ from the previous observation *Y*_*i−*1_ only (Fig. 1c). The observations’ deviation from this piecewise deterministic process is modeled again with a negative binomial distribution

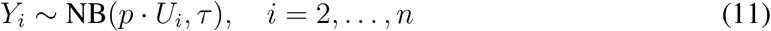

for reasons stated above, but here it accounts for process error. Thus, by neglecting observation error, parameter estimation reduces to a nonlinear autoregressive problem with independent residuals.

For multiple time series replicates, we straightforwardly iterate Eqs. 10,11 for *Y*_*ij*_ and *U*_*ij*_ (*j* = 1, …, *m*) with a single set of model parameters *θ*, to evaluate the joint likelihood ℒ (*θ, τ*) in one statistical model.

### State-space model (SSM)

This approach assumes both observation error and process error are present. It requires explicitly modelling the time series of estimates of true abundances *Z*_*i*_ (*i* = 1, …, *n*) as latent states. These are unknown a-priori, but can be estimated together with the model parameters *θ* from the data during the model fitting process. For each “guess” of *Z* and *θ*, predictions are generated one-step-ahead

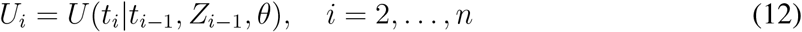

from the previous states *Z*_*i−*1_ (Fig. 1d). The process error in these predictions is modelled with a lognormal distribution

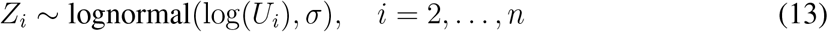

with a scale parameter *σ*. Hence, log(*Z*_*i*_) follows a normal distribution with mean log(*U*_*i*_) and standard deviation *σ*. A continuous distribution is required here for *U*_*i*_, *Z*_*i*_ *∈* ℝ. Additionally, the state parameters *Z*_*i*_ are confronted with the integer-valued observations *Y*_*i*_ with a negative binomial distribution as in the previous models

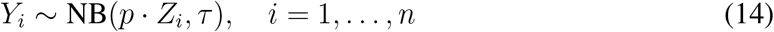

accounting for observation error. Eqs. 13,14 define the likelihood function ℒ (*θ, σ, τ, Z*_1_, …, *Z*_*n*_). When dealing with multiple time series replicates *Y*_*ij*_ (*j* = 1, …, *m*), we fit *m* individual time series of true states *Z*_1*j*_, …, *Z*_*nj*_ with a joint set of *θ*, using the likelihood function ℒ (*θ, σ, τ, Z*_11_, …, *Z*_*nm*_) in one statistical model.

### Parameter estimation

We used MCMC to sample from the posterior probability distribution of the model parameters given the observations *P* (*θ*|*Y*) *∼ P* (*Y* |*θ*) · *P* (*θ*), where *P* (*Y* |*θ*) = ℒ (*θ*) is the likelihood function and *P* (*θ*) denotes some prior distribution for the model parameters *θ*. We coded the models using the “rstan” package (Stan Development Team, 2018) and used the built-in Runge-Kutta method for numerical solutions of ODE predictions *U* (*t*), and the no-u-turn sampler for computing the posterior. Vague or uninformative prior distributions *P* (*θ*) were chosen for all model parameters to guarantee that the measured model performance was not confounded with prior information (Table S1).

Each model fit was computed by 2000 warmup steps and 2000 samples in three chains, adding up to 6000 posterior samples. We discarded every dataset from subsequent analysis if either of the three fitting methods did not converge (more than 100 divergent iterations).

### Evaluation

For each of the 10,000 model fits per scenario, we computed bias and root mean squared error (RMSE) for every parameter *θ*_*i*_ from the posterior distribution 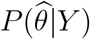 to evaluate accuracy, where 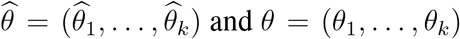 and *θ* = (*θ*_1_, …, *θ*_*k*_) denote the *k* estimated and true model parameters, respectively (e.g. *θ*_1_ = *r*_0_, *θ*_2_ = *K* for logistic growth). Relative bias

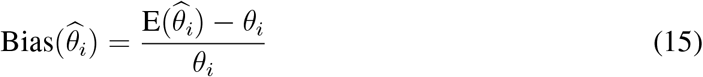

is a measure of the point estimate (posterior mean F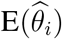, while the relative RMSE

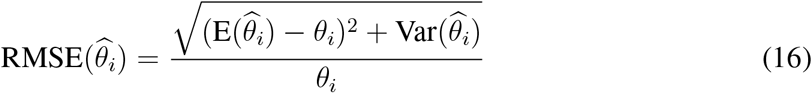

also accounts for the uncertainty of the estimation (by including the posterior variance 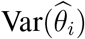Additionally, we investigated the effects of process and observation error on model accuracy to test, e.g., if the statistical approaches OBS, PROC and SSM are affected differently by the amount of both error sources in the data. First, for each fit, we computed overall measures of accuracy as geometric means 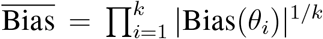 and 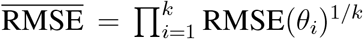over absolute Bias and RMSE of all model parameters *θ*_*i*_, respectively. Then, we fitted generalized additive models (GAMs) to these measures using observation error and process error as predictors and visualized the response surfaces, see Figs. S2–S13. We used the “mgcv” package (Wood, 2011) with model formulae 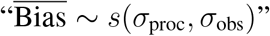 and 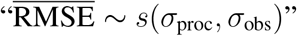

## Results

### Fitting simulated data

When analyzing time series with OBS and PROC, almost all fits converged properly and only less than 1% of all experiments had problems with divergent iterations. For SSM, around 5% of all fits did not converge, with a maximum of 9% for our most complex model (predatorprey system). There are generic strategies when encountering convergence issues, like employing more informative priors, using a longer warmup phase for the MCMC sampler, or increasing the sampler’s accuracy at the cost of computation speed (Stan Development Team, 2022), which we did not further investigate case-by-case given the amount of involved experiments.

In most scenarios, OBS and SSM performed best in terms of the model parameters’ bias (Fig. 2) and RMSE (Fig. 3). In cases where models were identifiable via SSM, OBS performed comparably or just slightly less accurately, while PROC mostly produced the least accurate parameter estimates.

**Figure 2:**
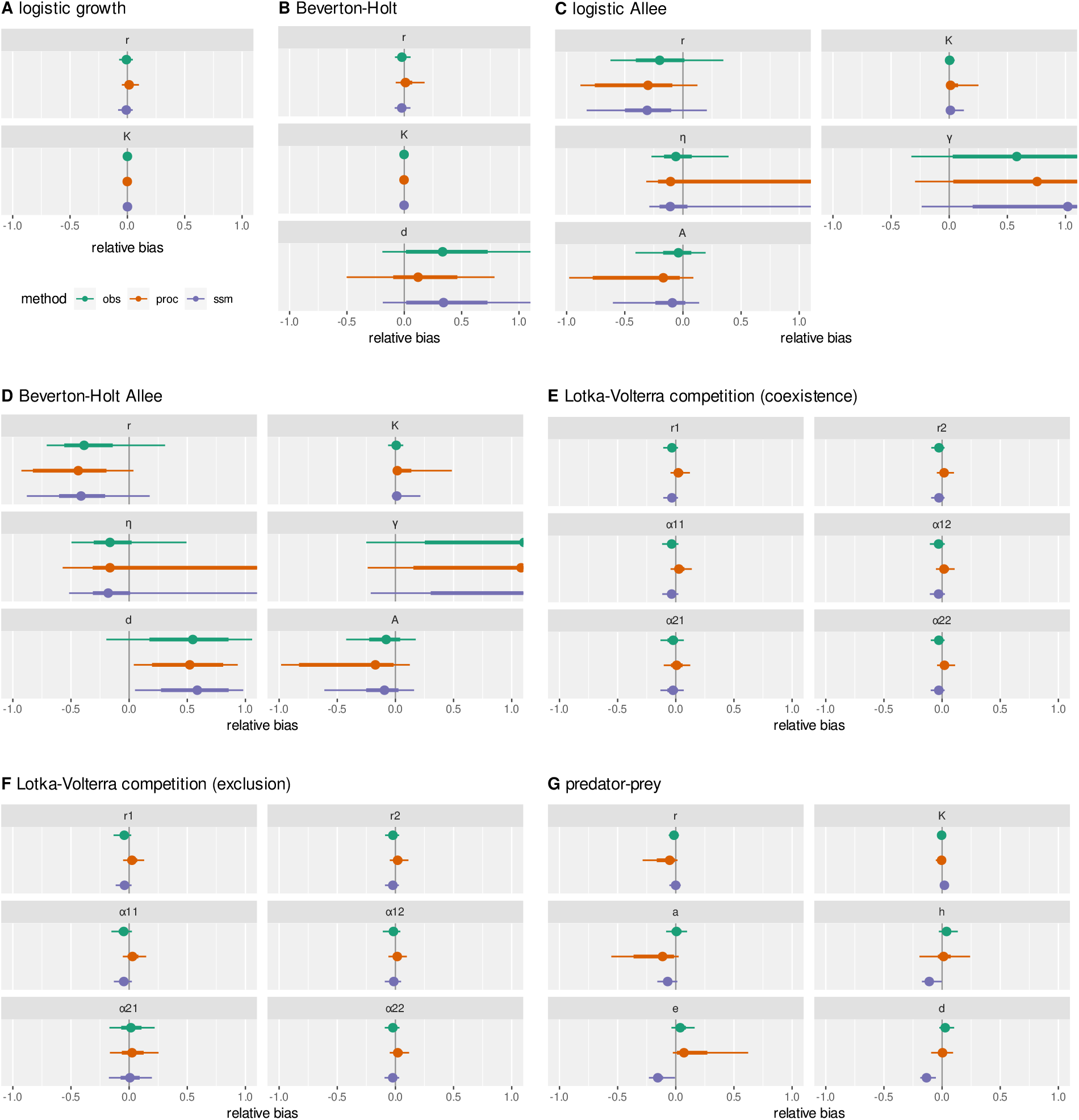
Distribution of relative bias, model parameters vs. statistical models for all population dynamics models (10,000 experiments per scenario). Dots are mean, thin lines are 95%, and bold lines are 66% quantiles. For Allee models, *A* is computed from other model parameters and not a free parameter itself.

**Figure 3:**
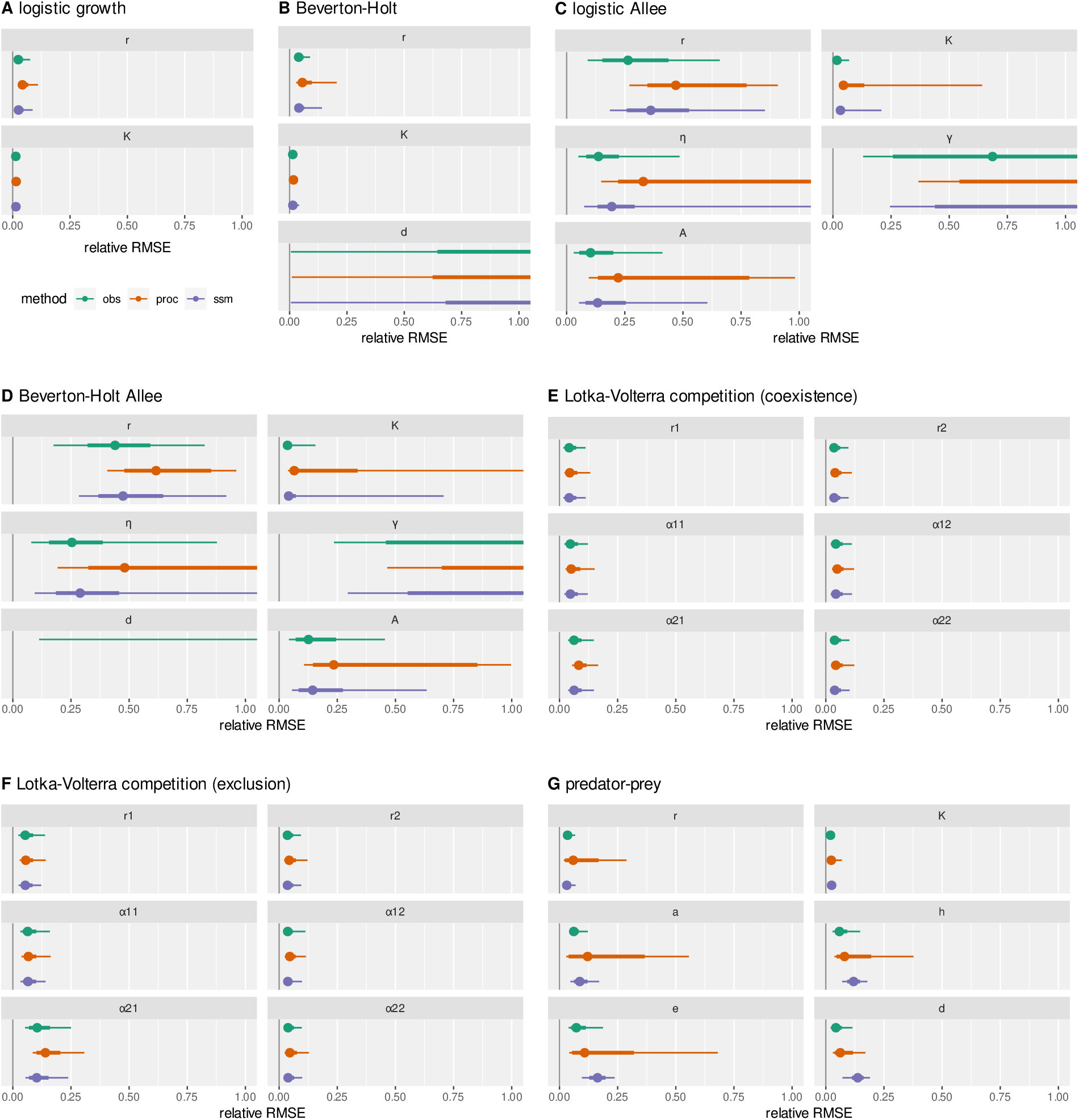
Distribution of relative RMSE, model parameters vs. statistical models for all population dynamics models (10,000 experiments per scenario). Dots are mean, thin lines are 95%, and bold lines are 66% quantiles. For Allee models, *A* is computed from other model parameters and not a free parameter itself.

For the **logistic growth** model (Eq. 1), all models produced unbiased estimates. PROC estimates of the growth rate *r* were slightly less precise, especially for high observation error (Fig. S2).

In the **Beverton-Holt** model (Eq. 3), parameter *d* defines the mortality rate as well as the level of process error in the individual-based simulations 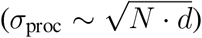. Further, with increasing

*d*, the density-regulation function converges to the linear version of logistic growth and time series resemble those of the logistic growth model. In our parametrization, a nonlinear effect in the density-regulation function on the stochastic time series was only visible for *d <* 0.4 (approximately). Therefore, the exact values of large *d* were not identifiable, which caused high variation in the estimates and their uncertainty. But if the data provided evidence for the Beverton-Holt model vs. logistic growth (*d* small), then all three statistical models provided accurate estimates (Fig. S3).

The parameter estimates of the **logistic growth with Allee effect** model (Eq. 2) were mostly biased. OBS and SSM still produced more accurate estimates than PROC. Its estimates were heavily biased if not both process error and observation error were low (Fig. S4). The inaccuracy was most pronounced for high observation error. All statistical models suffered from highly correlated parameters *r, η* and *γ* in the posterior distribution (of a single fit). We emphasize that this is not a problem with MCMC convergence, but rather a problem with practical identifiability (Raue et al., 2009). Information on this nonlinear, four-parameter population growth rate is lost due to process and observational noise, and cannot be recovered accurately by statistical inference. However, the critical population density *A*(*r, K, η, γ*) (see Supplementary Information), which separates positive and negative population growth, was estimated comparably accurately by OBS and SSM.

With one additional parameter *d* for mortality, the **Beverton-Holt with Allee effect** model (Eq. 4) featured problems similar to the Beverton-Holt model and the logistic Allee effect model. Interestingly, model performance suffered for low *d*, which is associated with low process error and strong Beverton-Holt density-regulation (Fig. S5). We assume that the two nonlinear density-effects cannot be distinguished accurately by any of the statistical models, and that this five parameter model is practically not identifiable.

We tested a two-species **Lotka-Volterra competition** (Eq. 5) model in a **coexistence** scenario (both species reached a positive steady state) using additional control data (single species time series, each growing to their carrying capacity). The accuracy level was generally high (Fig. S6). Without control data, results were similar albeit slightly less accurate. Especially PROC performed worse under the presence of observation error (Fig. S9).

In a **competitive exclusion** scenario (one species was outcompeted and went extinct, while the other one grew to its carrying capacity) using additional control data, accuracy levels generally were high and just slightly lower than in the coexistence scenario. OBS and SSM produced marginally better estimates than the PROC approach. Estimation errors grew with observation error (Fig. S7), especially for interspecific competition coefficient *α*_21_ of the out-competed species 1. Without control data however, estimation accuracy decreased in total (Fig. S10), partially leading to biased estimates for all three fitting approaches. Without data on the carrying capacity *K*_1_ =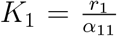 of the outcompeted species, model identifiability suffered, and most strikingly for PROC.

We tested a **predator-prey** model (Eq. 6a, 6b) with cyclic dynamics. The parameters were chosen such that the system experienced approximately two to three full cycles in the observed time (depending on the level of process error). First, we used additional single-species control time series (resource growing to its carrying capacity, consumer going extinct). SSM and OBS provided mostly unbiased estimates, with slight underestimation of predator’s model parameters for SSM. Here, the posterior distributions of the individual fittings showed correlations in these parameters. PROC generally featured biased estimates and was highly sensitive to observation error (Fig. S8). Second, we fitted the model without any control data (Fig. S11). While accuracy decreased for parameters estimated with OBS and PROC, SSM results still were comparable to the estimates with control data when observation error was not too high.

### Additional scenarios

The OBS and the SSM fitting approach performed quite well in some two-species scenarios, even under the presence of process and observation error. We repeated the analysis of the Lotka-Volterra competition model and the predator-prey model to validate the methods under more challenging conditions.

We fitted the **Lotka-Volterra competition** model using **fewer replicates**, i.e. only two instead of ten time series replicates of the two-species mixtures, and only one instead of five single-species control time series each. Our assumption was that, even though process error can speed up or slow down dynamics, these effects would average out over ten replicates and would inform parameters (e.g. the joint average growth rate) correctly. We found that this generally holds even with fewer replicates, but results were slightly less accurate regarding bias (Fig. S12). However, fewer observations also meant higher posterior uncertainty and therefore higher RMSE, especially for *α*_21_, PROC performed worst here. In particular, OBS estimates were as accurate as SSM estimates across all levels of observation and process error. We also fitted **predator-prey** models to **longer time series** over 35 days instead of 14 days, such that the system featured approximately five to seven full cycles. Additional single-species control time series were used as before. Our assumptions were that process error did not affect the estimation quality significantly in the shorter time series, since the regularity of the process (constant cyclic frequency) was not seriously disturbed, and that it would only decrease with longer time series, leading to less accurate results especially for the OBS model.

Fitting longer time series (Fig. S13), results for OBS were indeed slightly less precise than for the shorter time series, while SSM results did not change. We conclude that, while OBS is still quite robust against the level of process variation and the resulting irregularity tested here, SSM should be used for longer time series with irregular cycles.

Finally, we simulated experiments for **logistic growth with a lower carrying capacity** (*K* = 10^3^ instead of *K* = 10^4^). Generally, process and observation error scale with square root of population abundance as verified above 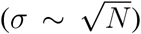, therefore relative errors decrease with abundance 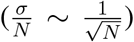. As expected, the accuracy of estimated parameters descreased in experiments with lower abundances (Fig. S14), but PROC suffered most.

### Fitting an empirical data set

We fitted the Beverton-Holt model (Eq. 3) with all three statistical approaches to seven empirical datasets, comprised of six time series replicates each (Fig. 4, see Fig. S16 for full dataset and Figs. S17–S23 for posterior predictive checks). Since all observed densities were positive and non-integer, we used lognormally distributed residuals for process and observation errors. Note that densities are non-integer here because the data was collected using video-recording and analysis and densities are calculated as averages over the observation time of one video. Nonlinear effects on the density-regulation function were detected in datasets 4,

**Figure 4:**
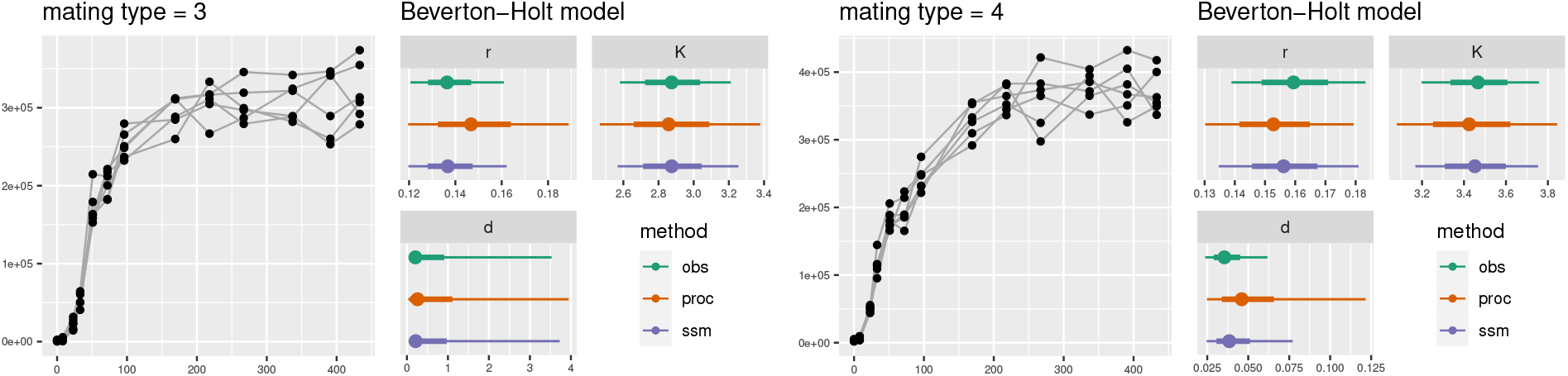
Two out of seven empirical datasets and posterior distributions of fitted Beverton-Holt models, including model parameters *r, K, d*. For parameter *K*, x-axis labels show *K/*10^5^. Left: *d* unidentifiable indicating logistic growth, right: small *d* identifiable.

6 and 7 by all three statistical models, indicated by low values of mortality rate *d*. For the remaining datasets, larger values of *d* were estimated with a high uncertainty in the posterior distributions, which suggested no clear evidence for the Beverton-Holt model over the logistic growth model. Estimates of the three model parameters *r, K* and *d* were similar across the three statistical approaches, but posterior uncertainty was generally higher in the PROC estimates.

## Discussion

Our work shows that using deterministic ODEs and Bayesian inference, it is possible to accurately estimate parameters from time series data of stochastic IBMs including process and observation error. Practical identifiability, which is based both on the model and data quality (Raue et al., 2009), was validated for several widely-used population models with respect to the different statistical models used here. Importantly, our work covers multiple dimensions of complexity, both in terms of population ecology and statistical models, from simple single species models, such as the logistic model, to multispecies community models, and, in terms of statistical models, from pure observation (OBS), via process error (PROC) to state-space models (SSM). Overall, we can show that OBS and SSM can be generally preferred over PROC in the context of deterministic prediction models. It is important to keep in mind that our work is motivated by data from experimental laboratory systems, that is, systems where environmental stochasticity is typically low.

More precisely, OBS can compete with the more complex SSM, especially when a true steady state *>* 0, such as an equilibrium density, is reached (logistic model, Beverton-Holt model if *d* is small and data is not logistic, Lotka-Volterra competition model) or when the biological models lead to regular cycles, such as in some predator-prey dynamics. With additional single-species time series as control data in the predator-prey systems, OBS produced even better estimates than SSM, while the opposite was found when no control data was used. For data featuring irregular cycles (e.g., in longer time series), we also endorse the use of SSMs. The good performance of OBS for predator-prey models is in line with recent studies (Rosenbaum et al., 2019; DeLong and Lyon, 2020). Also, Barraquand and Gimenez (2021) demonstrated with a discrete-time stochastic model, that parameters can be more easily estimated from (noisy) limit cycles than from time series converging to a steady state in general. Therefore, OBS may be a viable alternative to SSM, especially if SSMs suffer, for example, from extensive model complexity or convergence problems (Bolker, 2008). Auger-Méthé et al. (2016) discuss identifiability issues with linear SSMs, especially if measurement error is larger than process error, or that *σ*_obs_ and *σ*_proc_ often cannot be estimated accurately (e.g. Knape, 2008). Here, we consider these as nuisance parameters and focus on model parameters. While we observe a decrease of model performance with measurement error for all three models (Figs. S2–S13), this does not generally reach a level we would consider as unidentifiable for SSMs. Contrary to the studies cited above, which investigate stationary time series, our datasets feature transient or cycling dynamics, containing more information on the nonlinear processes parameters.

We modelled process variation in PROC and SSM by assuming a piecewise deterministic process between sampling times *t*_*i*_ and *t*_*i*+1_ (ODE integration), where error is introduced in *t*_*i*+1_ (Auger-Méthé et al., 2021). A more elaborate treatment of process error, that potentially could increase estimation accuracy, requires either an analytical expression of error propagation in the stochastic population process to compute the likelihood, or stochastic simulations during model fitting for a likelihood approximation. While the former is often not available, the latter requires computationally even more demanding methods like sequential Monte Carlo (SMC) or Approximate Bayesian Computing (ABC) (Hartig et al., 2011). While we acknowledge that SSMs classically use piecewise (log-)linear predictions models instead of ODEs, we here aimed at comparing three ways of statistically treating error structure and used identical continuous prediction models for each. How linearized ODEs (Euler’s method) and temporal resolution (Clark and Bjørnstad, 2004) affect model performance was not tested here.

Our work also shows that estimating Allee effect strengths and more generally fitting population growth models with Allee effects is challenging. For these models, the 4- or even 5-parameter density-regulation functions may be overparameterised and nonidentifiable especially in the presence of process and observation error. As a consequence, these models may only be useful for data from highly controlled experiments in combination with SSMs. It it important to note that these models can be derived mechanistically (Thieme, 2003), which may allow to inform priors of one or multiple model parameters which could help make the SSM more accurate. Of course, as an alternative, non-mechanistic formulations such as 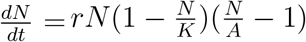 with less parameters could also be used.

In the case of two-species systems we highly recommend the use of control time series from single species settings (Figs. S6 & S9, S7 & S10, S8 & S11). If these are not available, SSMs should be used for predator-prey systems. Biological settings that lead to the exclusion of one species, such as some of the Lotka-Volterra competition models we have used here, make estimates imprecise without such control data, unless the observation error is very small.

While we here explored model identifiability based on time series data alone, a Bayesian approach easily allows including additional sources of information to improve parameter estimation. This information could enter the model via informed priors or hierarchical models (Kindsvater et al., 2018), or by using multiple, potentially heterogeneous data sources (Barraquand and Gimenez, 2019). For example, feeding experiments (Rosenbaum and Rall, 2018) can additionally inform functional response parameters in predator-prey models (Barraquand and Gimenez, 2021).

In conclusion, we have explored multiple dimensions of complexity, both in terms of biological complexity as well as in terms of statistical model complexity, in order to pinpoint which error structure one should use when fitting classical deterministic ODE models to empirical data, from single species to community dynamics and trophic interactions. Our results show that, overall, observation error models and state-space models outperform process error models for data that one may expect to be collected from experimental laboratory populations. Importantly, our continuous-time models allow us to include uneven sampling intervals (and therefore missing values), because the model is not linearised within a time-step as in discrete-time models. More generally, our work shows that under the conditions specified above, deterministic models seem to be sufficient to describe the stochastic dynamics emerging from process and observation errors.

## Supporting information

Tutorial

## Acknowledgements

This is publication ISEM-YYYY-XXX of the Institut des Sciences de l’Evolution – Montpellier. This work was supported by a grant from the Agence Nationale de la Recherche (No.: ANR-19-CE02-0015) to EAF. BR gratefully acknowledges the support of iDiv funded by the German Research Foundation (DFG-FZT 118, 202548816). The scientific results have in part been computed at the High-Performance Computing Cluster EVE of the Helmholtz Centre for Environmental Research (UFZ) and iDiv, and we thank Christian Krause for technical support.

## Authors contributions

BR and EAF conceived the ideas and designed methodology. EAF developed the simulation model and BR developed the statistical analysis framework. BR analysed the data. Both authors contributed to writing the manuscript.

