## Supplement for "Confronting population models with experimental microcosm data: from trajectory matching to state-space models"

#### Supplement Information

Benjamin Rosenbaum[1,2] and Emanuel A. Fronhofer[3]

1. German Centre for Integrative Biodiversity Research (iDiv), Halle-Jena-Leipzig, Germany
2. Institute of Biodiversity, Friedrich Schiller University Jena, Jena, Germany
3. ISEM, Université de Montpellier, CNRS, IRD, EPHE, Montpellier, France

### Reparametrisations

#### Logistic growth models

Solving $r - \alpha N = 0$ (Eq. 1) for $N$ defines the carrying capacity

$$K = \frac{r}{\alpha}.$$

Conversely, the $r - \alpha$ formulation can be parametrized with $K$ using

$$\alpha = \frac{r}{K}.$$

For the logistic Allee model (Eq. 2), we solve

$$r - \alpha N - \frac{\eta}{1 + \gamma N} = 0$$

for $N$, which yields the quadratic equation

$$N^2 + \frac{\alpha - \gamma r}{\alpha\gamma} N + \frac{\eta - r}{\alpha\gamma} = 0.$$

Using $p = \frac{\alpha - \gamma r}{\alpha\gamma}$ and $q = \frac{\eta - r}{\alpha\gamma}$, carrying capacity $K$ and the critical point $A$ are calculated by

$$K = -\frac{p}{2} + \sqrt{\left(\frac{p}{2}\right)^2 - q}$$
$$A = -\frac{p}{2} - \sqrt{\left(\frac{p}{2}\right)^2 - q}.$$

To parameterise the logistic Allee model with a carrying capacity $K$, we set the population model (Eq. 2) to zero, insert $K$, and solve

$$\left(r - \alpha K - \frac{\eta}{1 + \gamma K}\right) K = 0$$

for $\alpha$:

$$\alpha = \frac{1}{K}\left(r - \frac{\eta}{1 + \gamma K}\right).$$

#### Beverton-Holt models

The carrying capacity of the Beverton-Holt model (Eq. 3) is calculated by solving

$$\frac{r+d}{1+\beta N} - d = 0$$

for $N$, then

$$K = \frac{r}{\beta d}.$$

Conversely, the model can be parameterised by a $K$ using

$$\beta = \frac{r}{Kd}.$$

For the Beverton-Holt Allee model (Eq. 3), we solve

$$\frac{b}{1+\beta N} - d - \frac{\eta}{1+\gamma N} = 0$$

for $N$, which yields the quadratic equation

$$N^2 + \frac{d\alpha + d\gamma + \eta\alpha - b\gamma}{d\alpha\gamma} N + \frac{\eta - r}{d\alpha\gamma} = 0.$$

Using $p = \frac{d\alpha+d\gamma+\eta\alpha-b\gamma}{d\alpha\gamma}$ and $q = \frac{\eta-r}{d\alpha\gamma}$, carrying capacity $K$ and the critical point $A$ are calculated by

$$K = -\frac{p}{2} + \sqrt{\left(\frac{p}{2}\right)^2 - q}$$
$$A = -\frac{p}{2} - \sqrt{\left(\frac{p}{2}\right)^2 - q}.$$

To parameterise the Beverton-Holt Allee model with a carrying capacity $K$, we set the population model (Eq. 4) to zero, insert $K$, and solve

$$\left(\frac{b}{1+\beta K} - d - \frac{\eta}{1+\gamma K}\right) K = 0$$

for $\beta$:

$$\beta = \frac{1}{K}\left(\frac{b}{d + \frac{\eta}{1+\gamma K}} - 1\right).$$

### Model parameters and priors

Table S1: Model parameters, their meaning and tested values. For free parameters to be estimated, prior distributions are provided.

| **Parameter** | **Values** | **Meaning** | **Prior** | **Type** |
|---|---|---|---|---|
| $r_0$ | 0.1 | resource intrinsic growth rate | lognormal(-2,1) | vague |
| $d_R$ | $\in [0, 1]$ | resource death rate (also determines process error) | lognormal(0,1) | vague |
| $b$ | $r_0 + d_R$ | resource birth rate | - | |
| $K$ | $10^4$ | resource carrying capacity | lognormal(9,1) | vague |
| | $2 \cdot 10^4$ | resource carrying capacity (predator-prey model) | | |
| $\alpha$ | $\frac{r_0}{K}$ | resource intraspecific competition coefficient (log. model) | - | |
| | $\alpha(r_0, \eta, \gamma, K)$ | resource intrasp. competition coefficient (log. Allee model) | - | |
| $\beta$ | $\frac{r_0}{d_R K}$ | resource intrasp. competition coefficient (B.-H. model) | - | |
| | $\beta(r_0, d_R, \eta, \gamma, K)$ | resource intrasp. competition coefficient (B.-H. Allee model) | - | |
| $\eta$ | 0.125 | Allee effect strength at low densities | normal(0,1) | vague |
| $\gamma$ | 0.0005 | Allee effect strength at high densities | gamma(2,1) | positive |
| $\alpha_{i,j}$ | $\in [0.5 \cdot 10^{-5}, 1.5 \cdot 10^{-5}]$ | resource interspecific competition coefficient | gamma(2,1) | positive |
| $a$ | $3.333 \cdot 10^{-4}$ | consumer search efficiency | gamma(2,1) | positive |
| $h$ | 0.5 | consumer handling time for type II functional response | gamma(2,1) | positive |
| $e$ | $\in [0.05, 0.15]$ | consumer conversion factor | uniform(0,1) | bounded |
| $d_C$ | $\in [0.05, 0.15]$ | consumer death rate | gamma(2,1) | positive |
| $T_{max}$ | 336 h | maximal simulation time | | |
| $R_0$ | $\in [10^2, 10^3]$ | resource starting density | | |
| $C_0$ | 400 | consumer starting density | | |

### Figures

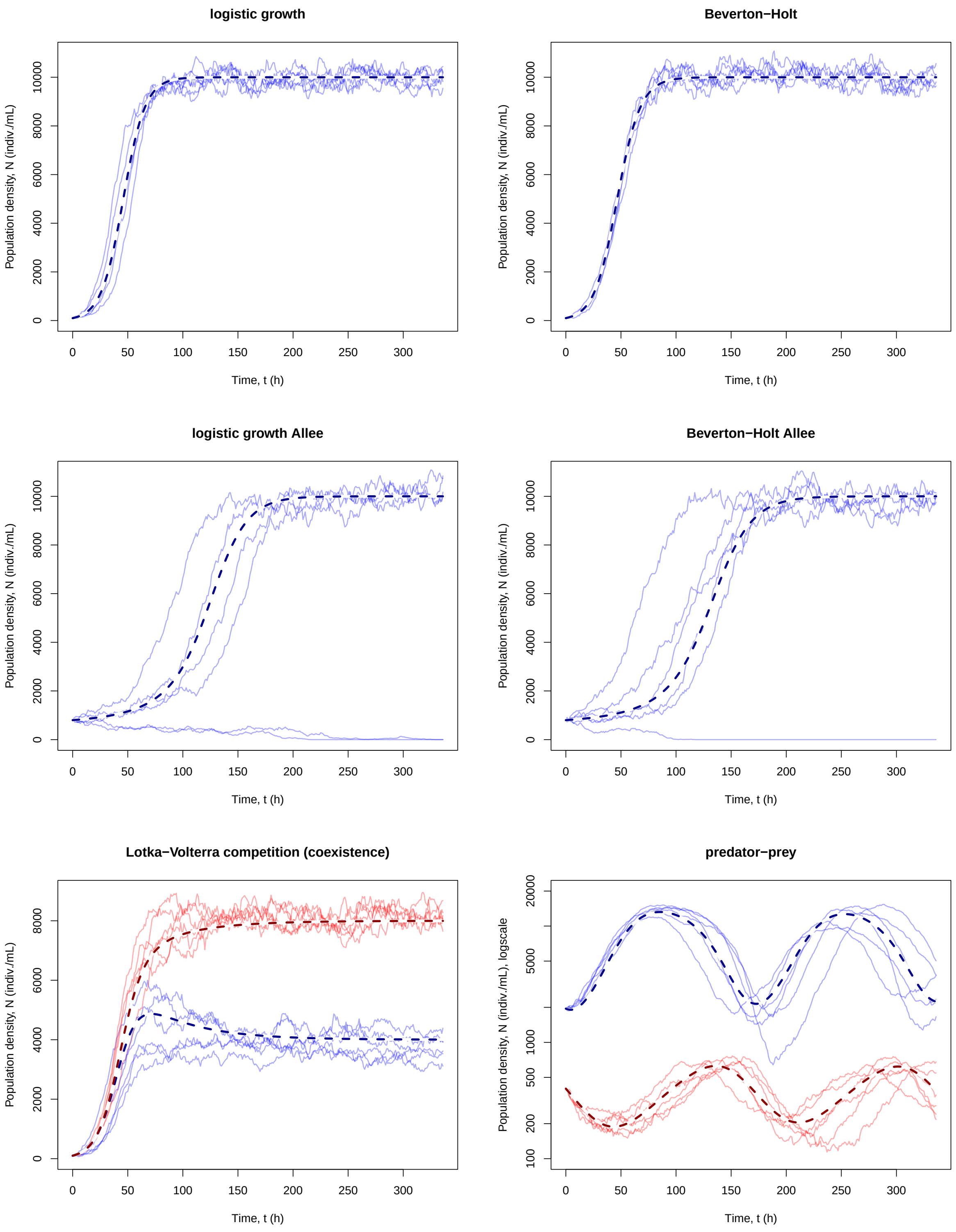


Figure S1: Examples of a deterministic (bold dashed) and 6 replicates of stochastic (thin solid) simulations, each. Maximum process error was used ($d = 1.0$), no observation error. In 2-species-systems (Lotka-Volterra and predator-prey) species are distinguished by color. Y-axis for predator-prey is on logscale.

### Logistic growth

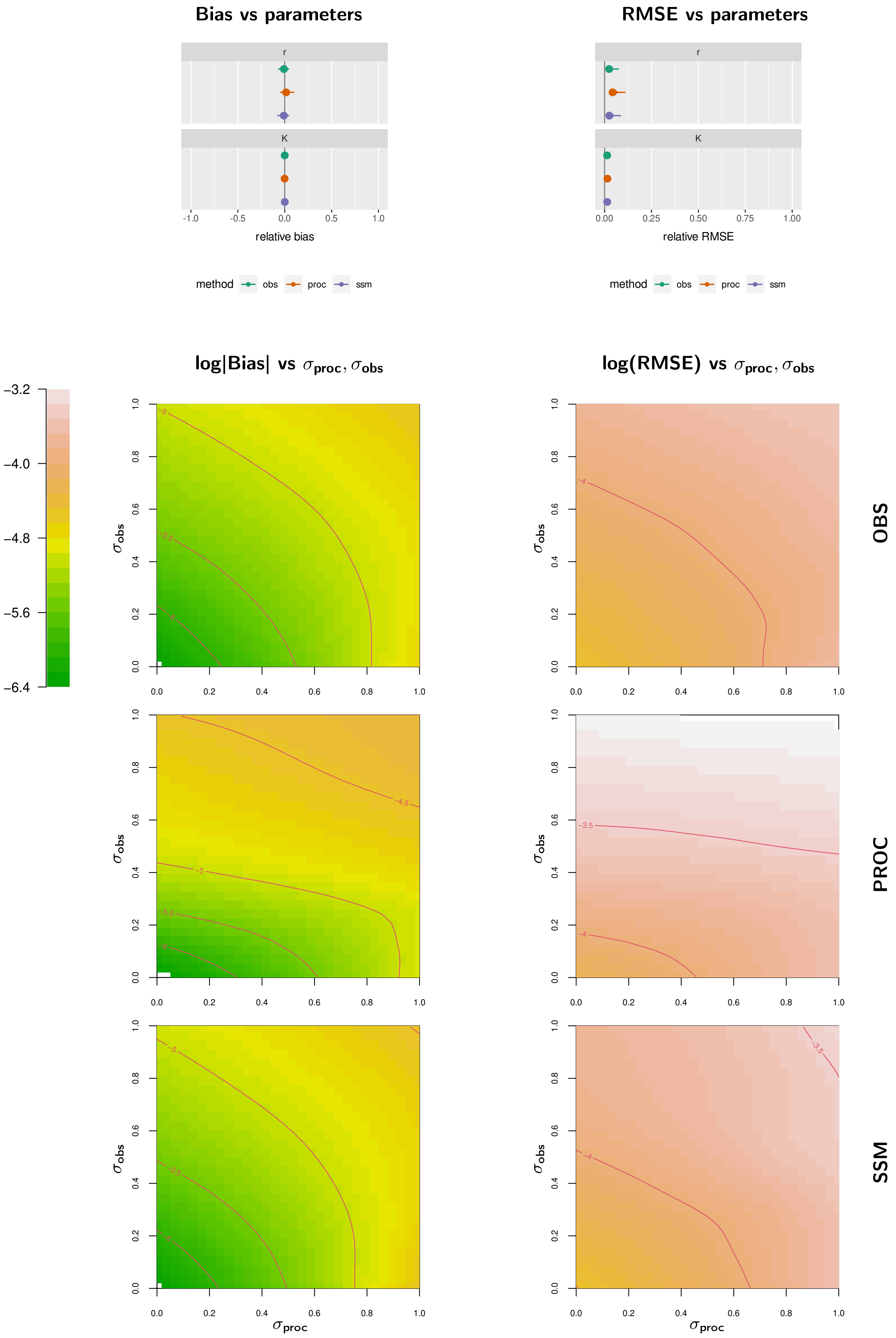


Figure S2: Logistic growth model. Relative Bias (left column) and relative RMSE (right column) for each parameter (1st row), statistical models separated by colors. 2d plots (GAMs) for log of mean absolute Bias (left column) and log of mean RMSE (right column) vs. $(\sigma_{\text{proc}}, \sigma_{\text{obs}})$, statistical models separated by row (2nd to 4th row).

### Beverton-Holt

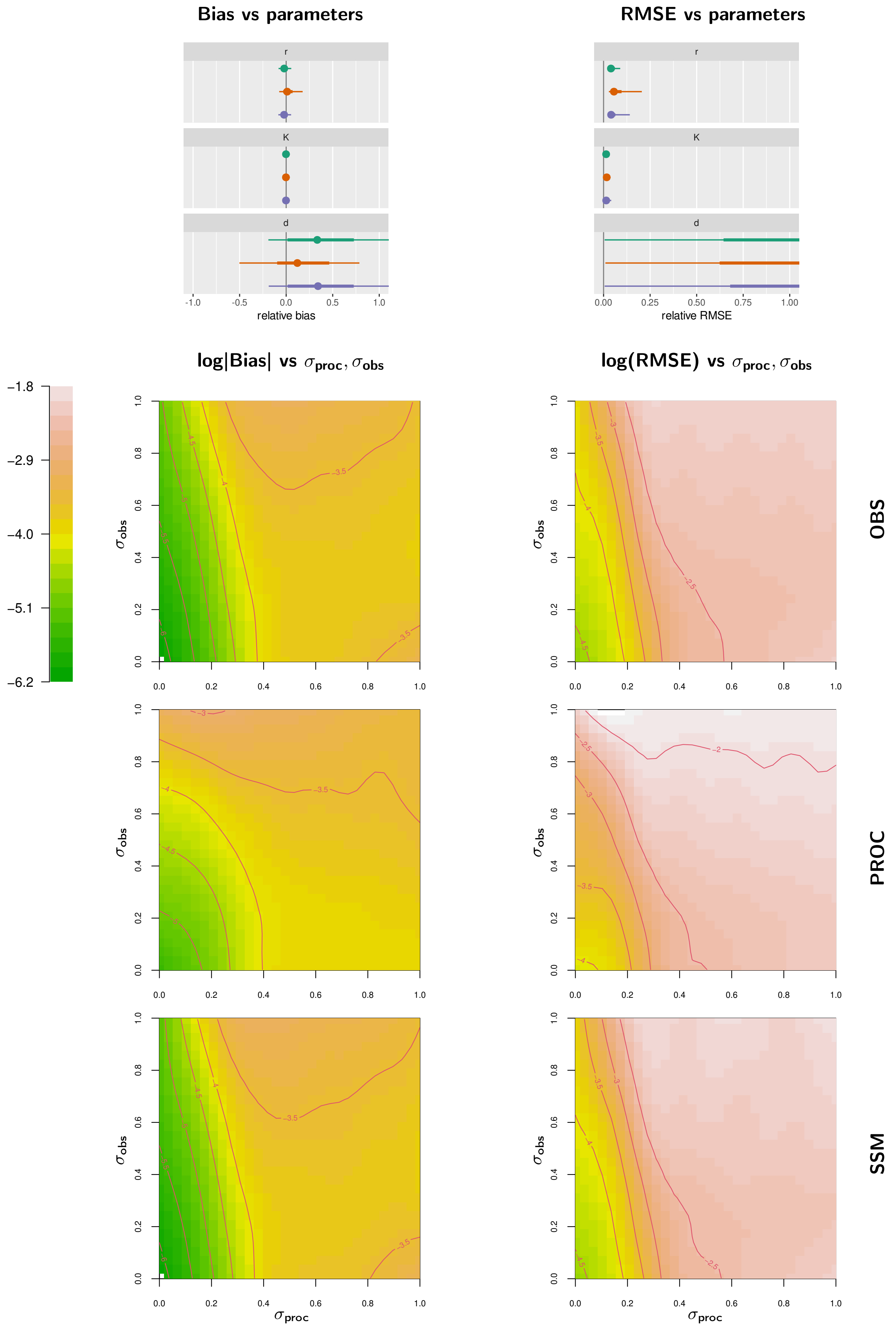


Figure S3: Beverton-Holt model. Organisation as in Fig. S2.

#### logistic Allee

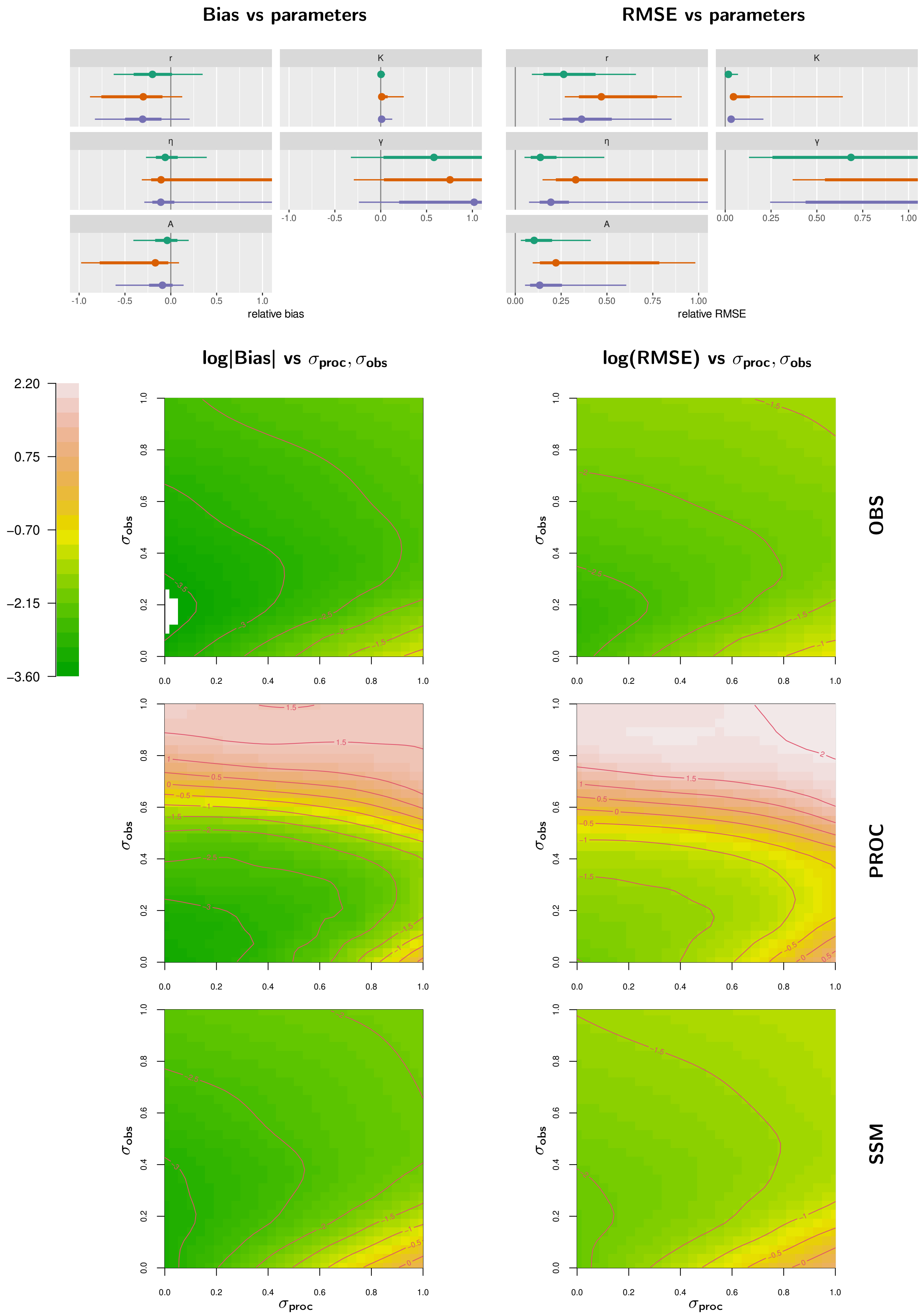


Figure S4: Logistic Allee model. Organisation as in Fig. S2.

### Beverton-Holt Allee

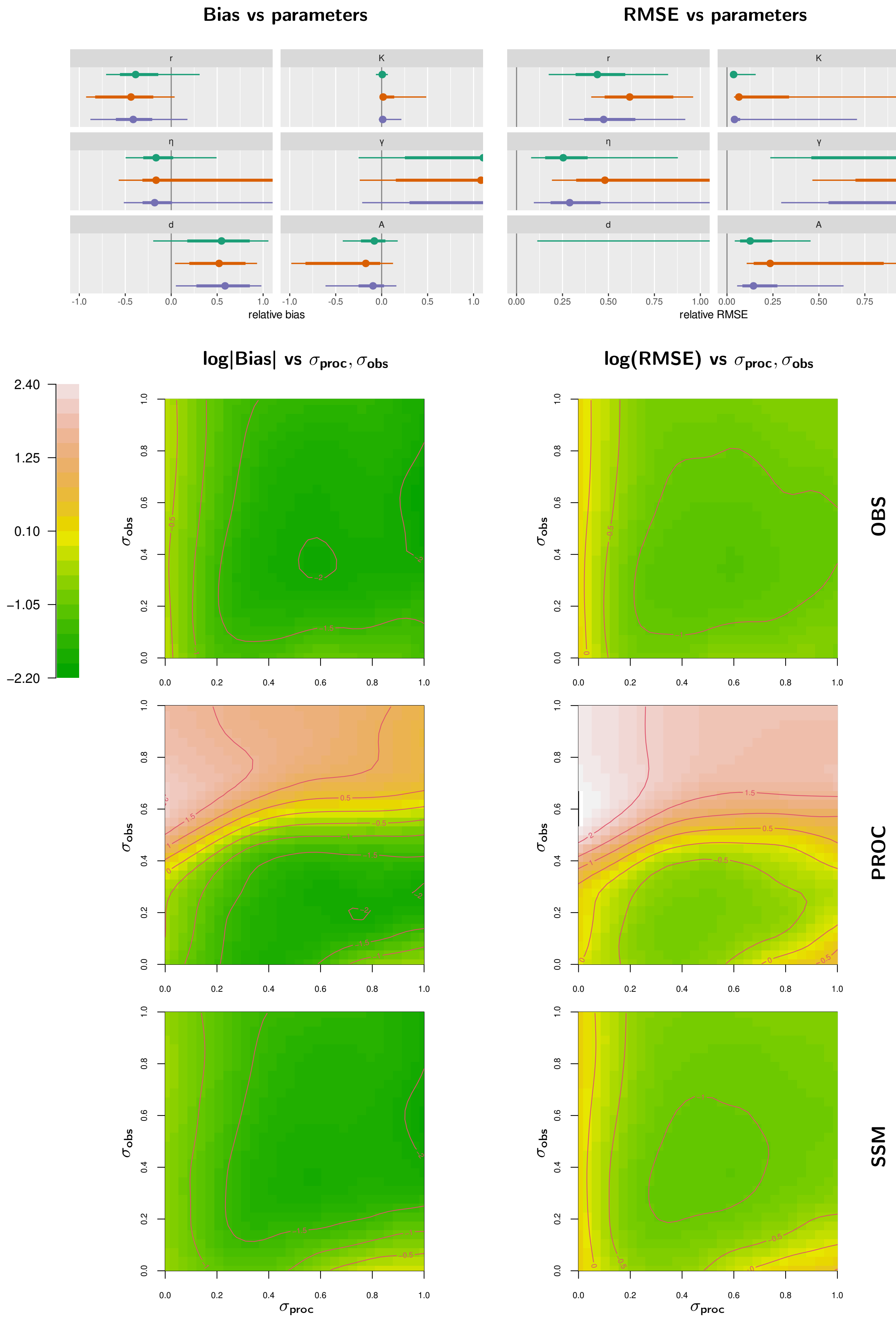


Figure S5: Beverton-Holt Allee model. Organisation as in Fig. S2.

#### Lotka-Volterra competition (coexistence)

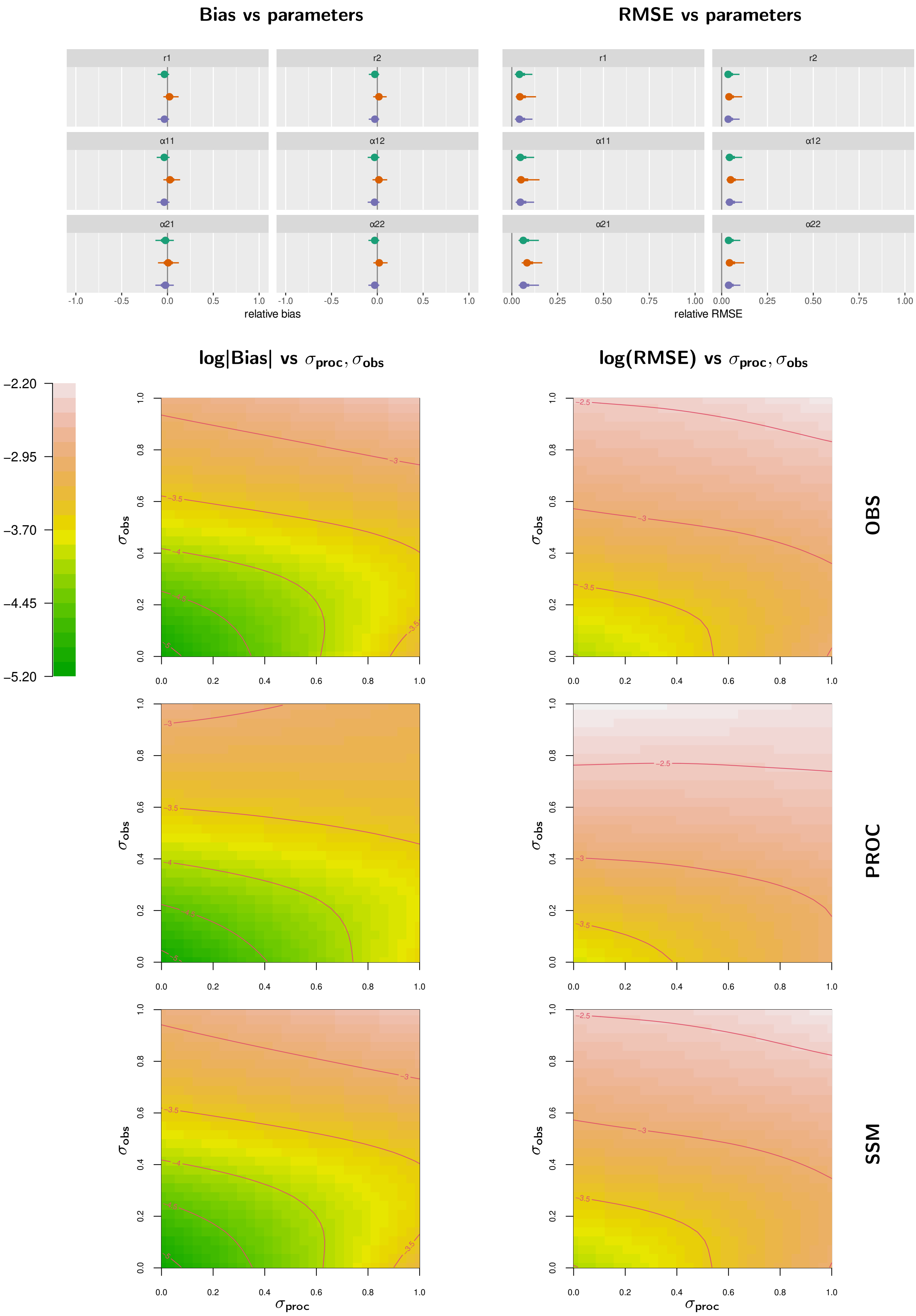


Figure S6: Lotka-Volterra competition model (coexistence). Organisation as in Fig. S2.

### Lotka-Volterra competition (exclusion)

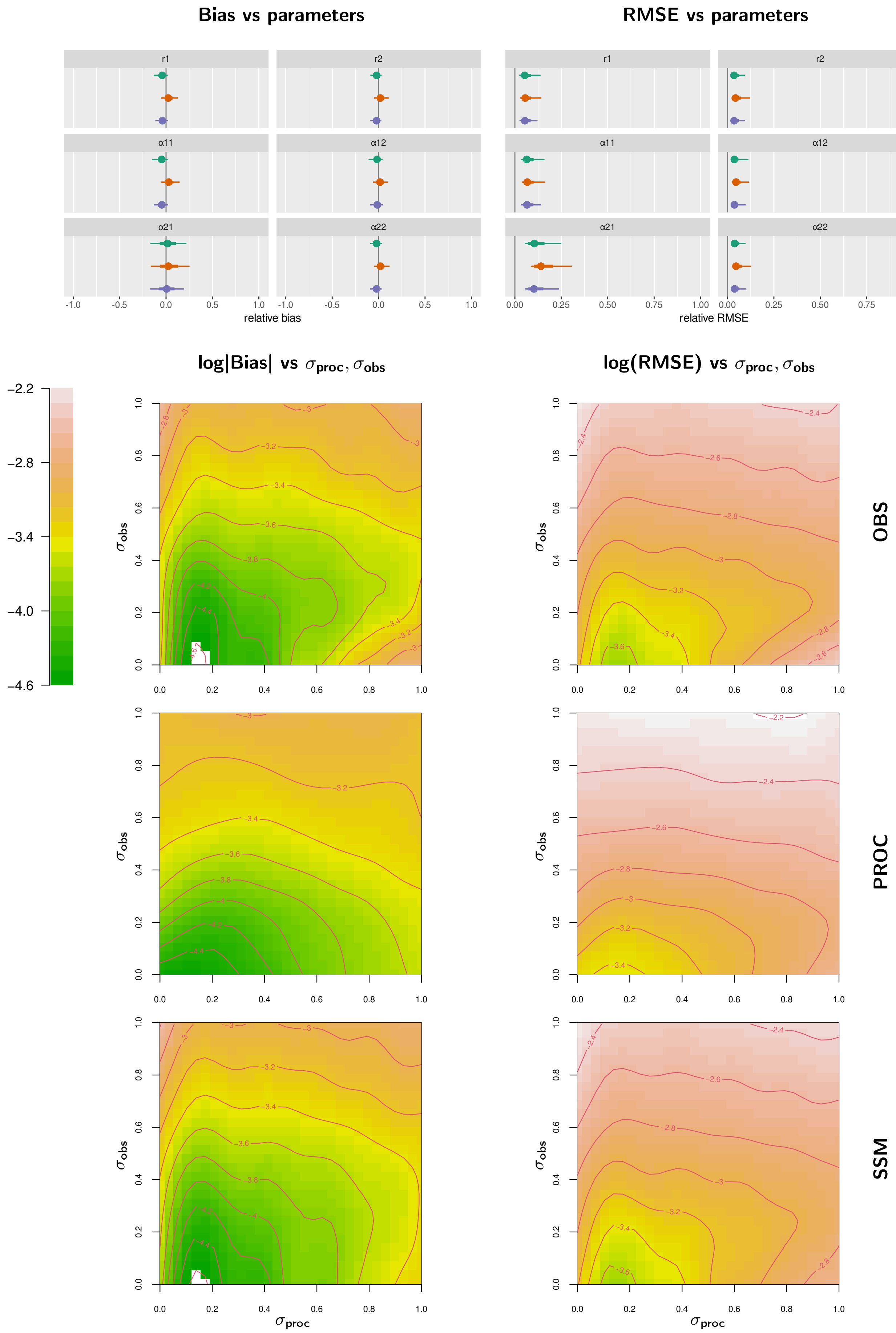


Figure S7: Lotka-Volterra competition model (exclusion). Organisation as in Fig. S2.

### Predator-prey

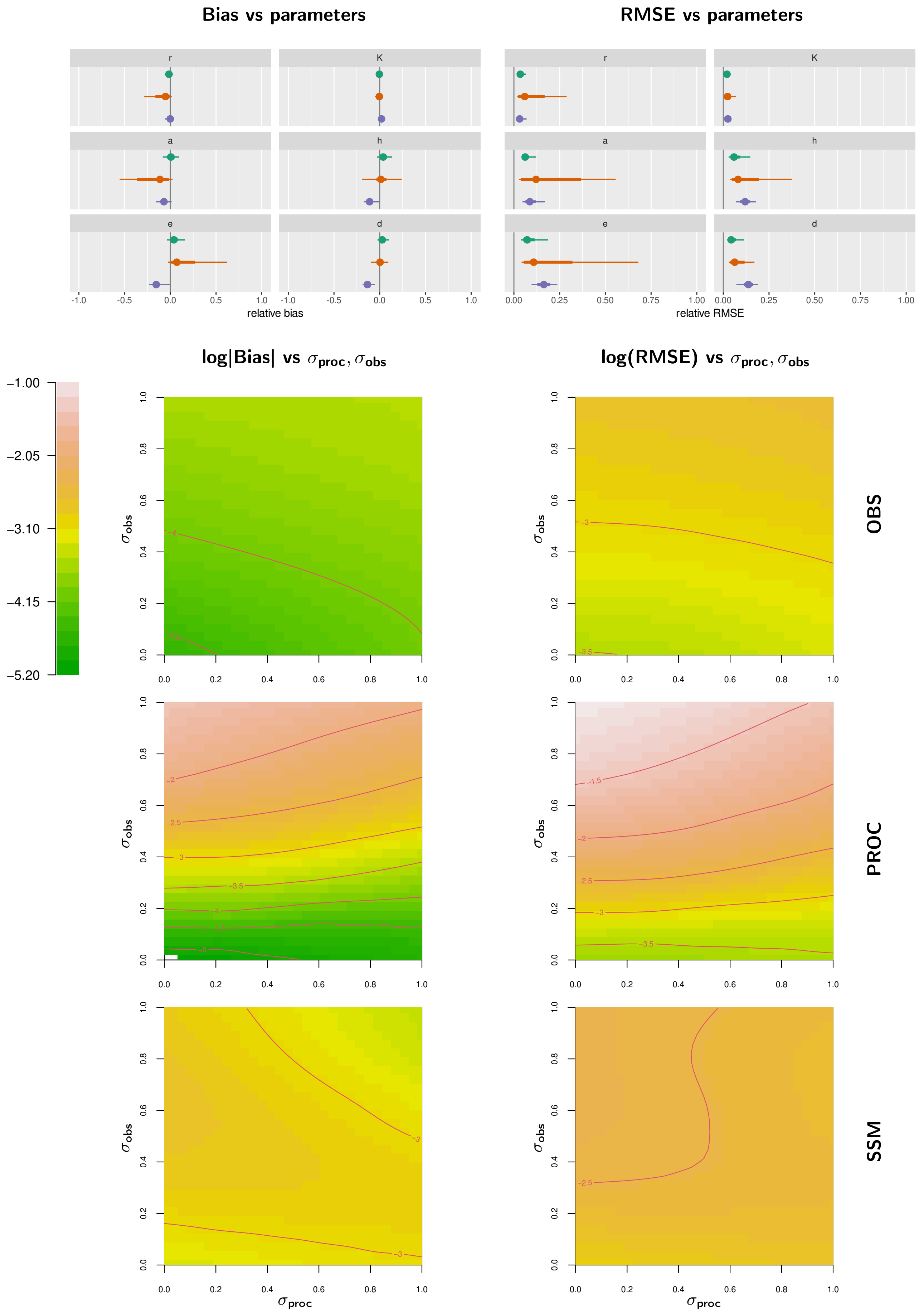


Figure S8: Predator-prey model. Organisation as in Fig. S2.

### Lotka-Volterra competition (coexistence) without control data

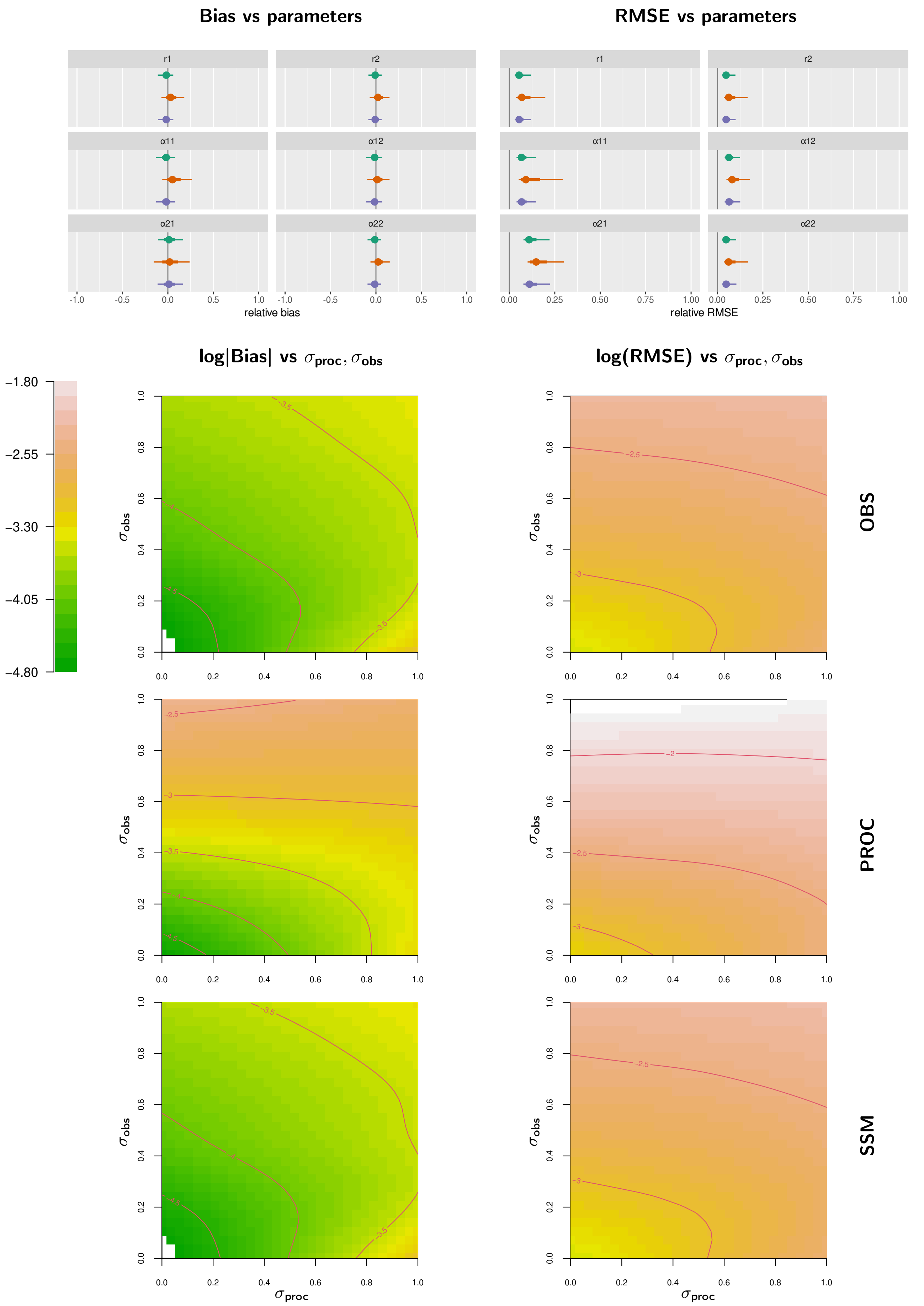


Figure S9: Lotka-Volterra competition model (coexistence) without control data. Organisation as in Fig. S2.

### Lotka-Volterra competition (exclusion) without control data

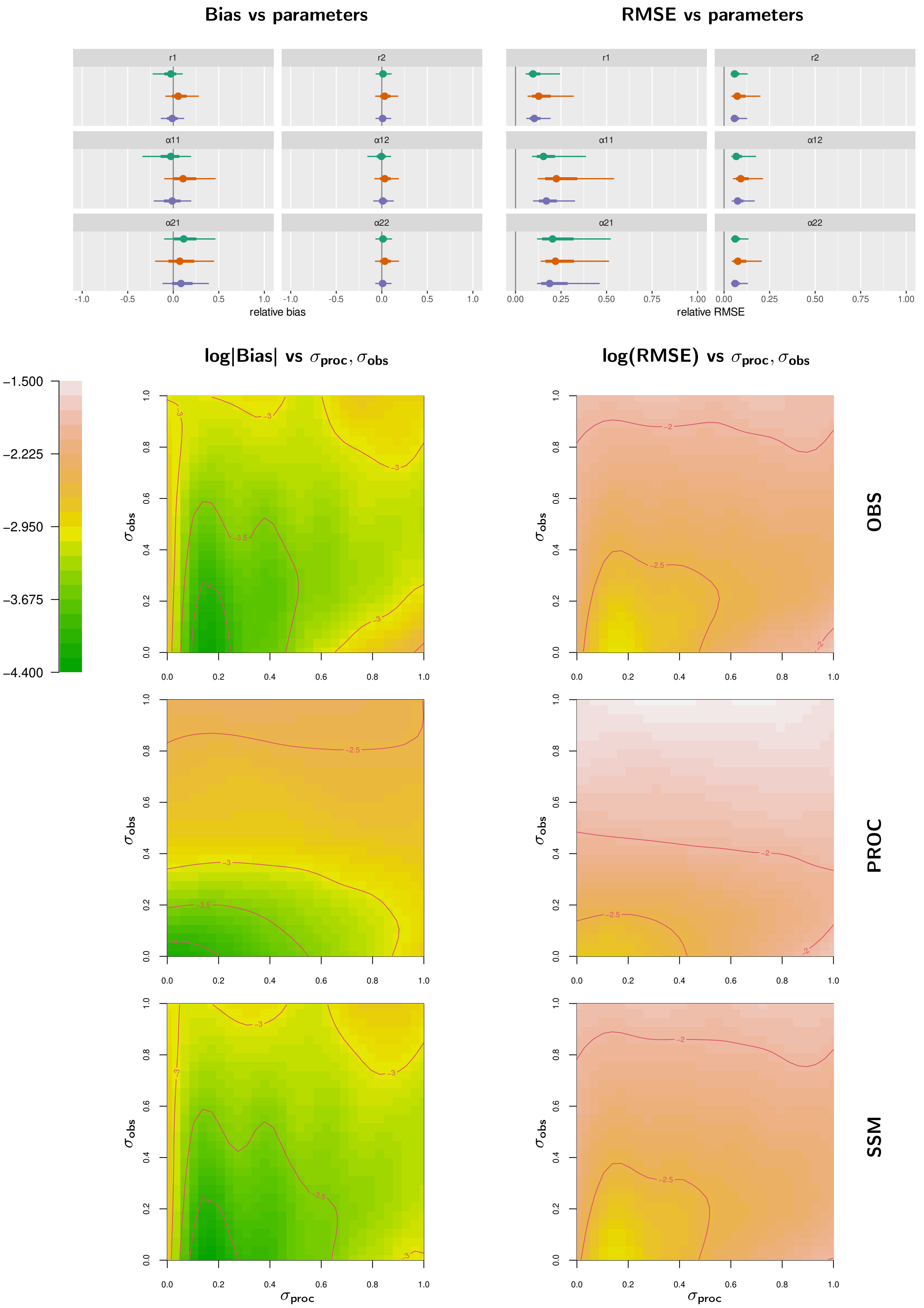


Figure S10: Lotka-Volterra competition model (exclusion) without control data. Organisation as in Fig. S2.

### Predator-prey without control data

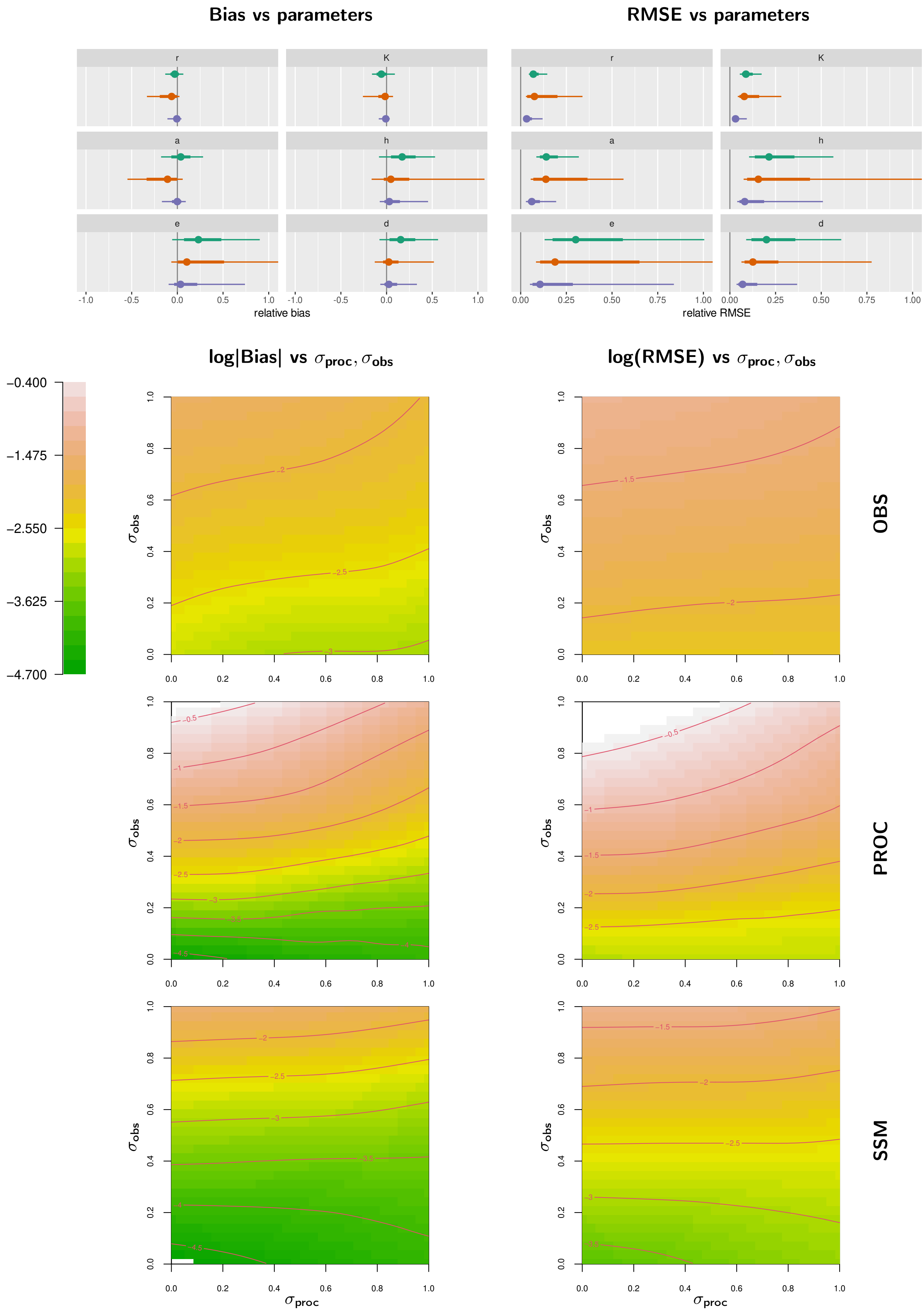


Figure S11: Predator-prey model without control data. Organisation as in Fig. S2.

#### Lotka-Volterra competition (coexistence), fewer replicates (two)

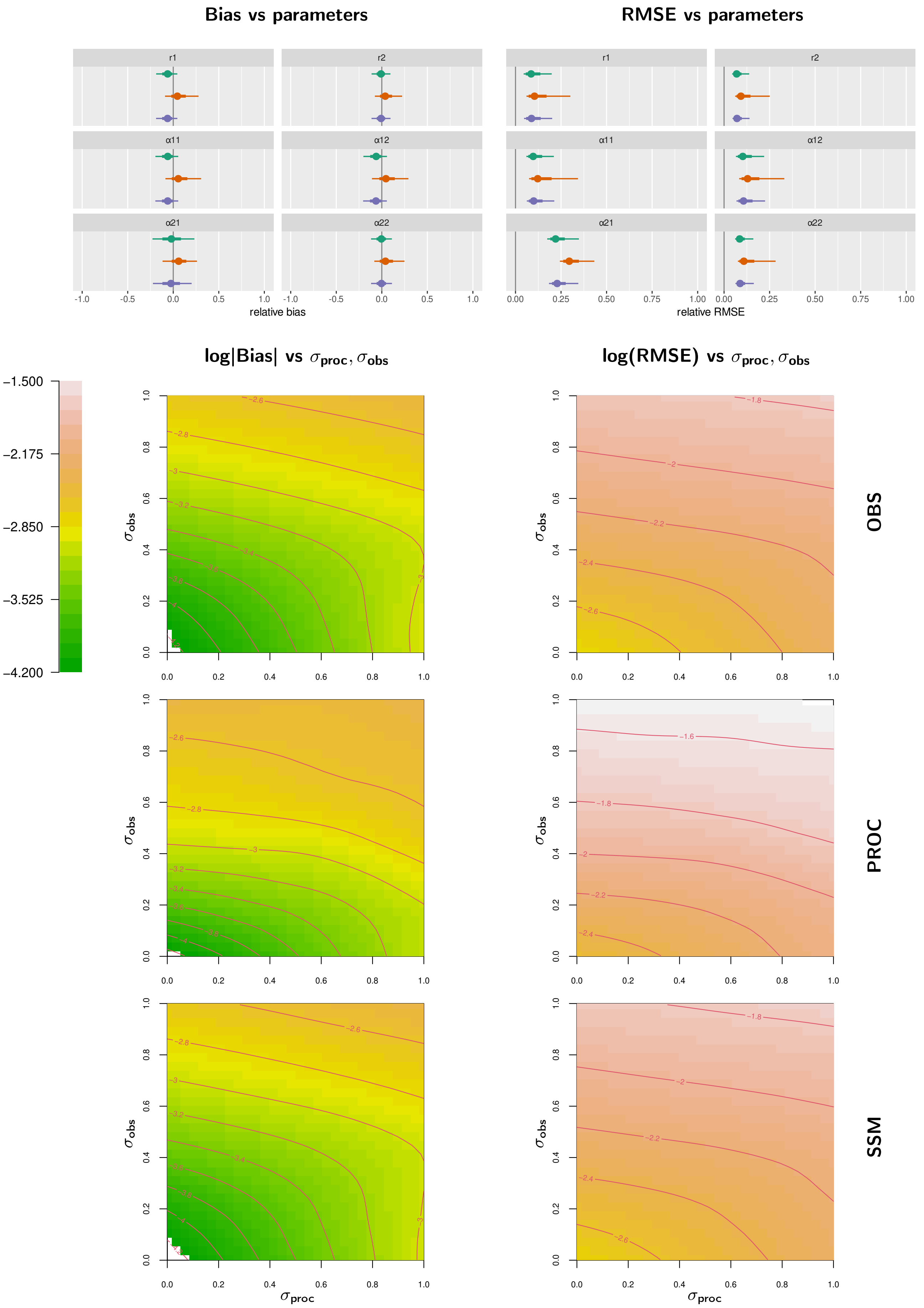


Figure S12: Lotka-Volterra competition model (coexistence), fewer replicates (two). Organisation as in Fig. S2.

### Predator-prey, longer time series

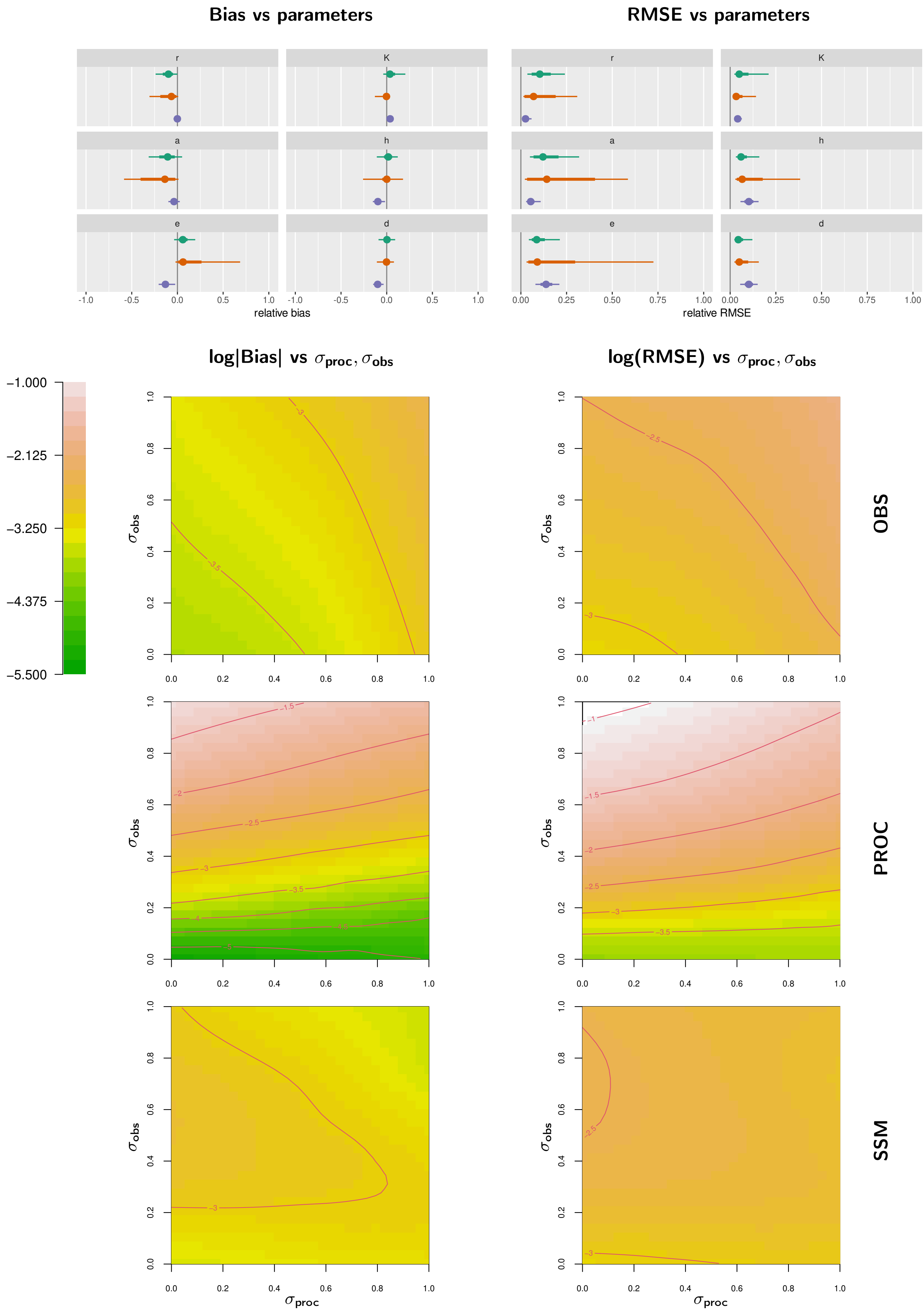


Figure S13: Predator-prey model, longer time series. Organisation as in Fig. S2.

### Logistic growth, lower carrying capacity $K$=1000

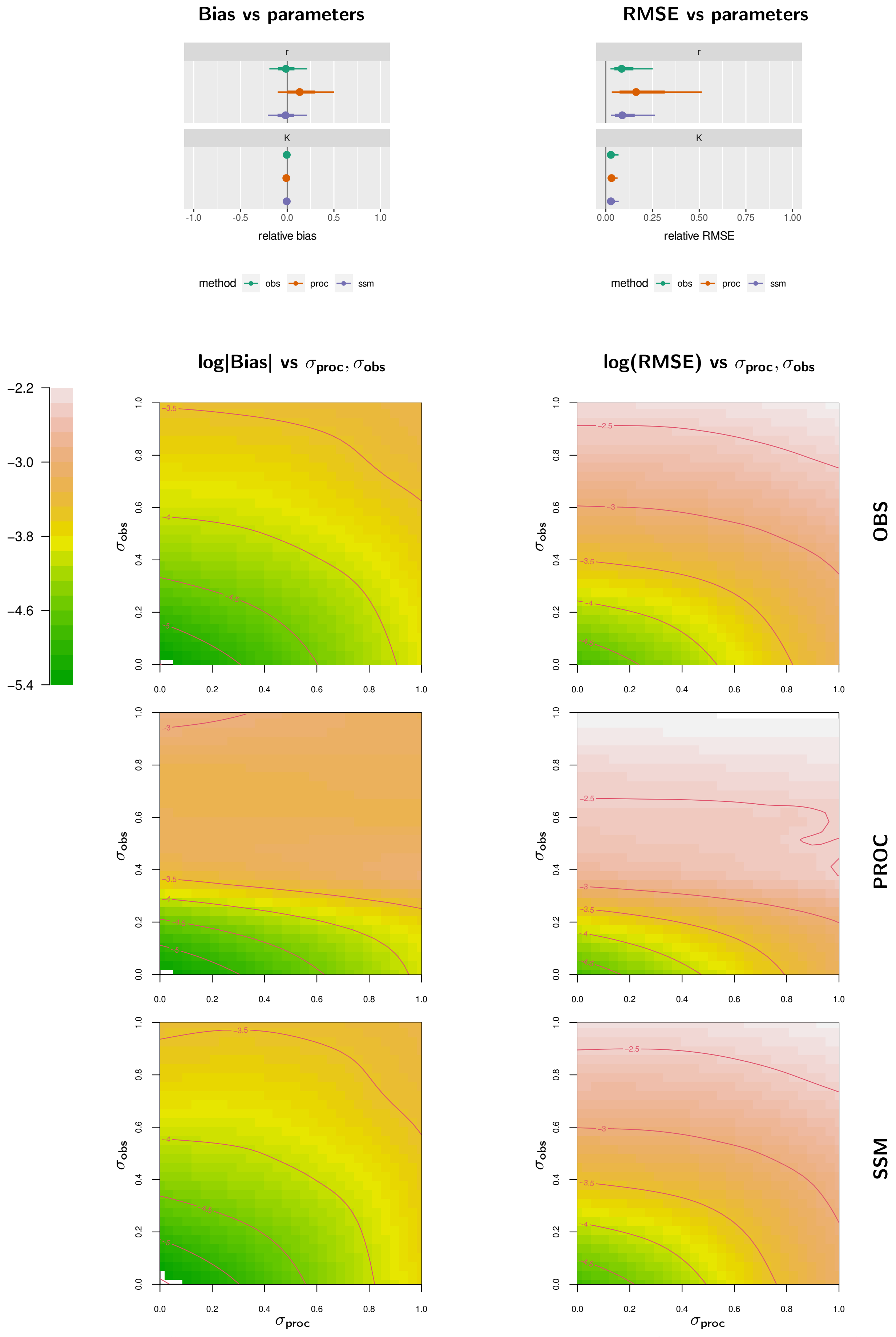


Figure S14: Logistic growth, lower carrying capacity $K = 1000$. Organisation as in Fig. S2.

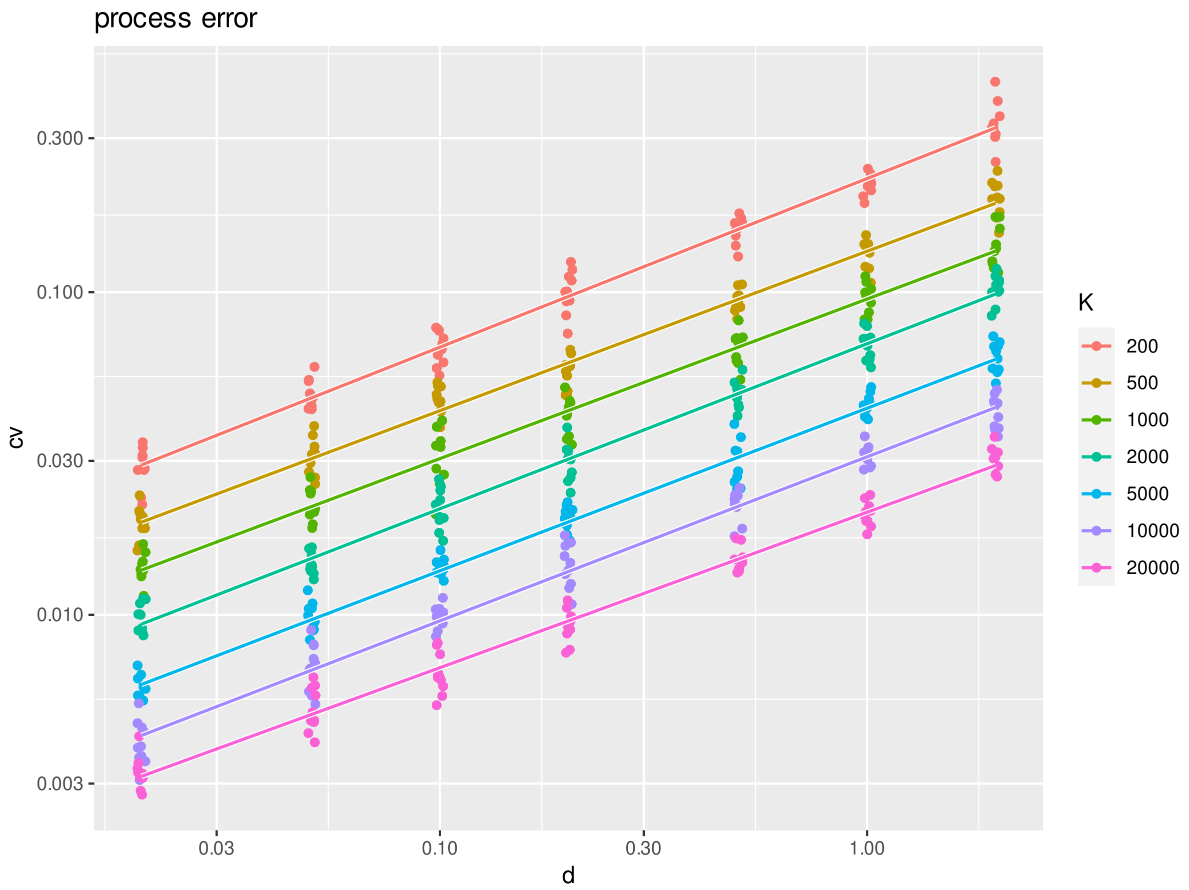


Figure S15: Process error quantification for the logistic growth stochastic simulations. Statistical linear model "$\log(\sigma_{\text{proc}}) \sim \log(d) + \log(K)$" on logscale reveals slopes $\beta_d = 0.502$, $\beta_K = 0.499$ ($n = 490$, $P < 0.001$), indicating a scaling of $\sigma_{\text{proc}} \sim \sqrt{d \cdot K}$.

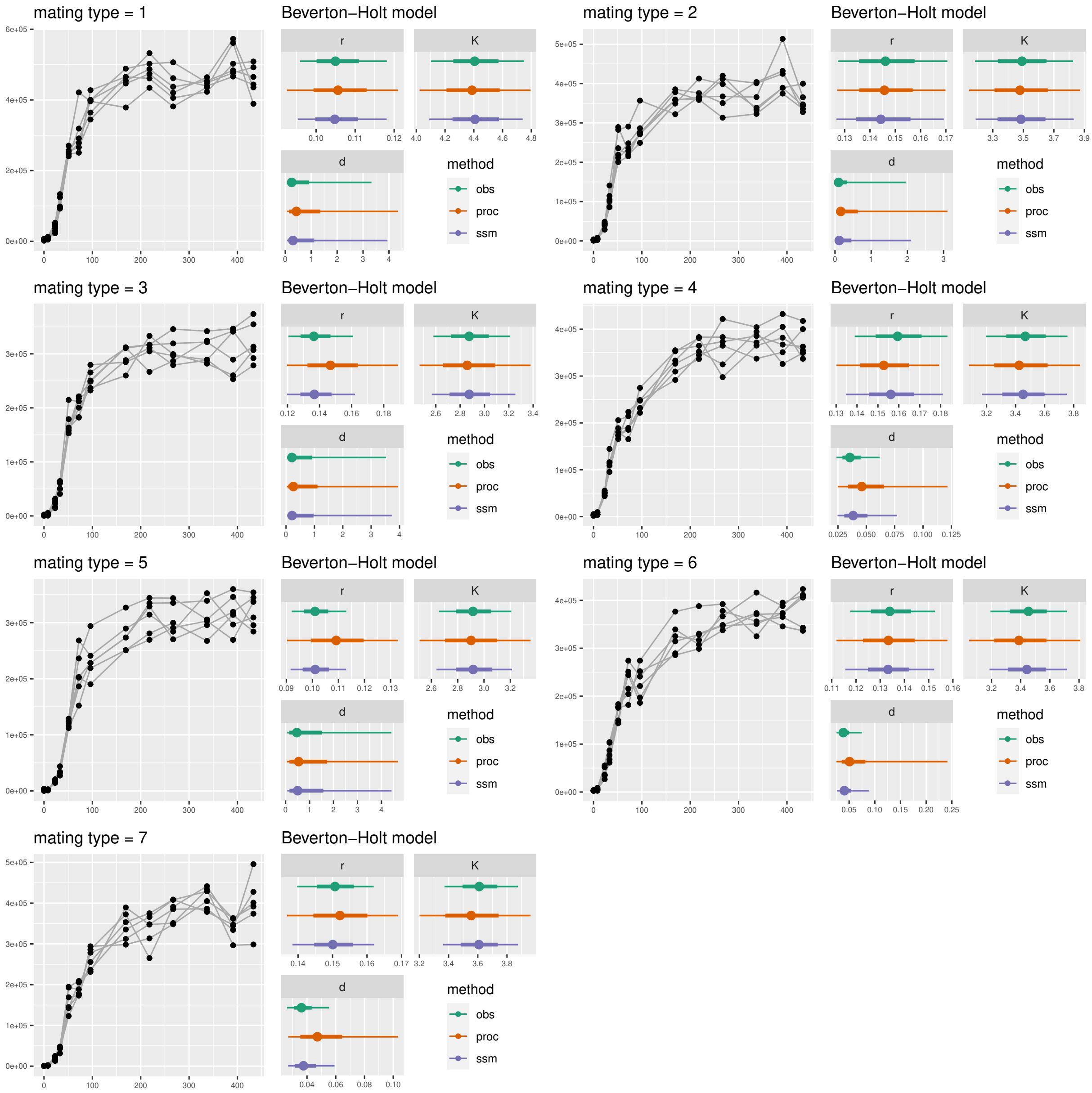


Figure S16: All seven empirical datasets and posterior distributions of fitted Beverton-Holt models, including model parameters $r, K, d$.

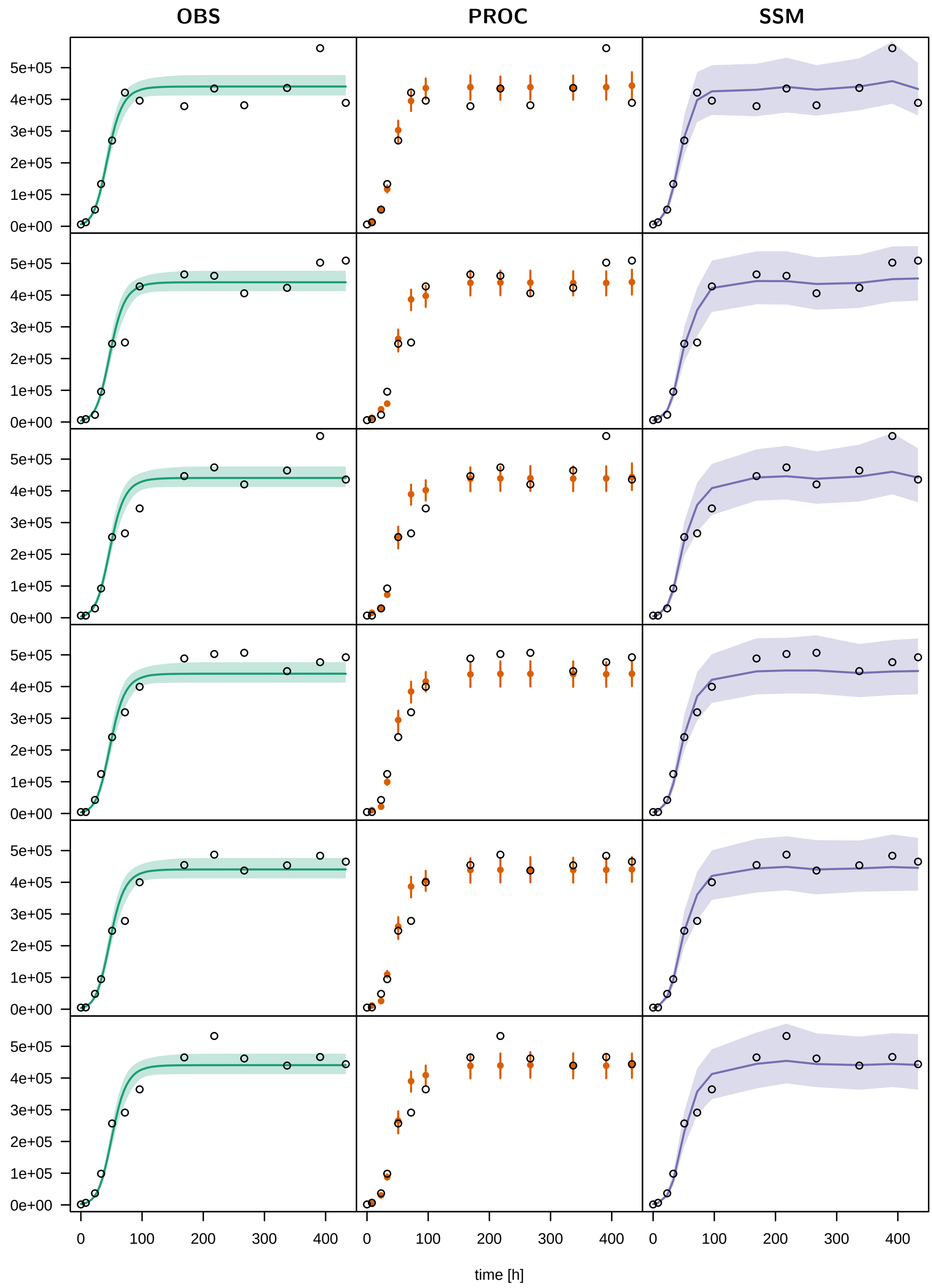


Figure S17: Posterior prediction plots for empirical dataset 1. Each row displays a different replicate (1–6) of this dataset, each column represents a different statistical model (OBS, PROC, SSM). OBS: Median and 95% credible intervals for the fitted trajectory are shown. PROC: Median and 95% credible intervals for the one-step-ahead prediction are shown. SSM: Median and 95% credible intervals for the estimated underlying true states $Z_i$ are shown (linear interpolation between timesteps). See the attached manual for code and additional description of posterior prediction plots.

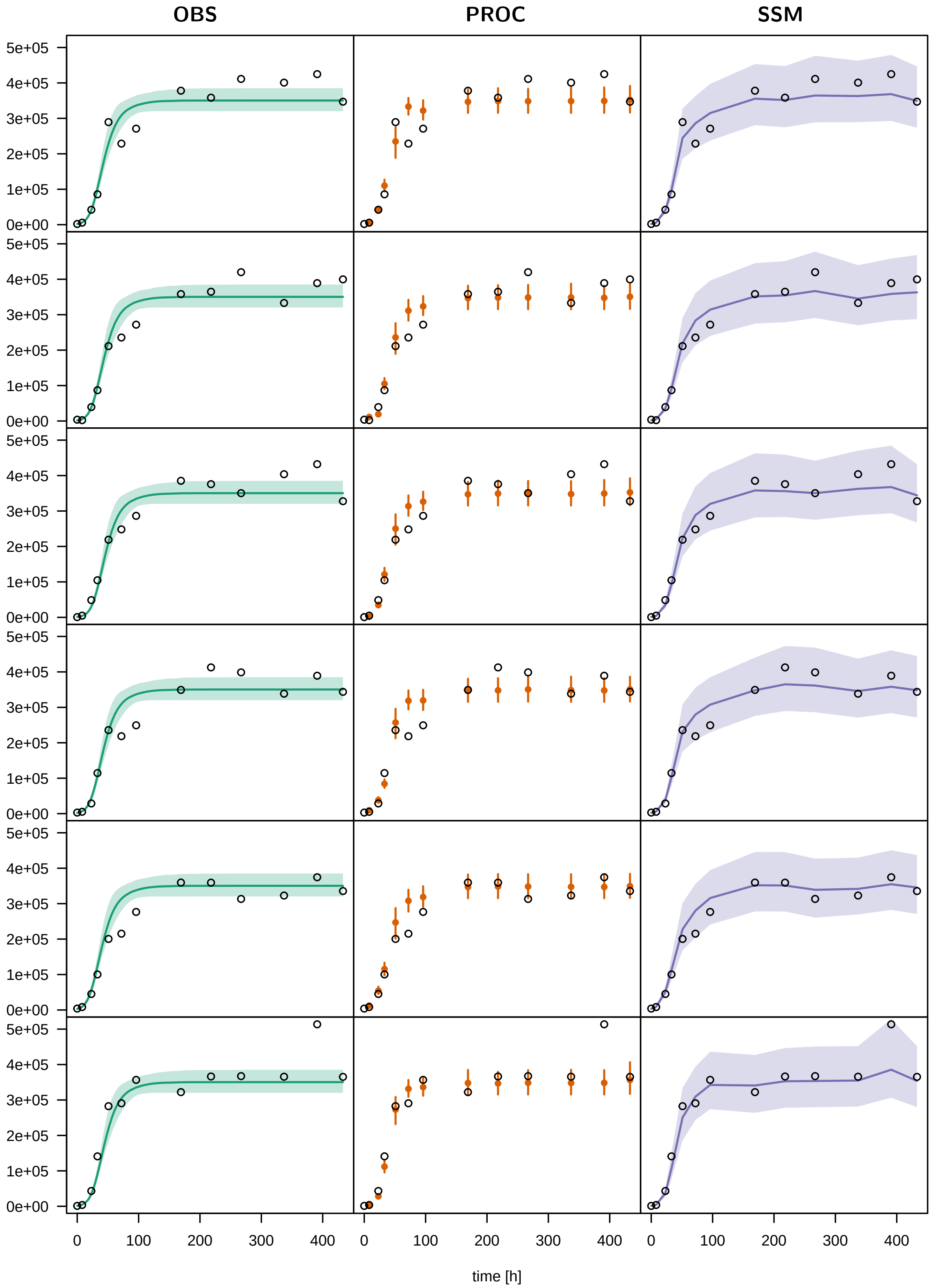


Figure S18: Posterior prediction plots for empirical dataset 2. Organisation as in Fig. S17

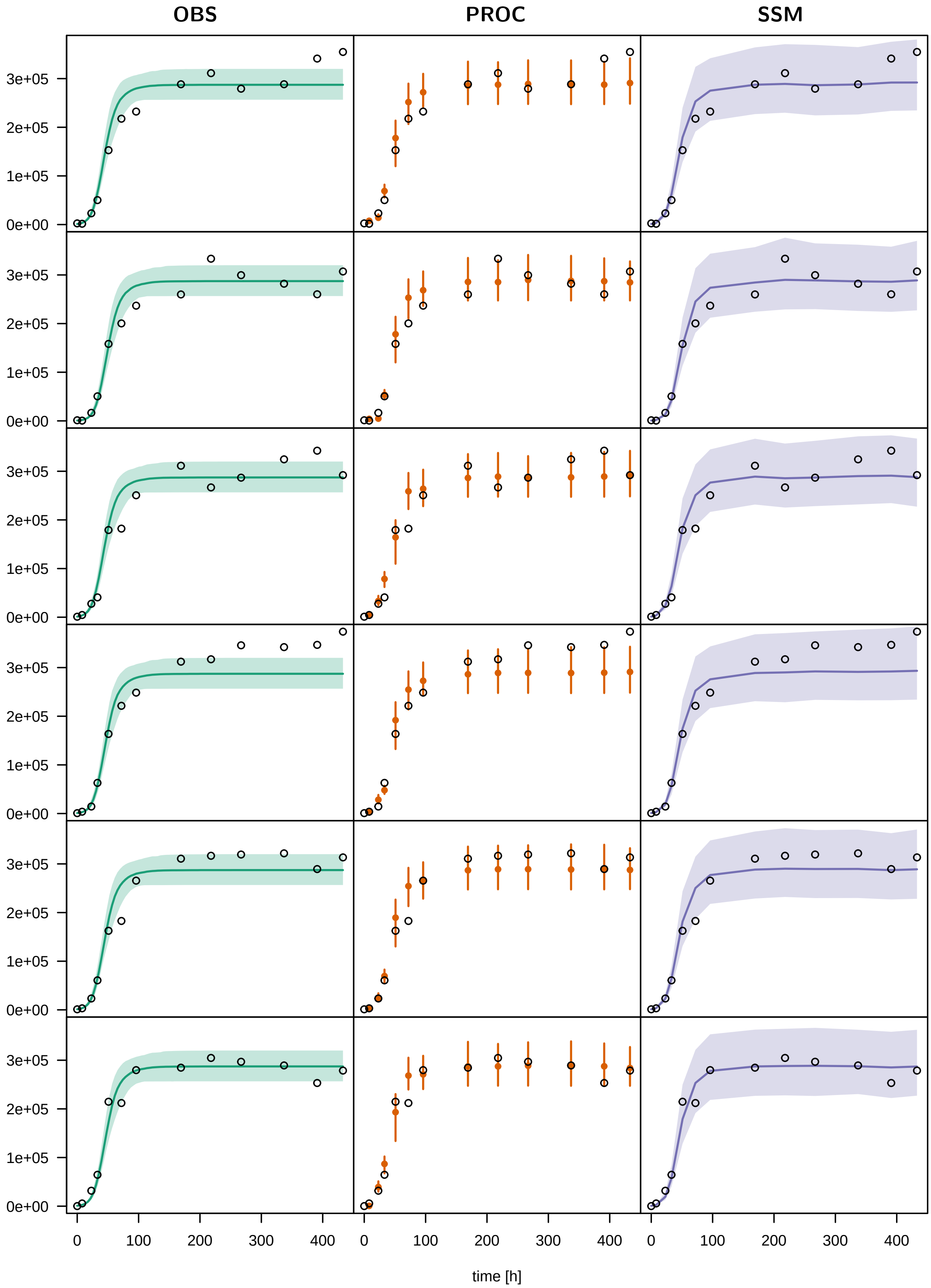


Figure S19: Posterior prediction plots for empirical dataset 3. Organisation as in Fig. S17

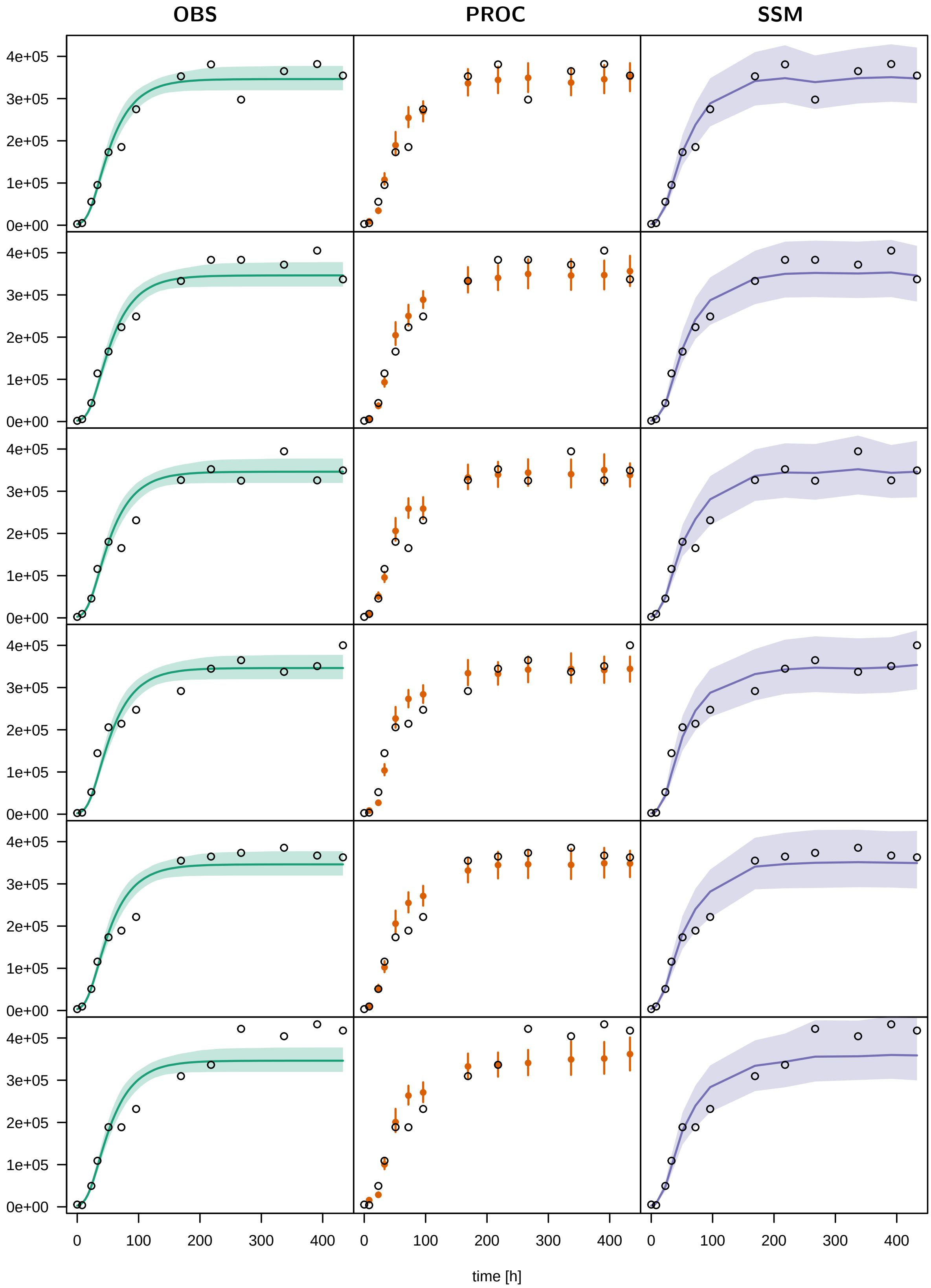


Figure S20: Posterior prediction plots for empirical dataset 4. Organisation as in Fig. S17

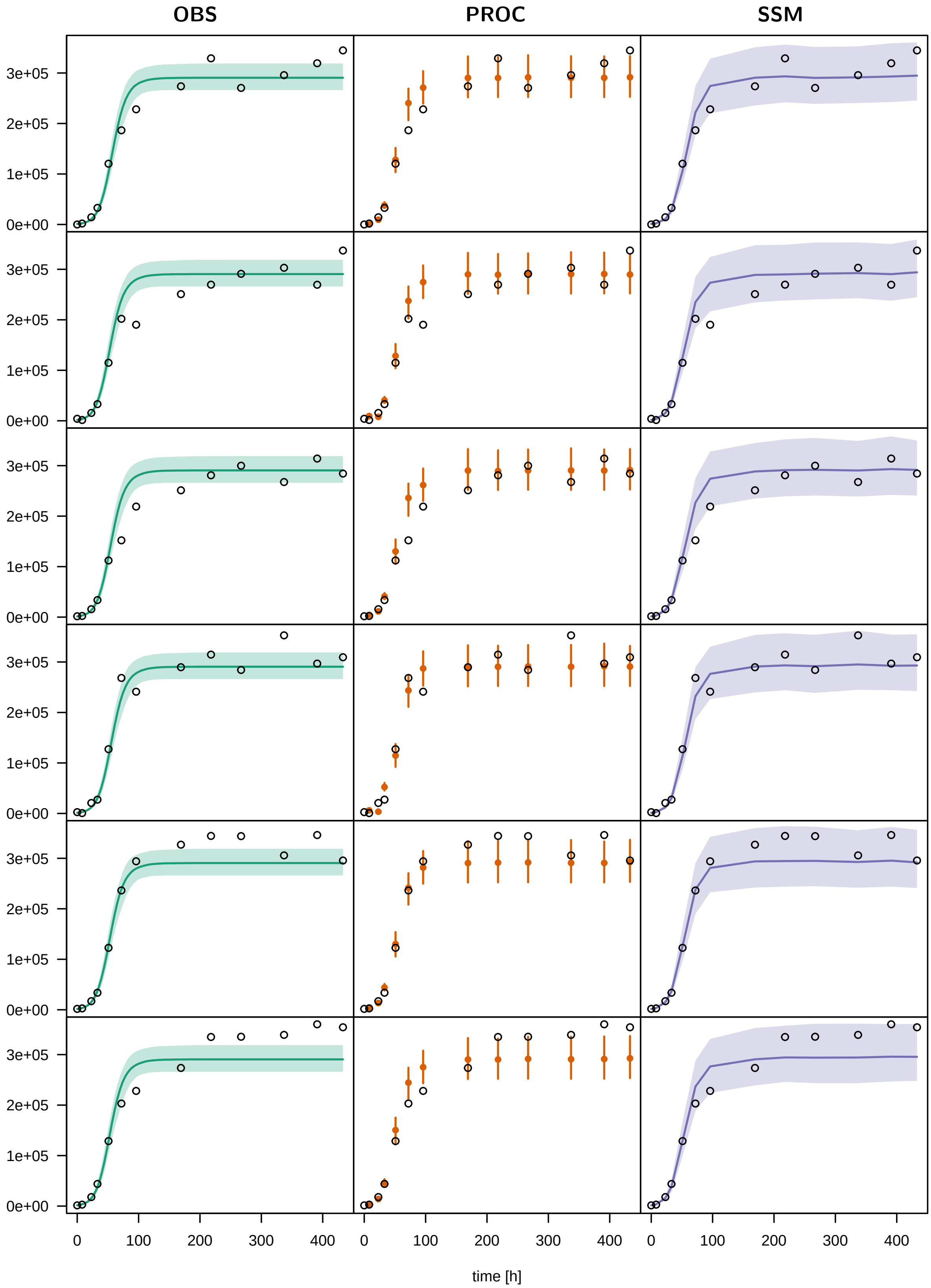


Figure S21: Posterior prediction plots for empirical dataset 5. Organisation as in Fig. S17

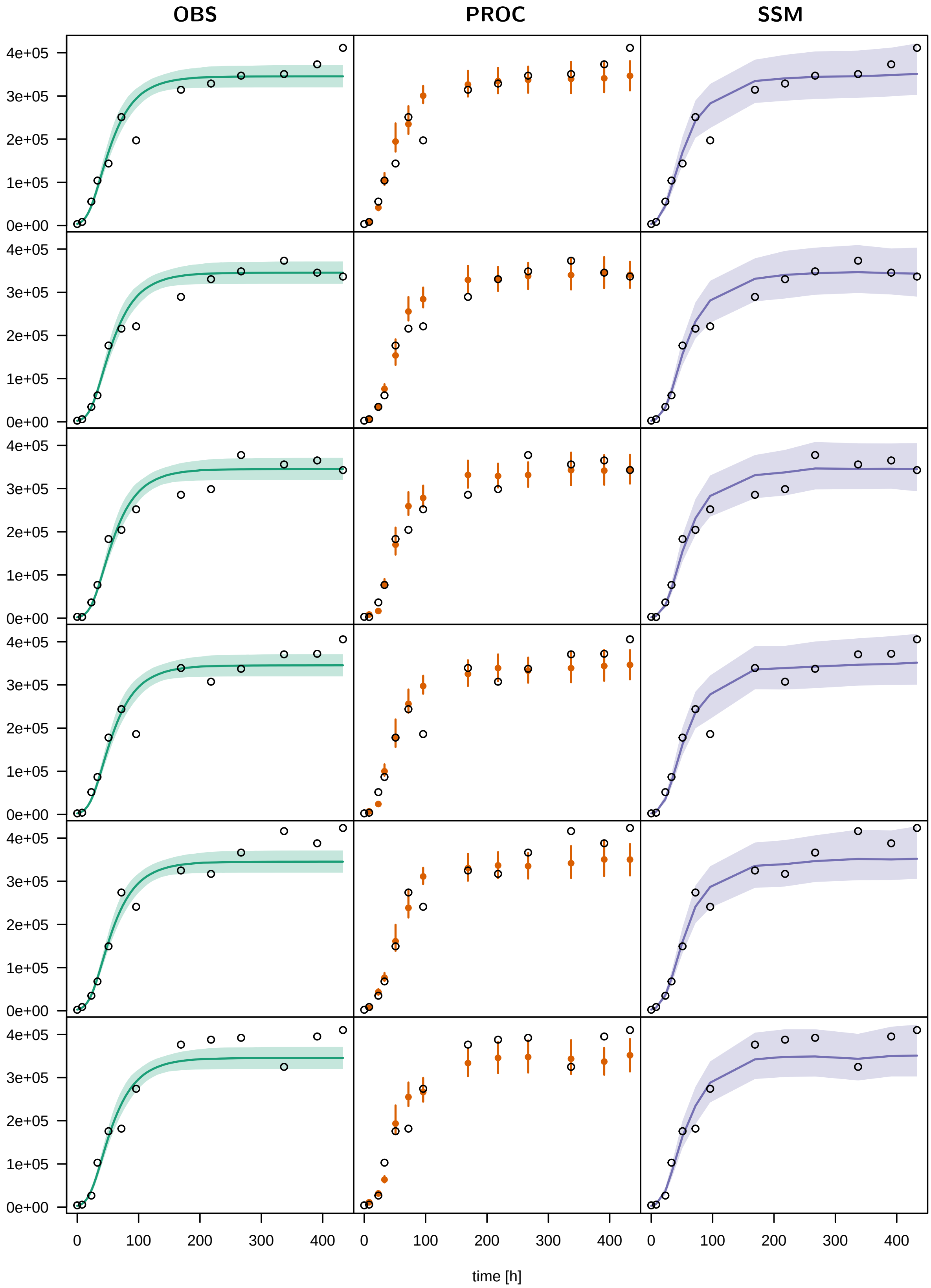


Figure S22: Posterior prediction plots for empirical dataset 6. Organisation as in Fig. S17

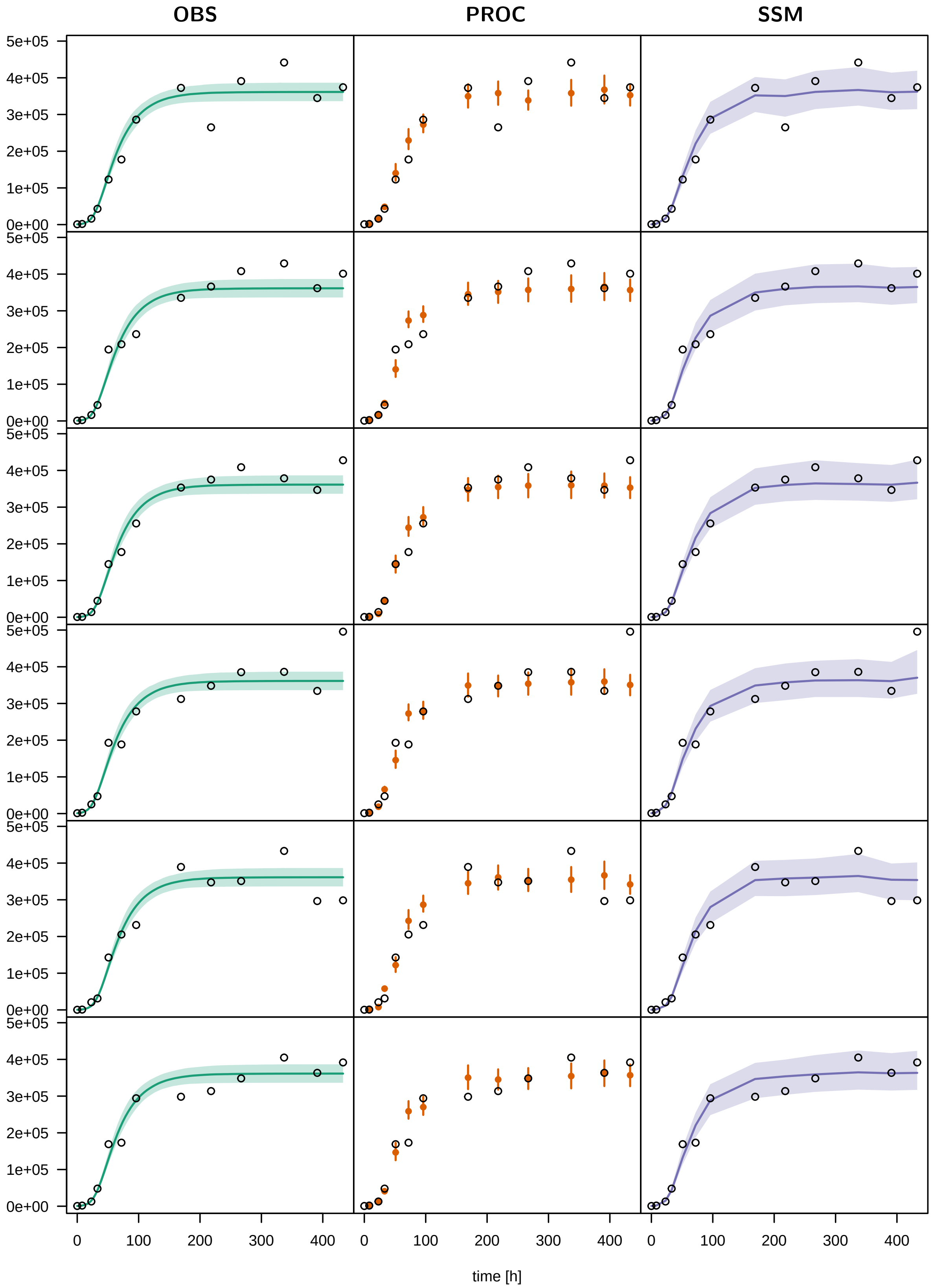


Figure S23: Posterior prediction plots for empirical dataset 7. Organisation as in Fig. S17
