## Supplementary material for "Confronting population models with experimental microcosm data: from trajectory matching to state-space models": Tutorial

#### Contents

**1 Logistic growth - OBS model** **2**

**2 Logistic growth - PROC model** **11**

**3 Logistic growth - State-space model** **19**

**4 Consumer resource - OBS model** **27**

**5 Consumer resource - PROC model** **40**

**6 Consumer resource - State-space model** **52**

Data and code for this manual can be found under:

https://github.com/benjamin-rosenbaum/fitting_deterministic_population_models

### 1 Logistic growth - OBS model

#### 1.1 Model

The logistic growth model can be written in the $r, \alpha$ or the $r, K$ formulation, where $K = \frac{r}{\alpha}$, or $\alpha = \frac{r}{K}$:

$$\begin{aligned}\frac{dN}{dt} &= (r - \alpha N)N \\ &= r(1 - \frac{N}{K})N\end{aligned}$$

Using the trajectory fitting method (observation error only), we will fit all $m$ time series replicates with a joint set of parameters $r$ and $K$ ("complete pooling"), while allowing for $m$ individual initial states $N_0 = N(t_0)$ as free parameters.

#### 1.2 Data preparation

The dataset `data_example_logistic.csv` contains `m=6` timeseries with `n=18` observations of densities [ind] each, including their dates [h]. The fraction of the observed volume for sampling was `f=0.1`.

```
rm(list=ls())
library("rstan")

data.in = read.csv("data_example_logistic.csv", row.names=1)

head(data.in)
```

```
##       X1 X2  X3  X4  X5  X6  X7   X8   X9  X10  X11  X12  X13  X14  X15  X16
## time   0 12  24  36  48  60  72   96  120  144  168  192  216  240  264  288
## N1    10 36 106 270 562 914 994  968  904 1032 1010 1004  968 1010 1048  950
## N2    24 30 136 262 532 846 942 1024 1024 1010 1016 1002 1082 1056  958 1088
## N3    10 20 146 336 660 896 938  946 1016 1044  966  994 1004 1008 1086  932
## N4    12 44 120 274 572 768 916 1010 1058 1066  992 1006  940  910 1016  998
## N5    10 20  62 142 484 656 922  934 1000 1052  988 1004 1036 1056 1020  932
##        X17  X18
## time   312  336
## N1     952 1072
## N2    1078  954
## N3    1006 1034
## N4     916  972
## N5    1062  976
```

For Stan, the data has to be coded as a named list, including dimensions of the data `n` and `m`, `times` as a vector, the `m` x `n` matrix `N` containing the observed abundances, and the sampling fraction `f`.

```
n = ncol(data.in)
m = nrow(data.in)-1

times = as.numeric(data.in[1, ])
N = unname(as.matrix(data.in[2:(m+1), ]))

data = list(n = n,
            m = m,
            t = times,
            N = N,
            f = 0.1)

str(data)
```

```
## List of 5
##  $ n: int 18
```

```
##  $ m: num 6
##  $ t: num [1:18] 0 12 24 36 48 60 72 96 120 144 ...
##  $ N: int [1:6, 1:18] 10 24 10 12 10 10 36 30 20 44 ...
##  $ f: num 0.1
```

##### 1.3 Stan model

The Stan model is coded as a string, which is compiled later. A step by step walkthrough is given below.

```
stanmodelcode = '
functions{
  real[] odemodel(real t, real[] N, real[] p, real[] x_r, int[] x_i){
    // p[1]=r, p[2]=K
    real dNdt[1];
    dNdt[1] = p[1]*(1-(N[1]/p[2]))*N[1];
    return dNdt;
  }
}

data{
  int n; // observations
  int m; // replicates
  real t[n];
  int N[m,n];
  real f;
}

parameters{
  real<lower=0> r;
  real<lower=0> K;
  real<lower=0> tau;
  real<lower=0> N0sim[m]; // individual initial values for replicates
}

model{
  real p[2];
  real Nsim[n-1,1]; // simulated values, matrix. dim1 = time without t0, dim2 = dim_ODE = 1

  // priors
  r ~ lognormal(-2,1);
  K ~ lognormal(9,1);
  N0sim ~ normal(0,100);
  tau ~ gamma(2,0.1);

  // parameters for integrator
  p[1] = r;
  p[2] = K;

  for (j in 1:m){
    // integrate ODE
    Nsim = integrate_ode_rk45(odemodel, {N0sim[j]}, t[1], t[2:n], p, rep_array(0.0,0), rep_array(0,0
    // likelihood
    N[j,1] ~ neg_binomial_2(N0sim[j]*f, tau);
    for (i in 2:n){
      N[j,i] ~ neg_binomial_2(Nsim[i-1,1]*f, tau);
    }
  }
}
```

```
generated quantities{
  real alpha = r/K;
}
'
```

##### 1.3.1 functions{ } block

The population growth equation (the ODE) is coded as a function. This function is later used for numerical integration. Although the logistic growth ODE has a closed-form analytical solution, we still use numerical solutions here, such that users can exchange the ODE for other models easily.

Stan requires a specific format: The output is a real-valued array, and arguments time `t`, state `N`, and model parameters `p` are provided. Additional arguments `x_r` and `x_i` for real- and integer-valued data have to be defined, but are not used here.

The population growth rate `dNdt` is computed using the state `N` and parameters `p` (including `r` and `K`) and returned as the function's output.

```
real[] odemodel(real t, real[] N, real[] p, real[] x_r, int[] x_i){
  // p[1]=r, p[2]=K
  real dNdt[1];
  dNdt[1] = p[1]*(1-(N[1]/p[2]))*N[1];
  return dNdt;
}
```

##### 1.3.2 data{ } block

All data have to be declared including data types and dimensions. Names have to be identical to the named list which was defined in R before.

```
int n; // observations
int m; // replicates
real t[n];
int N[m,n];
real f;
```

##### 1.3.3 parameters{ } block

Here, all free parameters are declared. This includes model parameters `r` and `K`, and a parameter `tau` for the residual distribution (overdispersion of the negative binomial). Additionally, the initial states of each of the `m` modelled time series `N0sim[m]` in the total volume are estimated, too (defined as a real-valued array). Its true states are unknown and observations are assumed to be measured with error. All parameters are required to be positive.

```
real<lower=0> r;
real<lower=0> K;
real<lower=0> tau;
real<lower=0> N0sim[m]; // individual initial values for replicates
```

##### 1.3.4 model{ } block

This block defines how the posterior computed, using prior distributions and the likelihood function. First, some intermediate variables for the ODE simulation are declared. `p` is an array which stores model parameters `r` and `K`. `Nsim` will contain the output of the numerical integration with dimensions `n-1` (at all timepoints minus the first one), and the second dimension is number of states (here just 1).

```
real p[2];
real Nsim[n-1,1]; // simulated values, matrix. dim1 = time without t0, dim2 = dim_ODE = 1
```

Then, priors are defined for the model parameters. Weakly informative priors are chosen for both model parameters `r` and `K`. The overdispersion parameter `tau` of the residuals is constrained >0, but arbitrary small values are possible. We choose a gamma prior that decreases towards zero, such that sampling at low values near the boundary doesn't cause convergence issues. The parameters for the initial values `N0sim` in the total volume are assigned weak priors, too.

```
r ~ lognormal(-2,1);
K ~ lognormal(9,1);
N0sim ~ normal(0,100);
tau ~ gamma(2,0.1);
```

After the priors are defined, predictions are computed. This includes numerically integrating the ODE, which requires all model parameters are stored in one array `p`.

```
p[1] = r;
p[2] = K;
```

In a loop over all `m` time series replicates, numerical integration is performed. Each time series `j` gets its individual initial state `N0sim[j]`. The integration routine `integrate_ode_rk45()` is must be called in a specific format. Arguments are the function `odemodel` as defined in the `functions{}` block, the initial state is converted to an array using `{ }`, then initial time `t[1]` and output times `t[2:n]`, and model parameters `p`. For additional variables `x_r` and `x_i` (mandatory but not used here), empty arrays are generated.

After computing the predicted time series `Nsim`, it is confronted with the data `N`: a negative binomial distribution defines likelihood values. Since the data was sampled in a fraction `f=0.1`, the distribution's mean is `Nsim[,]*f`, with some overdispersion `tau`.

Since predictions by `integrate_ode_rk45()` can only be generated for $t > t_1$, data `N[j,1]` and `N[j,i]` ($i > 1$) have to be treated separately. The first observation `N[j,1]` is distributed around the prediction in $t_1$, which is equal to the free parameter `N0sim[j]*f` (estimated initial state). The remaining observations `N[j,i]` ($i > 1$) are distributed around the predictions in $t_i$, respectively, which are equal to the output `Nsim[i-1,1]*f`. It is `i-1` instead of `i` here, because the output `Nsim` starts in $t_2$ and not in $t_1$.

```
for (j in 1:m){
  // integrate ODE
  Nsim = integrate_ode_rk45(odemodel, {N0sim[j]}, t[1], t[2:n], p, rep_array(0.0,0), rep_array(0,0
  // likelihood
  N[j,1] ~ neg_binomial_2(N0sim[j]*f, tau);
  for (i in 2:n){
    N[j,i] ~ neg_binomial_2(Nsim[i-1,1]*f, tau);
  }
}
```

##### 1.3.5 generated quantities{ } block

Optionally, other quantities can be defined in this block. Samples of their posterior distributions are saved, too. E.g., we can save the term $\alpha = \frac{r}{K}$' of the logistic growth model's $r$,$\alpha$-formulation.

```
real alpha = r/K;
```

#### 1.4 Model fitting

After defining the model code, we specify some parameters for the MCMC model fitting. Usually, 3–5 MCMC chains are used. These can be run in parallel on multicore processors, when options are specified accordingly. Further, we set the total number of iterations and number of warmup iterations for the sampling. The total number of posterior MCMC samples is `chains*(iter-warmup)`.

```
# stan options
chains = 3
rstan_options(auto_write = TRUE)
options(mc.cores = chains)
iter   =  4000
warmup =  2000
```

Then starting values for all free parameters are provided as a list (in case of multiple chains: a list of lists). For the time series intial states `N0sim`, data in `t[1]` scaled up from the sampled fraction `f` to the total volume are a natural choice. Remaining parameters are given guesses. Stan usually converges quickly towards the parameter region featuring a high probability density. But with ODEs involved, good initial guesses that do not cause numerical issues in ODE integration are often required.

```
# initial values for sampling
init=rep(list(list(r=0.1,
                   K=1e4,
                   N0sim=data$N[, 1]/data$f,
                   tau=10))
         ,chains)
```

Finally, the model is compiled and sampling is started.

```
stanmodel = stan_model(model_code=stanmodelcode)

fit = sampling(stanmodel,
               data=data,
               iter=iter,
               warmup=warmup,
               chains=chains,
               init=init
)
```

#### 1.5 Model diagnostics

Convergence of the MCMC chains is checked, e.g. by looking at estimated model parameters. We check the effective sample size `n_eff` and the Gelman-Rubin diagnostics `Rhat` (should be <1.01), indicating that chains have mixed.

```
print(fit, digits=3, pars=c("r","K"), probs=c(0.025, 0.5, 0.975))
```

```
## Inference for Stan model: 5fc336e64b9bc8d6e9fa54d21c97f38b.
## 3 chains, each with iter=4000; warmup=2000; thin=1;
## post-warmup draws per chain=2000, total post-warmup draws=6000.
##
##          mean se_mean     sd     2.5%       50%     97.5% n_eff Rhat
## r       0.098   0.000  0.002    0.094     0.098     0.102  1644    1
## K   10039.947   1.407 98.894 9848.147 10039.359 10234.801  4943    1
##
## Samples were drawn using NUTS(diag_e) at Mon Apr  4 13:37:01 2022.
## For each parameter, n_eff is a crude measure of effective sample size,
## and Rhat is the potential scale reduction factor on split chains (at
## convergence, Rhat=1).
```

Additionally, chains are inspected visually via trace- and density plots.

```
library("coda")
samples=As.mcmc.list(fit)
plot(samples[, c("r","K")])
```

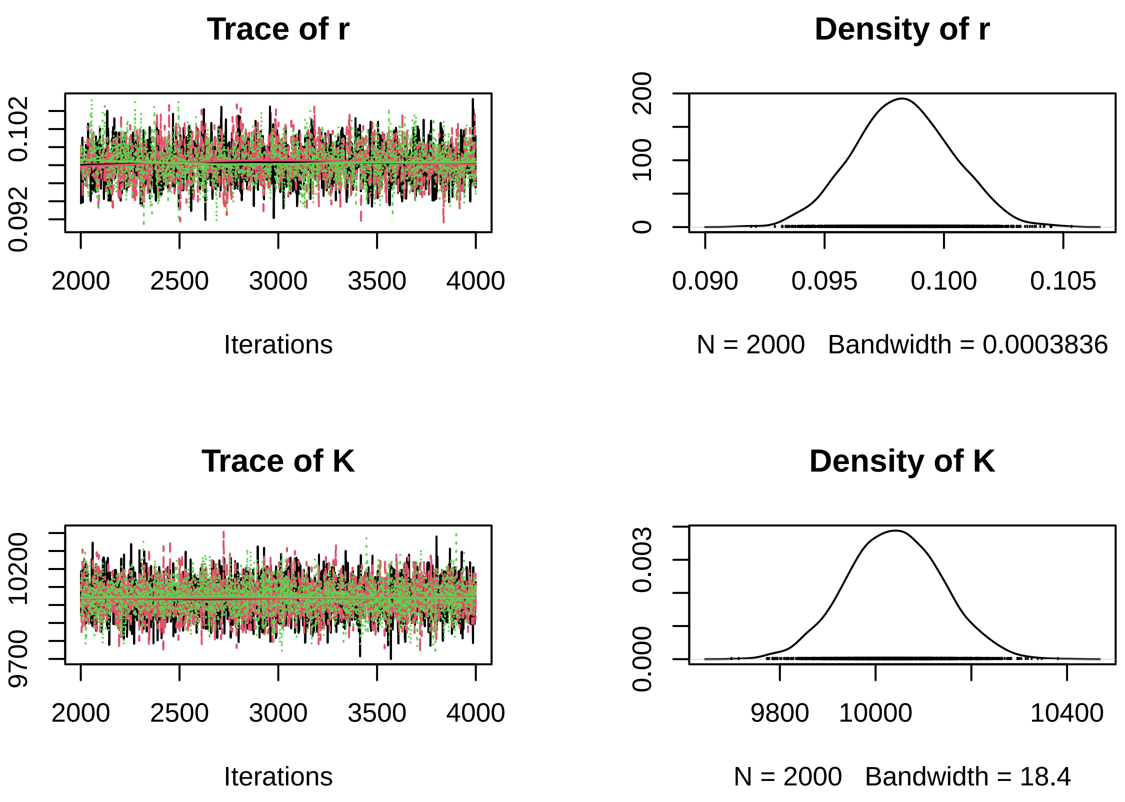


A pairs plot can detect correlations in model parameters. Almost perfect correlation would indicate non-identifiability.

```
pairs(fit, pars=c("r","K"))
```

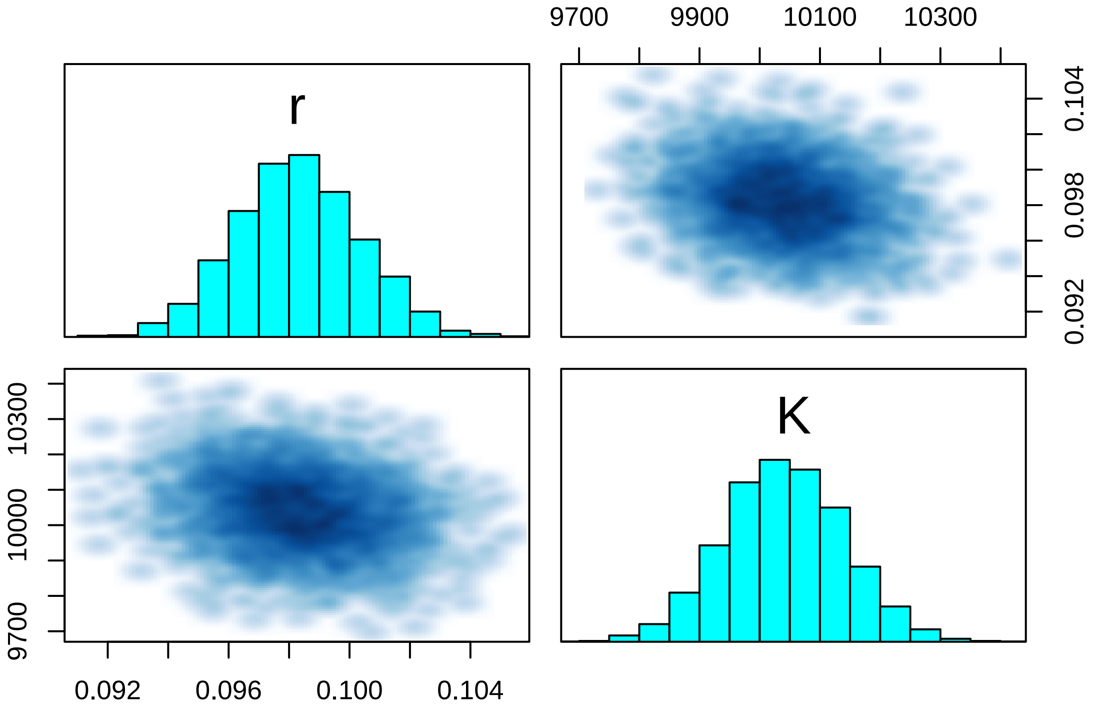


Optional: We also computed posterior samples for $\alpha = \frac{r}{K}$ in the `generated quantities{}` block.

```
print(fit, digits=7, pars=c("alpha"), probs=c(0.025, 0.5, 0.975))
```

```
## Inference for Stan model: 5fc336e64b9bc8d6e9fa54d21c97f38b.
## 3 chains, each with iter=4000; warmup=2000; thin=1;
## post-warmup draws per chain=2000, total post-warmup draws=6000.
```

```
##
##          mean se_mean    sd    2.5%     50%    97.5% n_eff     Rhat
## alpha 9.8e-06       0 2e-07 9.3e-06 9.8e-06 1.03e-05  1828 0.999713
##
## Samples were drawn using NUTS(diag_e) at Mon Apr  4 13:37:01 2022.
## For each parameter, n_eff is a crude measure of effective sample size,
## and Rhat is the potential scale reduction factor on split chains (at
## convergence, Rhat=1).
```

```
plot(samples[, "alpha"])
```

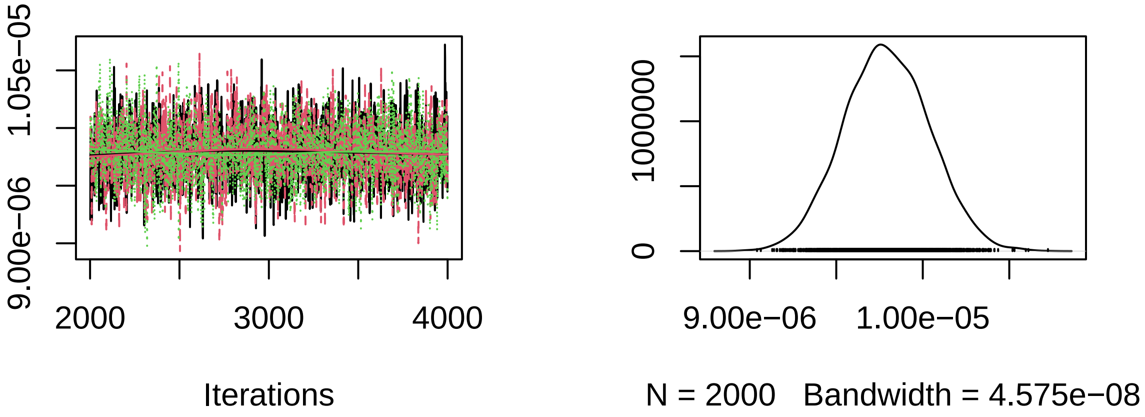


#### 1.6 Posterior predictions

Posterior predictions can generated from sampled posterior parameters. Alternatively, this can be done while sampling using `generated quantities{}` block in the model code. Here, we use the `deSolve` package for numerical integration to compute posterior predictions. First, we code an R function defining the population growth rate (ODE).

```
library("deSolve")

ode.model = function(t,N,p){
  dNdt =  p[1]*N[1]*(1.0-N[1]/p[2])
  return(list(dNdt))
}
```

Samples of the posterior distribution are converted to matrix. Each column contains all samples of a single parameter, while each row contains one sample from the multivariate posterior distribution.

```
post = as.matrix(fit)
head(post)
```

```
##           parameters
## iterations          r        K      tau   N0sim[1] N0sim[2] N0sim[3] N0sim[4]
##       [1,] 0.09376932 10186.42 209.2768 152.26885 120.2389 154.1004 154.2818
##       [2,] 0.09739955  9941.89 143.1738  99.11496 145.6647 130.1438 106.4487
##       [3,] 0.09969756 10179.57 151.2216 129.70831 104.0364 115.8647 124.5975
##       [4,] 0.09925819  9986.83 204.4218 104.97922 128.4086 160.1702 103.7960
##       [5,] 0.09714696 10152.46 186.0134 107.27475 118.1637 129.7602 123.9067
##       [6,] 0.09860314 10057.07 169.5889 113.77948 125.4977 134.1352 127.4644
##           parameters
## iterations N0sim[5] N0sim[6]        alpha     lp__
##       [1,] 63.49991 91.42657 9.205324e-06 477914.4
##       [2,] 77.89300 94.25055 9.796884e-06 477916.3
##       [3,] 58.00113 91.40117 9.793884e-06 477917.5
##       [4,] 65.18200 80.87335 9.938909e-06 477920.4
##       [5,] 54.94124 84.13801 9.568806e-06 477922.7
##       [6,] 70.38513 91.17071 9.804361e-06 477923.9
```

`t.pred` includes times for which predictions are generated. The empty matrix `N.pred` will store all predicted values: each row will contain one sample of a predicted timeseries.

```
n.post = 1000 # nrow(post)
t.pred = seq(from=min(data$t), to=max(data$t), by=1)
N.pred = matrix(NA, nrow=n.post, ncol=length(t.pred))
```

Predictions are generated for each time series replicate seperately (for-loop over `i`). For each sample from the posterior (row `j` in matrix `post[j, ]`), the trajectory is computed by numerically integrating the ODE with parameters `r` and `K`, starting in initial state `N0sim[i]`. Each column `k` of `N.pred[ ,k]` contains posterior predictions in `t.pred[k]`. We calculate quantiles of these predictions and plot them against the data, which are scaled up to the total volume by dividing by the sampling fraction: `data$N[,]/data$f`.

Note that predicted time series differ only in their initial states `N0sim[i]`, since model parameters `r` and `K` are identical across time series replicates $i$.

```
par(mfrow=c(3,2), mar=c(3,3,0,0), oma=c(3,3,1,1))
for(i in 1:m){
  for(j in 1:n.post){
    N.pred[j, ] = as.data.frame(lsoda(y     = c( N = post[j, paste0("N0sim[",i,"]")] ),
                                      times = t.pred,
                                      func  = ode.model,
                                      parms = c(post[j, "r"],
                                                post[j, "K"])
    ))$N
  }

  N.pred.qs = apply(N.pred, 2, function(x) quantile(x, probs=c(0.025,0.500,0.975), na.rm=TRUE))

  plot(data$t, data$N[i, ]/data$f,
             ylim=c(0,1.1e4), type="n", xlab="", ylab="")
  polygon(c(t.pred, rev(t.pred)), c(N.pred.qs[1, ], rev(N.pred.qs[3, ])),
          col = adjustcolor("red",alpha.f=0.25),  border = NA)
  lines(t.pred, N.pred.qs[2, ], col="red", lwd=1.5)
  points(data$t, data$N[i, ]/data$f)
}
mtext("Time [h]", side=1, outer=TRUE, line=1)
mtext("Abundance [ind]", side=2, outer=TRUE, line=1)
```

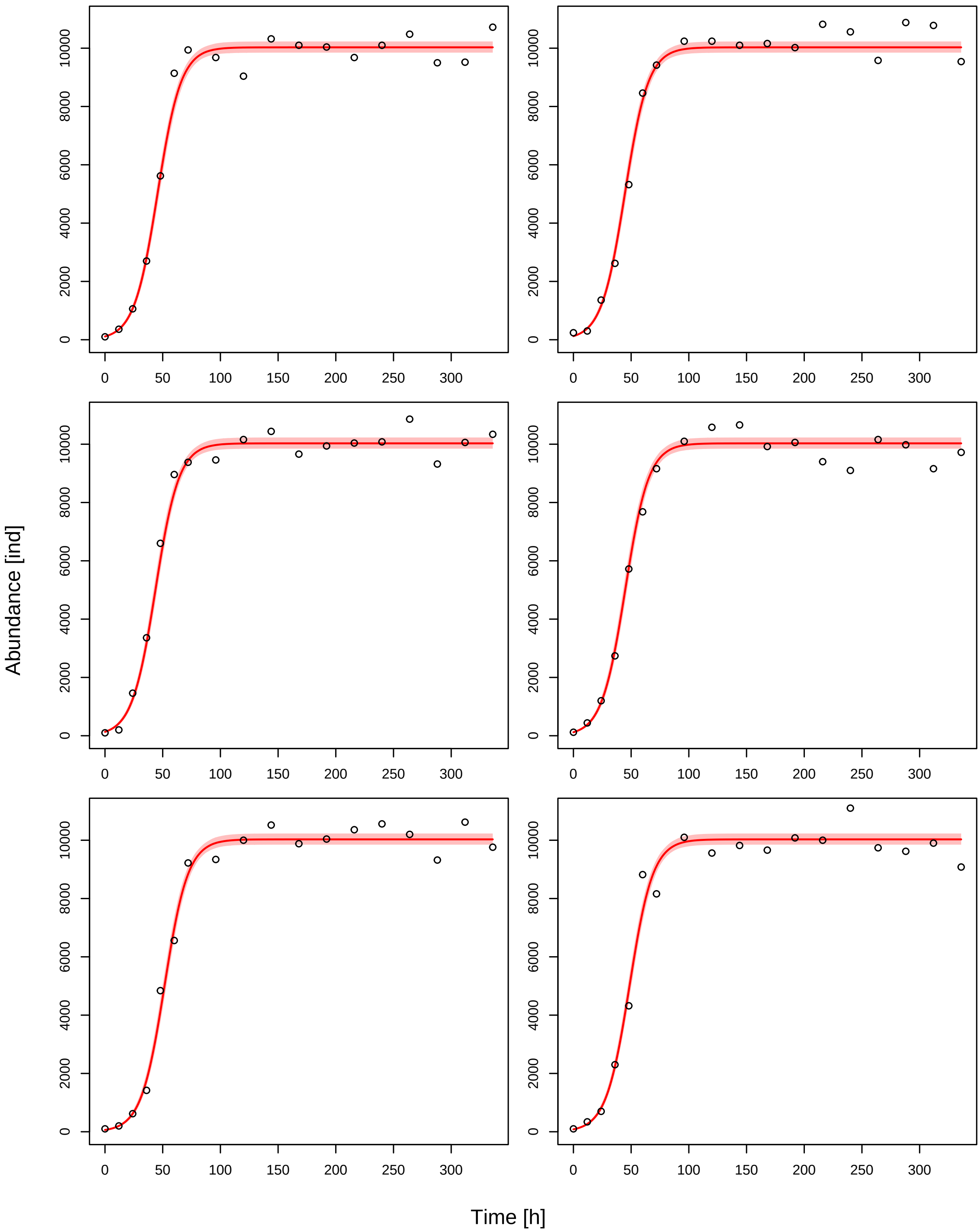
Abundance [ind]
Time [h]
0
2000
4000
6000
8000
10000
0
50
100
150
200
250
300

### 2 Logistic growth - PROC model

#### 2.1 Model

The logistic growth model can be written in the $r, \alpha$ or the $r, K$ formulation, where $K = \frac{r}{\alpha}$, or $\alpha = \frac{r}{K}$:

$$\frac{dN}{dt} = (r - \alpha N)N$$
$$= r(1 - \frac{N}{K})N$$

Using one-step-ahead fitting (process error only), we will fit all datapoints $N_t$ of $m$ timeseries replicates with predictions based on their previous observations $N_{t-1}$. A joint set of model parameters $r$ and $K$ is used.

#### 2.2 Data preparation

The dataset `data_example_logistic.csv` contains `m=6` timeseries with `n=18` observations of densities [ind] each, including their dates [h]. The fraction of the observed volume for sampling was `f=0.1`. Data preparation is identical to the OBS model manual.

```
rm(list=ls())
library("rstan")

data.in = read.csv("data_example_logistic.csv", row.names=1)

head(data.in)
```

```
##      X1 X2  X3  X4  X5  X6  X7   X8   X9  X10  X11  X12  X13  X14  X15  X16
## time  0 12  24  36  48  60  72   96  120  144  168  192  216  240  264  288
## N1   10 36 106 270 562 914 994  968  904 1032 1010 1004  968 1010 1048  950
## N2   24 30 136 262 532 846 942 1024 1024 1010 1016 1002 1082 1056  958 1088
## N3   10 20 146 336 660 896 938  946 1016 1044  966  994 1004 1008 1086  932
## N4   12 44 120 274 572 768 916 1010 1058 1066  992 1006  940  910 1016  998
## N5   10 20  62 142 484 656 922  934 1000 1052  988 1004 1036 1056 1020  932
##       X17  X18
## time  312  336
## N1    952 1072
## N2   1078  954
## N3   1006 1034
## N4    916  972
## N5   1062  976
```

```
n = ncol(data.in)
m = nrow(data.in)-1

times = as.numeric(data.in[1, ])
N = unname(as.matrix(data.in[2:(m+1), ]))

data = list(n = n,
            m = m,
            t = times,
            N = N,
            f = 0.1)

str(data)
```

```
## List of 5
##  $ n: int 18
##  $ m: num 6
##  $ t: num [1:18] 0 12 24 36 48 60 72 96 120 144 ...
```

```
##  $ N: int [1:6, 1:18] 10 24 10 12 10 10 36 30 20 44 ...
##  $ f: num 0.1
```

#### 2.3 Stan model

The Stan model is coded as a string, which is compiled later. A step by step walkthrough is given below.

```
stanmodelcode = '
functions{
  real[] odemodel(real t, real[] N, real[] p, real[] x_r, int[] x_i){
    // p[1]=r, p[2]=K
    real dNdt[1];
    dNdt[1] = p[1]*(1-(N[1]/p[2]))*N[1];
    return dNdt;
  }
}

data{
  int n; // observations
  int m; // replicates
  real t[n];
  int N[m,n];
  real f;
}

parameters{
  real<lower=0> r;
  real<lower=0> K;
  real<lower=0> tau;
}

model{
  real p[2];
  real Nsim[1,1]; // simulated values, matrix. dim1 = 1 point in time, dim2 = dim_ODE = 1

  // priors (uninformative)
  r ~ lognormal(-2,1);
  K ~ lognormal(9,1);
  tau ~ gamma(2,0.1);

  // parameters for integrator
  p[1] = r;
  p[2] = K;

  for(j in 1:m){
    for(i in 1:(n-1)){
      if(N[j,i]>0){
        // integrate ODE
        Nsim = integrate_ode_rk45(odemodel, {N[j,i]/f}, t[i], {t[i+1]}, p, rep_array(0.0,0), rep_arr
        // likelihood
        N[j,i+1] ~ neg_binomial_2(Nsim[1,1]*f,tau);
      }
    }
  }
}
'
```

##### 2.3.1 functions{ } block

This is identical to the OBS model manual.

```
real[] odemodel(real t, real[] N, real[] p, real[] x_r, int[] x_i){
  // p[1]=r, p[2]=K
  real dNdt[1];
  dNdt[1] = p[1]*(1-(N[1]/p[2]))*N[1];
  return dNdt;
}
```

##### 2.3.2 data{ } block

This is identical to the OBS model manual.

```
int n; // observations
int m; // replicates
real t[n];
int N[m,n];
```

##### 2.3.3 parameters{ } block

Here, all free parameters are declared. This includes model parameters `r` and `K`, and a parameter `tau` for the residual distribution (overdispersion of the negative binomial). All parameters are required to be positive.

```
real<lower=0> r;
real<lower=0> K;
real<lower=0> tau;
```

##### 2.3.4 model{ } block

This block defines how the posterior computed, using prior distributions and the likelihood function. First, some intermediate variables for the ODE simulation are declared. `p` is an array which stores model parameters `r` and `K`. `Nsim` will contain the output of the numerical integration with dimensions `1` (predict only next state), and the second dimension is number of states (here just 1).

```
real p[2];
real Nsim[1,1]; // simulated values, matrix. dim1 = 1 point in time, dim2 = dim_ODE = 1
```

Priors: This is identical to the OBS model manual.

```
r ~ lognormal(-2,1);
K ~ lognormal(9,1);
tau ~ gamma(2,0.1);
```

After the priors are defined, predictions are computed. This includes numerically integrating the ODE, which requires all model parameters are stored in one array `p`.

```
p[1] = r;
p[2] = K;
```

In a loop over all `m` time series replicates and `n-1` timepoints, numerical integration is performed, to predict the next state `N[j,i+1]` from the current state `N[j,i]`. Observed data is abundance in sampling fraction `f`, so the initial values are scaled up to the total volume (`N[j,i]/f`) for numerical integration, and predictions are scaled down later to the sampling fraction (`Nsim[1,1]*f`) for calculating residuals. The integration routine `integrate_ode_rk45()` is must be called in a specific format. Arguments are the function `odemodel` as defined in the `functions{}` block. The initial state is converted to an array using `{ }`. Integration is performed from `t[i]` to `t[i+1]`. `p` contains the model parameters. For additional variables `x_r` and `x_i` (mandatory but not used here), empty arrays are generated.

After computing the predicted state `Nsim` (in `t[i+1]`), it is confronted with the data `N[j,i+1]`: a negative binomial distribution defines likelihood values. Since the data was sampled in a fraction `f=0.1`, the distribution's mean is `Nsim[,]*f`, with some overdispersion `tau`.

The prediction is only performed if `N[j,i]>0`. `N[j,i]=0` would predict a 0 in `t[i+1]` irrespective of the model parameters `r` and `K`, so it wouldn't contribute any information to the likelihood function.

```
for(j in 1:m){
  for(i in 1:(n-1)){
    if(N[j,i]>0){
      // integrate ODE
      Nsim = integrate_ode_rk45(odemodel, {N[j,i]/f}, t[i], {t[i+1]}, p, rep_array(0.0,0), rep_arr
      // likelihood
      N[j,i+1] ~ neg_binomial_2(Nsim[1,1]*f,tau);
    }
  }
}
```

#### 2.4 Model fitting

This is identical to the OBS model manual.

```
# stan options
chains = 3
rstan_options(auto_write = TRUE)
options(mc.cores = chains)
iter   =  4000
warmup =  2000
```

Then starting values for all free parameters are provided as a list (in case of multiple chains: a list of lists) with initial guesses. Stan usually converges quickly towards the parameter region featuring a high probability density. But with ODEs involved, good initial guesses that do not cause numerical issues in ODE integration are often required.

```
# initial values for sampling
init=rep(list(list(r=0.1,
                   K=1e4,
                   tau=10))
         ,chains)
```

Finally, the model is compiled and sampling is started.

```
stanmodel = stan_model(model_code=stanmodelcode)

fit = sampling(stanmodel,
               data=data,
               iter=iter,
               warmup=warmup,
               chains=chains,
               init=init
)
```

#### 2.5 Model diagnostics

Convergence of the MCMC chains is checked, e.g. by looking at estimated model parameters. We check the effective sample size `n_eff` and the Gelman-Rubin diagnostics `Rhat` (should be <1.01), indicating that chains have mixed.

```
print(fit, digits=3, pars=c("r","K"), probs=c(0.025, 0.5, 0.975))
```

```
## Inference for Stan model: b16cd10e0c1eaeb90e8e546724002b41.
## 3 chains, each with iter=4000; warmup=2000; thin=1;
## post-warmup draws per chain=2000, total post-warmup draws=6000.
##
##        mean se_mean      sd     2.5%      50%     97.5% n_eff  Rhat
## r     0.101   0.000   0.003    0.094     0.10     0.107  4940 1.001
```

```
## K 10021.477   2.206 154.556 9723.839 10019.26 10325.511  4909 1.000
##
## Samples were drawn using NUTS(diag_e) at Mon Apr  4 13:46:46 2022.
## For each parameter, n_eff is a crude measure of effective sample size,
## and Rhat is the potential scale reduction factor on split chains (at
## convergence, Rhat=1).
```

Additionally, chains are inspected visually via trace- and density plots.

```
library("coda")
samples=As.mcmc.list(fit)
plot(samples[, c("r","K")])
```

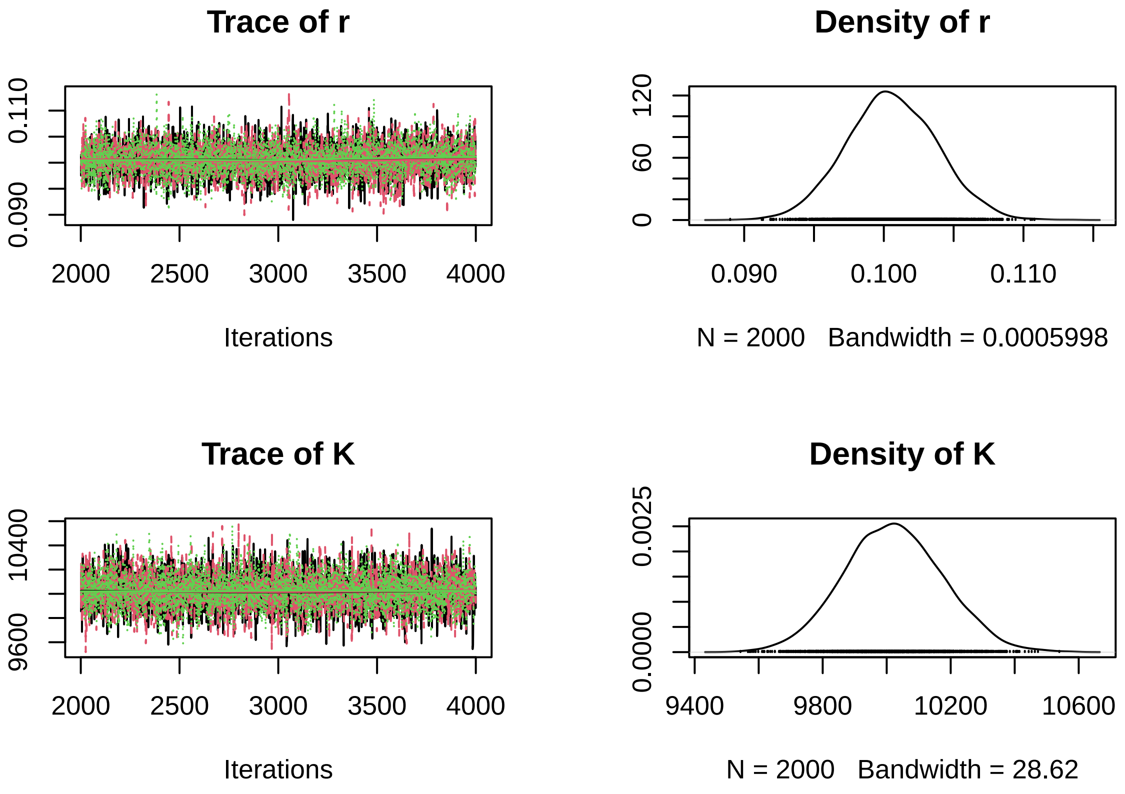


A pairs plot can detect correlations in model parameters. Almost perfect correlation would indicate non-identifiability.

```
pairs(fit, pars=c("r","K"))
```

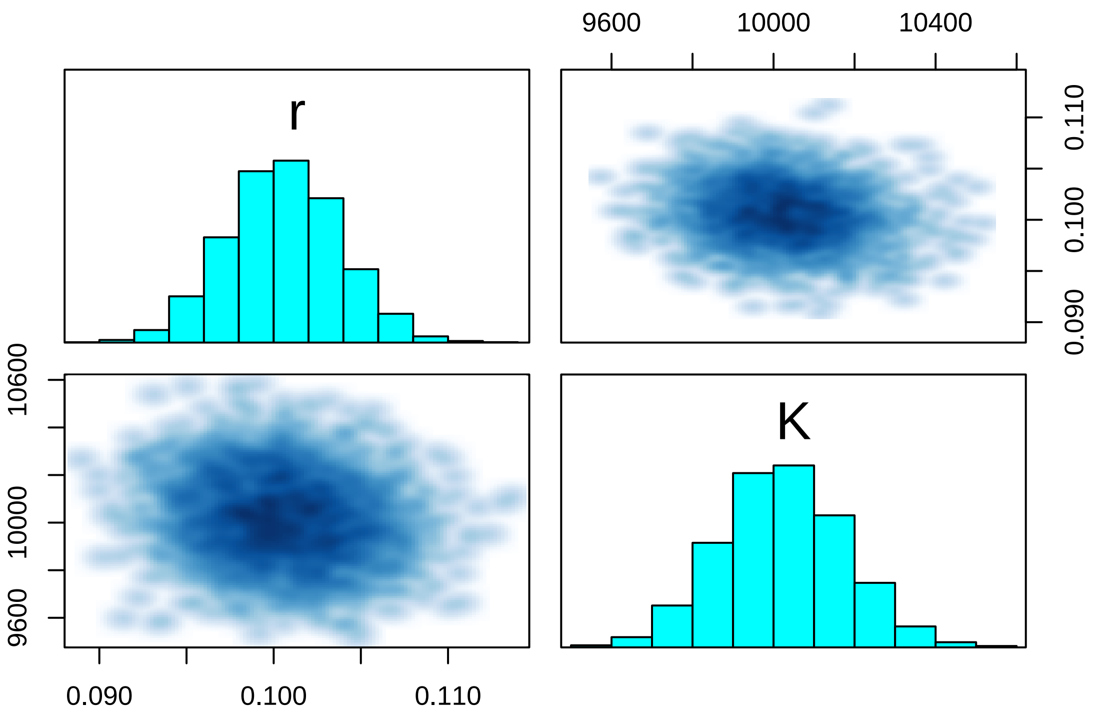


#### 2.6 Posterior predictions

Posterior predictions are generated one-step-ahead from sampled posterior parameters. Alternatively, this can be done while sampling using `generated quantities{}` block in the model code. Here, we use the `deSolve` package for numerical integration to compute posterior predictions. First, we code an R function defining the population growth rate (ODE).

```
library("deSolve")
ode.model = function(t,N,p){
  dNdt =  p[1]*N[1]*(1.0-N[1]/p[2])
  return(list(dNdt))
}
```

Samples of the posterior distribution are converted to matrix. Each column contains all samples of a single parameter, while each row contains one sample from the multivariate posterior distribution.

```
post = as.matrix(fit)
# head(post)
```

The empty matrix `N.pred` will store all predicted values: each row will contain one sample of a predicted timeseries. Predictions are computed for each time series replicate (for-loop over `i`) and each timepoint (for-loop over `k`) separately. For each sample from the posterior (row `j` in matrix `post[j, ]`), each prediction is computed by numerically integrating the ODE with parameters `r` and `K`, starting in last observation `data$N[i,k]/data$f` (scaled up to total volume). Each column `k` of `N.pred[ ,k]` contains posterior predictions in `t[k+1]`. We calculate quantiles of these predictions and plot them against the data.

Since these are one-step-ahead predictions from the last observation, we plot them pointwise in time, rather than a trajectory as in the OBS model.

```
post = as.matrix(fit)
n.post = nrow(post)
n.post = 1000

par(mfrow=c(3,2), mar=c(3,3,0,0), oma=c(3,3,1,1))
for(i in 1:m){

  N.pred = matrix(NA, nrow=n.post, ncol=length(data$t)-1)
```

```
  for(j in 1:n.post){
    for(k in 1:(length(data$t)-1)){
      N.pred[j,k] = as.data.frame(lsoda(y     = c( N = data$N[i,k]/data$f ),
                                        times = data$t[c(k,k+1)],
                                        func  = ode.model,
                                        parms = c(post[j, "r"],
                                                  post[j, "K"])
      ))$N[2]
    }
  }
  N.pred.qs = apply(N.pred, 2, function(x) quantile(x, probs=c(0.025,0.500,0.975), na.rm=TRUE))

  plot(data$t[2:n], data$N[i,2:n]/data$f,ylim=c(0,1.1e4), type="n", xlab="", ylab="")
  for(k in 1:(n-1)){
    lines(c(data$t[k+1],data$t[k+1]), c(N.pred.qs[1,k],N.pred.qs[3,k]), col="red", lwd=1.5)
  }
  points(data$t[2:n], N.pred.qs[2, ], col="red", pch="-", cex=1.5)
  points(data$t[2:n], data$N[i,2:n]/data$f, type="p")
}
mtext("Time [h]", side=1, outer=TRUE, line=1)
mtext("Abundance [ind]", side=2, outer=TRUE, line=1)
```

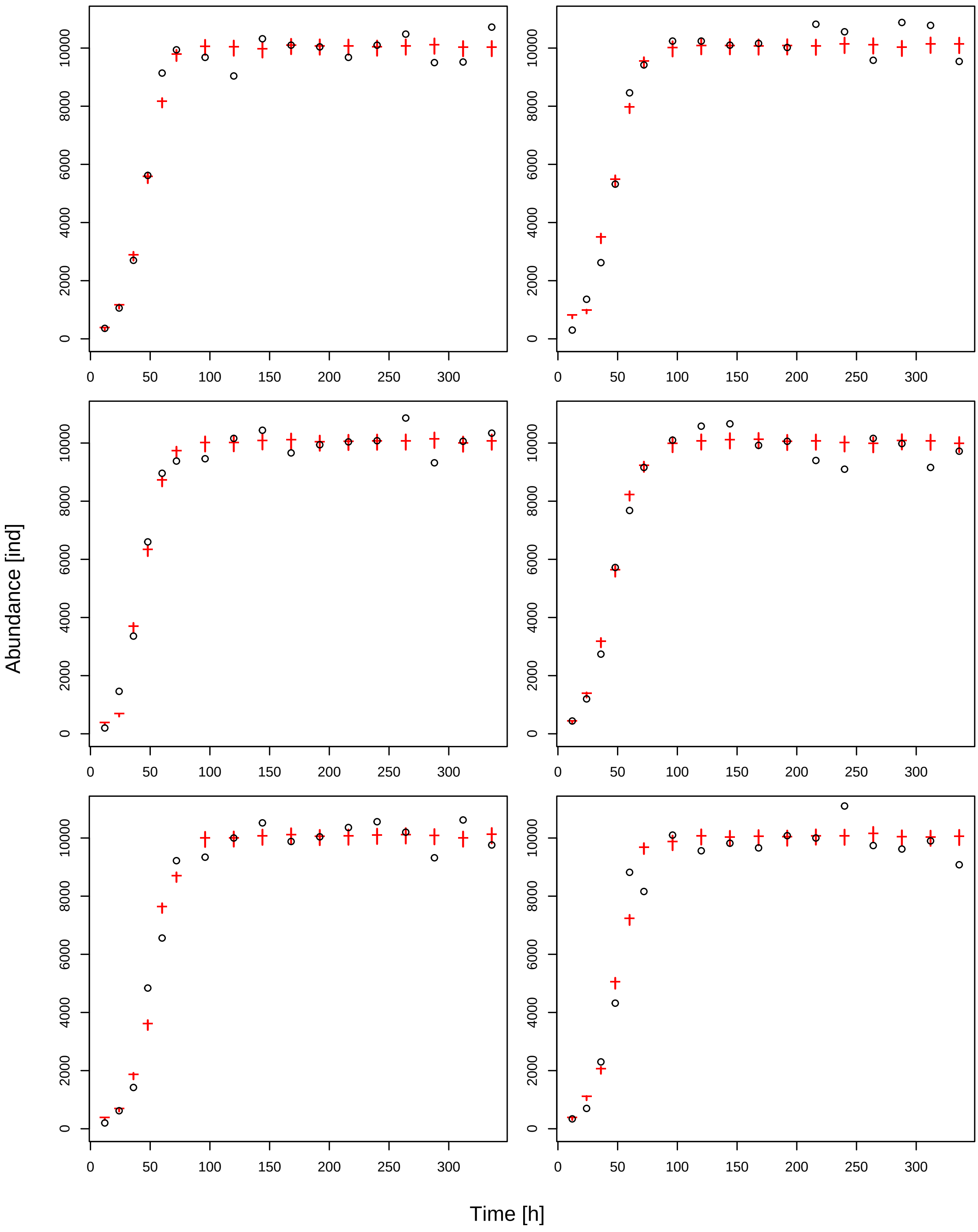
Abundance [ind]
Time [h]
0
2000
4000
6000
8000
10000
0
50
100
150
200
250
300

### 3 Logistic growth - State-space model

#### 3.1 Model

The logistic growth model can be written in the $r, \alpha$ or the $r, K$ formulation, where $K = \frac{r}{\alpha}$, or $\alpha = \frac{r}{K}$:

$$\begin{aligned}\frac{dN}{dt} &= (r - \alpha N)N \\ &= r(1 - \frac{N}{K})N\end{aligned}$$

Using a state-space model (including observation and process errors), we will fit all $m$ time series replicates with a joint set of parameters $r$ and $K$ ("complete pooling"). For each observation $N_{j,i}$ in $t_i$ of time series $j$, underlying true states `Ntrue[j,i]` are free parameters, which are estimated too.

#### 3.2 Data preparation

The dataset `data_example_logistic.csv` contains `m=6` timeseries with `n=18` observations of densities [ind] each, including their dates [h]. The fraction of the observed volume for sampling was `f=0.1`. Data preparation is identical to the OBS model manual.

```
rm(list=ls())
library("rstan")

data.in = read.csv("data_example_logistic.csv", row.names=1)

head(data.in)
```

```
##      X1 X2  X3  X4  X5  X6  X7   X8   X9  X10  X11  X12  X13  X14  X15  X16
## time  0 12  24  36  48  60  72   96  120  144  168  192  216  240  264  288
## N1   10 36 106 270 562 914 994  968  904 1032 1010 1004  968 1010 1048  950
## N2   24 30 136 262 532 846 942 1024 1024 1010 1016 1002 1082 1056  958 1088
## N3   10 20 146 336 660 896 938  946 1016 1044  966  994 1004 1008 1086  932
## N4   12 44 120 274 572 768 916 1010 1058 1066  992 1006  940  910 1016  998
## N5   10 20  62 142 484 656 922  934 1000 1052  988 1004 1036 1056 1020  932
##       X17  X18
## time  312  336
## N1    952 1072
## N2   1078  954
## N3   1006 1034
## N4    916  972
## N5   1062  976
```

```
n = ncol(data.in)
m = nrow(data.in)-1

times = as.numeric(data.in[1, ])
N = unname(as.matrix(data.in[2:(m+1), ]))

data = list(n = n,
            m = m,
            t = times,
            N = N,
            f = 0.1)

str(data)
```

```
## List of 5
##  $ n: int 18
##  $ m: num 6
```

```
##  $ t: num [1:18] 0 12 24 36 48 60 72 96 120 144 ...
##  $ N: int [1:6, 1:18] 10 24 10 12 10 10 36 30 20 44 ...
##  $ f: num 0.1
```

#### 3.3 Stan model

The Stan model is coded as a string, which is compiled later. A step by step walkthrough is given below.

```
stanmodelcode = '
functions{
  real[] odemodel(real t, real[] N, real[] p, real[] x_r, int[] x_i){
    // p[1]=r, p[2]=K
    real dNdt[1];
    dNdt[1] = p[1]*(1-(N[1]/p[2]))*N[1];
    return dNdt;
  }
}

data{
  int n; // observations
  int m; // replicates
  real t[n];
  int N[m,n];
  real f;
}

parameters{
  real<lower=0> r;
  real<lower=0> K;
  real<lower=0> sigma;
  real<lower=0> tau;
  real<lower=0> Ntrue[m,n]; // states in times t
}

model{
  real p[2];
  real Nsim[1,1]; // simulated values, matrix. dim1 = 1 point in time, dim2 = dim_ODE = 1

  // priors
  r ~ lognormal(-2,1);
  K ~ lognormal(9,1);
  sigma ~ gamma(2,1);
  tau ~ gamma(2,0.1);

  for(i in 1:m){
    Ntrue[i,1] ~ normal(0,100);
  }

  // parameters for integrator
  p[1] = r;
  p[2] = K;

  // process equations
  for(j in 1:m){
    for(i in 1:(n-1)){
      // integrate ODE
      Nsim = integrate_ode_rk45(odemodel, {Ntrue[j,i]}, t[i], {t[i+1]}, p, rep_array(0.0,0), rep_arr
      // process error
      Ntrue[j,i+1] ~ lognormal(log(Nsim[1,1]), sigma);
```

```
    }
  }

  // observation equations
  for(j in 1:m){
    for(i in 1:n){
      N[j,i] ~ neg_binomial_2(Ntrue[j,i]*f, tau);
    }
  }
}
'
```

##### 3.3.1 functions{ } block

This is identical to the OBS model manual.

```
real[] odemodel(real t, real[] N, real[] p, real[] x_r, int[] x_i){
  // p[1]=r, p[2]=K
  real dNdt[1];
  dNdt[1] = p[1]*(1-(N[1]/p[2]))*N[1];
  return dNdt;
}
```

##### 3.3.2 data{ } block

This is identical to the OBS model manual.

```
int n; // observations
int m; // replicates
real t[n];
int N[m,n];
real f;
```

##### 3.3.3 parameters{ } block

Here, all free parameters are declared. This includes model parameters `r` and `K`, and parameters `sigma` for the process residual distribution (scale parameter of lognormal) and `tau` for the observation residual distribution (overdispersion of the negative binomial). Additionally, for all `m` x `n` observations, the underlying true state `Ntrue` is estimated. All parameters are required to be positive.

```
real<lower=0> r;
real<lower=0> K;
real<lower=0> sigma;
real<lower=0> tau;
real<lower=0> Ntrue[m,n]; // states in times t
```

##### 3.3.4 model{ } block

This block defines how the posterior computed, using prior distributions and the likelihood function. First, some intermediate variables for the ODE simulation are declared. `p` is an array which stores model parameters `r` and `K`. `Nsim` will contain the output of the numerical integration with dimensions `1` (predict only next state), and the second dimension is number of states (here just 1).

```
real p[2];
real Nsim[1,1]; // simulated values, matrix. dim1 = 1 point in time, dim2 = dim_ODE = 1
```

Priors: This is identical to the OBS and PROC model manual, initial densities parameters `Ntrue[i,1]` are assigned a weak prior, too.

```
r ~ lognormal(-2,1);
K ~ lognormal(9,1);
sigma ~ gamma(2,1);
```

```
tau ~ gamma(2,0.1);

for(i in 1:m){
  Ntrue[i,1] ~ normal(0,100);
}
```

After the priors are defined, predictions are computed. This includes numerically integrating the ODE, which requires all model parameters are stored in one array `p`.

```
p[1] = r;
p[2] = K;
```

The likelihood consists of two parts: the process equations and the observation equations.

In a first loop over all `m` time series replicates and `n-1` timepoints, numerical integration is performed, to predict the next true state `Ntrue[j,i+1]` from the current true state `Ntrue[j,i]`. The integration routine `integrate_ode_rk45()` is must be called in a specific format. Arguments are the function `odemodel` as defined in the `functions{}` block. The initial state is converted to an array using `{ }`. Integration is performed from `t[i]` to `t[i+1]`. `p` contains the model parameters. For additional variables `x_r` and `x_i` (mandatory but not used here), empty arrays are generated.

After computing the predicted state `Nsim` (in `t[i+1]`), it is confronted with estimate of `Ntrue[j,i+1]`: we assume a lognormal distribution, `sigma` denoting the process error.

```
// process equations
for(j in 1:m){
  for(i in 1:(n-1)){
    // integrate ODE
    Nsim = integrate_ode_rk45(odemodel, {Ntrue[j,i]}, t[i], {t[i+1]}, p, rep_array(0.0,0), rep_arr
    // process error
    Ntrue[j,i+1] ~ lognormal(log(Nsim[1,1]), sigma);
  }
}
```

Second, the true states estimates `Ntrue[j,i]` are confronted with the data `N[j,i]`: a negative binomial distribution defines likelihood values. Since the data was sampled in a fraction `f=0.1`, the distribution's mean is `Ntrue[,]*f`, with some overdispersion `tau`.

```
// observation equations
for(j in 1:m){
  for(i in 1:n){
    N[j,i] ~ neg_binomial_2(Ntrue[j,i]*f, tau);
  }
}
```

#### 3.4 Model fitting

This is identical to the OBS model manual.

```
# stan options
chains = 3
rstan_options(auto_write = TRUE)
options(mc.cores = chains)
iter   =  4000
warmup =  2000
```

Then starting values for all free parameters are provided as a list (in case of multiple chains: a list of lists). For the time series true states estimates `Ntrue`, observed data scaled up from the sampled fraction `f` to the total volume are a natural choice. Remaining parameters are given guesses. Stan usually converges quickly towards the parameter region featuring a high probability density. But with ODEs involved, good initial guesses that do not cause numerical issues in ODE integration are often required.

```
# initial values for sampling
init=rep(list(list(r=0.1,
                   K=1e4,
                   sigma=0.1,
                   tau=10,
                   Ntrue=data$N/data$f
))
,chains)
```

Finally, the model is compiled and sampling is started.

```
stanmodel = stan_model(model_code=stanmodelcode)

fit = sampling(stanmodel,
               data=data,
               iter=iter,
               warmup=warmup,
               chains=chains,
               init=init
)
```

#### 3.5 Model diagnostics

Convergence of the MCMC chains is checked, e.g. by looking at estimated model parameters. We check the effective sample size `n_eff` and the Gelman-Rubin diagnostics `Rhat` (should be <1.01), indicating that chains have mixed.

```
print(fit, digits=3, pars=c("r","K"), probs=c(0.025, 0.5, 0.975))
```

```
## Inference for Stan model: c7c04773fe5c1c7b19f59f387bb18f00.
## 3 chains, each with iter=4000; warmup=2000; thin=1;
## post-warmup draws per chain=2000, total post-warmup draws=6000.
##
##        mean se_mean      sd     2.5%       50%     97.5% n_eff  Rhat
## r     0.098   0.000   0.002    0.094     0.098     0.102  1062 1.001
## K 10040.881   4.384 102.881 9838.075 10042.381 10243.292   551 1.004
##
## Samples were drawn using NUTS(diag_e) at Tue Apr  5 13:17:06 2022.
## For each parameter, n_eff is a crude measure of effective sample size,
## and Rhat is the potential scale reduction factor on split chains (at
## convergence, Rhat=1).
```

Additionally, chains are inspected visually via trace- and density plots.

```
library("coda")
samples=As.mcmc.list(fit)
plot(samples[, c("r","K")])
```

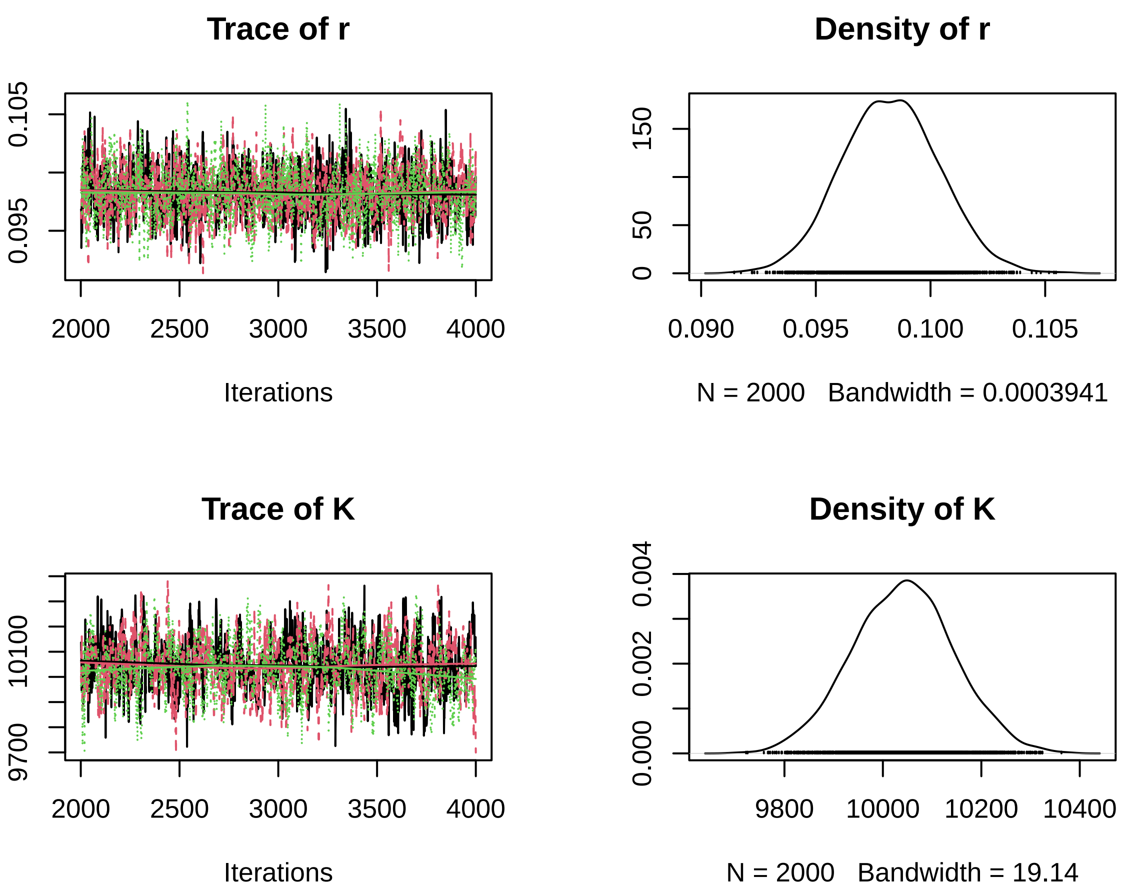


A pairs plot can detect correlations in model parameters. Almost perfect correlation would indicate non-identifiability.

```
pairs(fit, pars=c("r","K"))
```

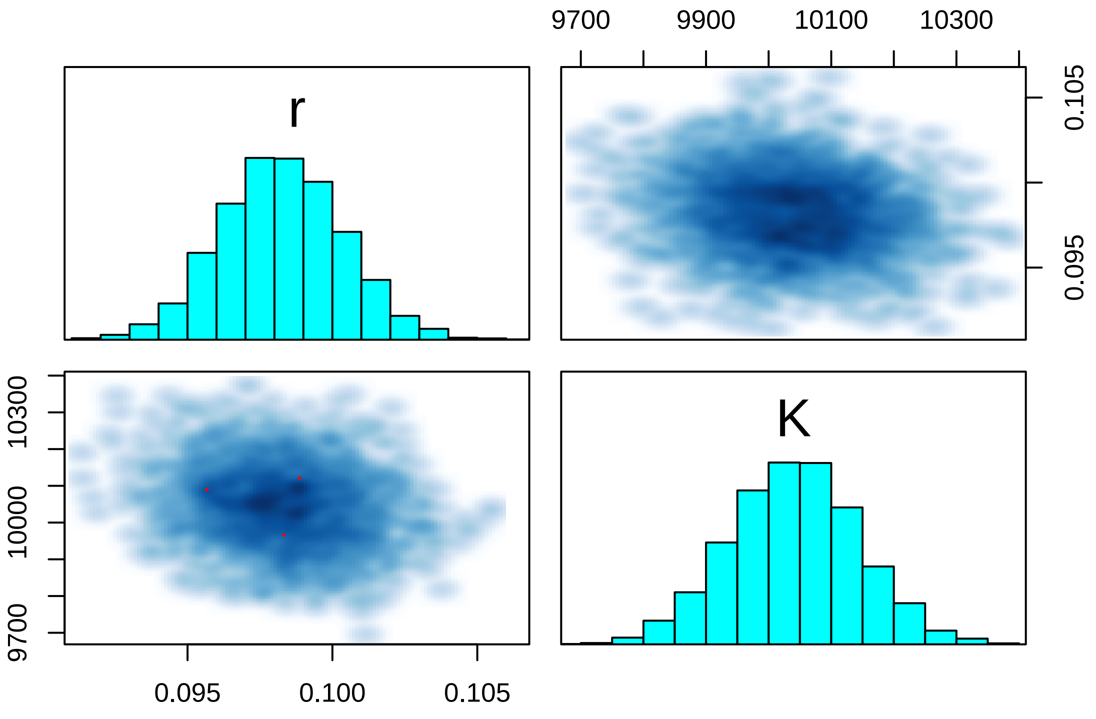

#### 3.6 Posterior predictions

Since underlying true states `Ntrue[i,j]` for all observations `N[i,j]` are already estimated, we can directly pull their means, medians or CIs from the summary table. We plot them against the data (upscaled to total volume `data$N[,]/data$f`), for simplicity as piecewise linear interpolations between observations.

```
summary = summary(fit)$summary

head(summary)
```

```
##                     mean      se_mean           sd         2.5%          25%
## r           9.825622e-02 6.497887e-05 2.117949e-03 9.417245e-02 9.680376e-02
## K           1.004088e+04 4.384263e+00 1.028807e+02 9.838075e+03 9.970937e+03
## sigma       1.852847e-02 5.506668e-04 7.154895e-03 7.871636e-03 1.320694e-02
## tau         1.776427e+02 4.422639e-01 3.146493e+01 1.228155e+02 1.550623e+02
## Ntrue[1,1] 1.121996e+02 3.348991e-01 1.113495e+01 9.176605e+01 1.043634e+02
## Ntrue[1,2] 3.555216e+02 8.201175e-01 2.952211e+01 3.009543e+02 3.348099e+02
##                      50%          75%        97.5%     n_eff     Rhat
## r           9.824891e-02 9.966586e-02 1.024132e-01 1062.3963 1.000884
## K           1.004238e+04 1.010909e+04 1.024329e+04  550.6482 1.003894
## sigma       1.744657e-02 2.257381e-02 3.565835e-02  168.8219 1.049309
## tau         1.752364e+02 1.972432e+02 2.458085e+02 5061.6297 1.000223
## Ntrue[1,1] 1.117634e+02 1.194843e+02 1.348983e+02 1105.4741 1.001460
## Ntrue[1,2] 3.545287e+02 3.760437e+02 4.160335e+02 1295.8135 1.001247
```

```
N.pred.qs = matrix(NA, nrow=3, ncol=length(data$t))

par(mfrow=c(3,2), mar=c(3,3,0,0), oma=c(3,3,1,1))

for(i in 1:m){
  for(j in 1:length(times)){
    N.pred.qs[, j] = summary[paste0("Ntrue[",i,",",j,"]"), c("2.5%", "50%", "97.5%") ]
  }

  plot(data$t, data$N[i, ]/data$f, type="n", ylim=c(0,1.1e4), log="")
  polygon(c(data$t, rev(data$t)), c(N.pred.qs[1, ], rev(N.pred.qs[3, ])),
          col = adjustcolor("red",alpha.f=0.25),  border = NA)
  lines(data$t, N.pred.qs[2, ], col="red", lwd=1.5)
  points(data$t, data$N[i, ]/data$f)
}
mtext("Time [h]", side=1, outer=TRUE, line=1)
mtext("Abundance [ind]", side=2, outer=TRUE, line=1)
```

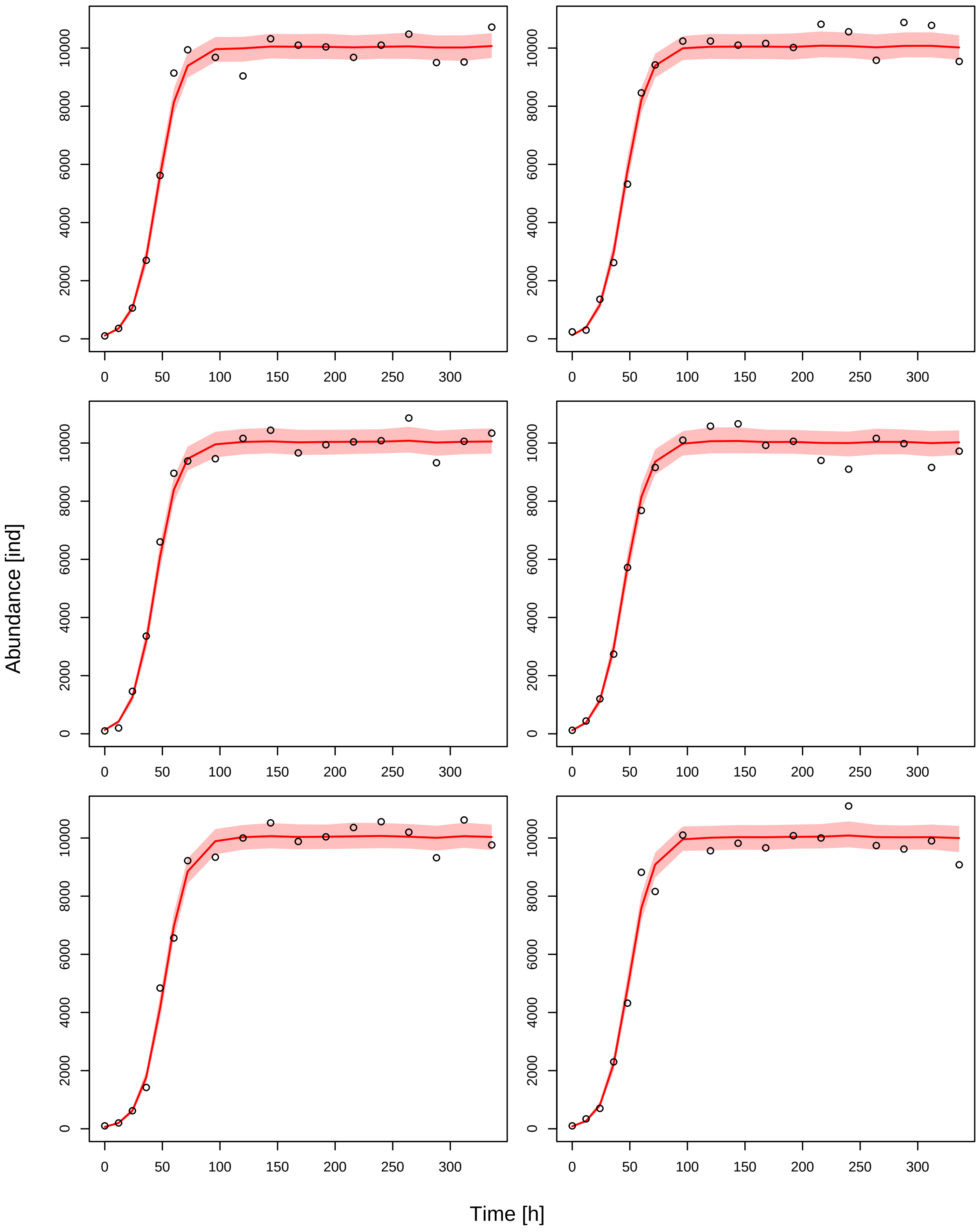
Abundance [ind]
Time [h]
0
2000
4000
6000
8000
10000
0
50
100
150
200
250
300

### 4 Consumer resource - OBS model

#### 4.1 Model

The Rosenzweig-MacArthur predator-prey model with logistic growth and Type II functional response reads

$$\frac{dN_1}{dt} = r(1 - \frac{N_1}{K})N_1 - \frac{ahN_1}{1 + ahN_1}N_2$$
$$\frac{dN_2}{dt} = e\frac{ahN_1}{1 + ahN_1}N_2 - dN_2$$

Using the trajectory fitting method (observation error only), we will fit all $m$ two-species time series replicates with a joint set of parameters $r, K, a, h, e, d$ ("complete pooling"), while allowing for $m$ individual initial states $N_{1,0} = N_1(t_0)$ and $N_{2,0} = N_2(t_0)$ as free parameters.

Additionally, we fit trajectories for $m_c$ control (single-species) time series of prey ($N_2 = 0$, logistic growth)

$$\frac{dN_1}{dt} = r(1 - \frac{N_1}{K})N_1$$

and $m_c$ control time series of predator ($N_1 = 0$, exponential decline)

$$\frac{dN_2}{dt} = -dN_2$$

These control data will provide additional information on $r, K$ and $d$.

#### 4.2 Data preparation

The dataset `data_example_consumer_resource.csv` contains `m=6` timeseries with `n=18` observations of densities [ind] each, including their dates [h]. The fraction of the observed volume for sampling was `f=0.1`.

```
rm(list=ls())
library("rstan")

data.in = read.csv("data_example_consumer_resource.csv", row.names=1)

head(data.in)
```

```
##       X1  X2  X3  X4  X5   X6   X7   X8   X9 X10 X11 X12  X13  X14  X15  X16
## time   0  12  24  36  48   60   72   96  120 144 168 192  216  240  264  288
## N1   200 207 228 370 664  958 1330 1253 1017 827 546 276  240  409 1094 1452
## N2   189 205 283 461 716 1002 1212 1032  610 227 245 600 1128 1383 1330  862
## N3   201 222 280 394 595  822 1097 1097  779 422 303 431 1001 1439 1161  912
## N4   198 214 241 428 641  924 1281 1413 1261 736 288 214  361  761 1168 1036
## N5   191 204 274 483 772 1060 1251 1100  688 333 267 612 1320 1595 1542 1128
##       X17 X18
## time  312 336
## N1   1291 903
## N2    329 119
## N3    400 146
## N4    596 285
## N5    817 389
```

For Stan, the data has to be coded as a named list, including dimensions of the data `n` and `m`, `times` as a vector. The `m` x `n` matrices `N1` and `N2` contain the observed densities of prey and predator, respectively, for the two-species mixtures. The `mc` x `n` matrices `N1C` and `N2C` contain the observed single-species densities of prey and predator, respectively.

```
n = 18
m = 6
mc = 3

times = as.numeric(data.in[1, ])
N1 = unname(as.matrix(data.in[2:(m+1), ]))
N2 = unname(as.matrix(data.in[(m+2):(2*m+1), ]))
N1C = unname(as.matrix(data.in[(2*m+2):(2*m+1+mc), ]))
N2C = unname(as.matrix(data.in[(2*m+2+mc):((2*m+1+2*mc)), ]))

data = list(n = n,
            m = m,
            mc = mc,
            t = times,
            N1 = N1,
            N2 = N2,
            N1C  = N1C,
            N2C  = N2C,
            f = 0.1)


str(data)
```

```
## List of 9
##  $ n  : num 18
##  $ m  : num 6
##  $ mc : num 3
##  $ t  : num [1:18] 0 12 24 36 48 60 72 96 120 144 ...
##  $ N1 : int [1:6, 1:18] 200 189 201 198 191 174 207 205 222 214 ...
##  $ N2 : int [1:6, 1:18] 48 38 41 42 56 40 35 29 27 34 ...
##  $ N1C: int [1:3, 1:18] 40 51 51 96 144 117 259 370 411 637 ...
##  $ N2C: int [1:3, 1:18] 195 185 201 104 98 105 58 56 44 40 ...
##  $ f  : num 0.1
```

#### 4.3 Stan model

The Stan model is coded as a string, which is compiled later. A step by step walkthrough is given below.

```
stanmodelcode='
functions{
  real[] odemodel(real t, real[] N, real[] p, real[] x_r, int[] x_i){
    // p[1]=r, p[2]=K, p[3]=b, p[4]=h, p[5]=e, p[6]=d
    real dNdt[2];
    dNdt[1] = p[1]*N[1]*(1.0-N[1]/p[2])-p[3]*N[1]*N[2]/(1.0+p[3]*p[4]*N[1]);
    dNdt[2] = p[5]*p[3]*N[1]*N[2]/(1.0+p[3]*p[4]*N[1])-p[6]*N[2];
    return dNdt;
  }
}

data{
  int n; // observations
  int m; // replicates
  int mc; // replicates control
  real t[n];
  int N1[m,n];
  int N2[m,n];
  int N1C[mc,n];
  int N2C[mc,n];
  real f;
```

```
}

parameters{
  real<lower=0> r;
  real<lower=0> K;
  real<lower=0> b;
  real<lower=0> h;
  real<lower=0, upper=1> e;
  real<lower=0> d;
  real<lower=0> tau[2];
  real<lower=0> N10sim[m]; // individual initial values for replicates
  real<lower=0> N20sim[m]; // individual initial values for replicates
  real<lower=0> N1C0sim[mc]; // individual initial values for replicates
  real<lower=0> N2C0sim[mc]; // individual initial values for replicates
}

model{
  real p[6];
  real Nsim[n-1,2]; // simulated values, matrix. dim1 = time without t0, dim2 = dim_ODE = 2
  real NCsim[n-1,1]; // simulated values, matrix. dim1 = time without t0, dim2 = dim_ODE = 1

  // priors
  r ~ lognormal(-2,1);
  K ~ lognormal(9,1);
  b ~ gamma(2,1);
  h ~ gamma(2,1);
  e ~ uniform(0,1);
  d ~ gamma(2,1);
  tau ~ gamma(2,0.1);

  //---------------------------------------------------------------------------
  // 2-species data
  //---------------------------------------------------------------------------

  // parameters for integrator
  p[1] = r;
  p[2] = K;
  p[3] = b;
  p[4] = h;
  p[5] = e;
  p[6] = d;

  for (j in 1:m){
    // integrate ODE
    Nsim = integrate_ode_rk45(odemodel, {N10sim[j],N20sim[j]}, t[1], t[2:n], p, rep_array(0.0,0), re
    // likelihood
    N1[j,1] ~ neg_binomial_2(N10sim[j]*f,tau[1]);
    N2[j,1] ~ neg_binomial_2(N20sim[j]*f,tau[2]);
    for (i in 2:n){
      N1[j,i] ~ neg_binomial_2(Nsim[i-1,1]*f,tau[1]);
      N2[j,i] ~ neg_binomial_2(Nsim[i-1,2]*f,tau[2]);
    }
  }

  //---------------------------------------------------------------------------
  // resource: control data
  //---------------------------------------------------------------------------
```

```
  for (j in 1:mc){
    // integrate ODE
    for(i in 2:n){
      NCsim[i-1,1] = K / ( 1+(K-N1C0sim[j])/N1C0sim[j] * exp(-r*(t[i]-t[1]))); // analytical solutio
    }
    // likelihood
    N1C[j,1] ~ neg_binomial_2(N1C0sim[j]*f,tau[1]);
    for (i in 2:n){
      N1C[j,i] ~ neg_binomial_2(NCsim[i-1,1]*f,tau[1]);
    }
  }

  //---------------------------------------------------------------------------
  // consumer: control data
  //---------------------------------------------------------------------------

  for (j in 1:mc){
    // integrate ODE
    for(i in 2:n){
      NCsim[i-1,1] = N2C0sim[j] * exp(-d*(t[i]-t[1])); // analytical solution
    }
    // likelihood
    N2C[j,1] ~ neg_binomial_2(N2C0sim[j]*f,tau[2]);
    for (i in 2:n){
      N2C[j,i] ~ neg_binomial_2(NCsim[i-1,1]*f,tau[2]);
    }
  }
}
'
```

##### 4.3.1 functions{ } block

The population growth equation (the ODE) is coded as a function. This function is later used for numerical integration.

Stan requires a specific format: The output is a real-valued array, and arguments time `t`, state `N` (contains prey and predator abundance), and model parameters `p` are provided. Additional arguments `x_r` and `x_i` for real- and integer-valued data have to be defined, but are not used here.

The rates of change `dNdt` are stored in an array of size 2 for prey and predator. It is computed using the 2d states `N` and parameters `p` and returned as the function's output.

```
real[] odemodel(real t, real[] N, real[] p, real[] x_r, int[] x_i){
  // p[1]=r, p[2]=K, p[3]=b, p[4]=h, p[5]=e, p[6]=d
  real dNdt[2];
  dNdt[1] = p[1]*N[1]*(1.0-N[1]/p[2])-p[3]*N[1]*N[2]/(1.0+p[3]*p[4]*N[1]);
  dNdt[2] = p[5]*p[3]*N[1]*N[2]/(1.0+p[3]*p[4]*N[1])-p[6]*N[2];
  return dNdt;
}
```

##### 4.3.2 data{ } block

All data have to be declared including data types and dimensions. Names have to be identical to the named list which was defined in R before.

```
int n; // observations
int m; // replicates
int mc; // replicates control
real t[n];
int N1[m,n];
```

```
int N2[m,n];
int N1C[mc,n];
int N2C[mc,n];
real f;
```

##### 4.3.3 parameters{ } block

Here, all free parameters are declared. This includes model parameters `r`,`K`,`b`,`h`,`e`,`d` and 2 parameters `tau` (for prey and predator observations, respectively) for the residual distribution (overdispersion of the negative binomial). Additionally, the initial states of each of the `m` modelled time series `N10sim[m]` and `N20sim[m]`, as well as the control time series `N1C0sim[mc]` and `N2C0sim[mc]`(defined as real-valued arrays), in the total volume are estimated, too. Its true states are unknown and observations are assumed to be measured with error.

```
real<lower=0> r;
real<lower=0> K;
real<lower=0> b;
real<lower=0> h;
real<lower=0, upper=1> e;
real<lower=0> d;
real<lower=0> tau[2];
real<lower=0> N10sim[m]; // individual initial values for replicates
real<lower=0> N20sim[m]; // individual initial values for replicates
real<lower=0> N1C0sim[mc]; // individual initial values for replicates
real<lower=0> N2C0sim[mc]; // individual initial values for replicates
```

##### 4.3.4 model{ } block

This block defines how the posterior computed, using prior distributions and the likelihood function. First, some intermediate variables for the ODE simulation are declared. `p` is an array which stores all 6 model parameters. `Nsim` will contain the output of the numerical integration with dimensions `n-1` (at all timepoints minus the first one), and the second dimension is number of states (2). `NCsim` will contain predictions for single species control time series, so only 1 state.

```
real p[6];
real Nsim[n-1,2]; // simulated values, matrix. dim1 = time without t0, dim2 = dim_ODE = 2
real NCsim[n-1,1]; // simulated values, matrix. dim1 = time without t0, dim2 = dim_ODE = 1
```

Then, priors are defined for the model parameters. Weakly informative priors are chosen for the model parameters `r` and `K`. The overdispersion parameter `tau` of the residuals is constrained >0, but arbitrary small values are possible. We choose a gamma prior that decreases towards zero, such that sampling at low values near the boundary doesn't cause convergence issues. We do the same for positively contraint model parameters `b`, `h` and `d`. Model parameter `e` is contrained between 0 and 1. The parameters for the initial values in the total volume are assigned weak priors, too.

```
// priors
r ~ lognormal(-2,1);
K ~ lognormal(9,1);
b ~ gamma(2,1);
h ~ gamma(2,1);
// e ~ gamma(2,1);
d ~ gamma(2,1);
tau ~ gamma(2,0.1);

N10sim ~ normal(0,2000);
N20sim ~ normal(0,2000);
N1C0sim ~ normal(0,2000);
N2C0sim ~ normal(0,2000);
```

After the priors are defined, predictions are computed. We start with the 2-species mixtures, and then single species time series for prey and predator (control data).

For the 2-species time series, predictions are computed via numerically integrating the ODE, which requires all model parameters are stored in one array `p`.

```
//---------------------------------------------------------------------------
// 2-species data
//---------------------------------------------------------------------------

// parameters for integrator
p[1] = r;
p[2] = K;
p[3] = b;
p[4] = h;
p[5] = e;
p[6] = d;
```

In a loop over all `m` time series replicates, numerical integration is performed. Each time series `j` gets its individual initial states `N10sim[j],N20sim[j]` for prey and predator. The integration routine `integrate_ode_rk45()` is must be called in a specific format. Arguments are the function `odemodel` as defined in the `functions{}` block, the 2 initial states are converted to an array using `{ }`, then initial time `t[1]` and output times `t[2:n]`, and model parameters `p`. For additional variables `x_r` and `x_i` (mandatory but not used here), empty arrays are generated.

After computing the predicted time series `Nsim`, it is confronted with the data: a negative binomial distribution defines likelihood values. Since the data was sampled in a fraction `f=0.1`, the distribution's mean is `Nsim[,]*f`, with some overdispersion `tau[1]` for the prey and `tau[2]` for the predator abundance.

Since predictions by `integrate_ode_rk45()` can only be generated for $t > t_1$, data `N1[j,1]` and `N1[j,i]` ($i > 1$) have to be treated separately. The first observation `N1[j,1]` is distributed around the prediction in $t_1$, which is equal to the free parameter `N10sim[j]*f` (estimated initial state). The remaining observations `N1[j,i]` ($i > 1$) are distributed around the predictions in $t_i$, respectively, which are equal to the output `Nsim[i-1,1]*f`. It is `i-1` instead of `i` here, because the output `Nsim` starts in $t_2$ and not in $t_1$.

Same applies for predator osbervations `N2[j,i]` and predictions `Nsim[i-1,2]*f`.

```
for (j in 1:m){
  // integrate ODE
  Nsim = integrate_ode_rk45(odemodel, {N10sim[j],N20sim[j]}, t[1], t[2:n], p, rep_array(0.0,0), re
  // likelihood
  N1[j,1] ~ neg_binomial_2(N10sim[j]*f,tau[1]);
  N2[j,1] ~ neg_binomial_2(N20sim[j]*f,tau[2]);
  for (i in 2:n){
    N1[j,i] ~ neg_binomial_2(Nsim[i-1,1]*f,tau[1]);
    N2[j,i] ~ neg_binomial_2(Nsim[i-1,2]*f,tau[2]);
  }
}
```

Next, we compute predictions for the single species prey time series (control data). They follow logistic growth, so we can compute them analytically, i.e. with a direct formula instead of numerical integration:

$$\widehat{N}_1(t) = \frac{K}{1 + \frac{K - N_{1,0}}{N_{1,0}} e^{-r(t-t_0)}}$$

This is generally faster than performing numerical integration. If no analytical formulation is available, numerical integration has to be performed by either using the same ODE function as above with initial values `{N1C0sim[j], 0}`, or by coding a separate function for the single-species growth model.

```
//---------------------------------------------------------------------------
// resource: control data
//---------------------------------------------------------------------------

for (j in 1:mc){
  // integrate ODE
```

```
    for(i in 2:n){
      NCsim[i-1,1] = K / ( 1+(K-N1C0sim[j])/N1C0sim[j] * exp(-r*(t[i]-t[1]))); // analytical solutio
    }
    // likelihood
    N1C[j,1] ~ neg_binomial_2(N1C0sim[j]*f,tau[1]);
    for (i in 2:n){
      N1C[j,i] ~ neg_binomial_2(NCsim[i-1,1]*f,tau[1]);
    }
  }
```

Finally, we do the same for the single species predator time series (control data). They follow exponential decline with an analytical solution

$$\widehat{N}_2(t) = N_{2,0}e^{-d(t-t_0)}$$

Again, if no analytical formulation is available, numerical integration has to be performed by either using the same ODE function as above with initial values `{0, N2C0sim[j]}`, or by coding a separate function for the single-species predator model.

```
  //---------------------------------------------------------------------------
  // consumer: control data
  //---------------------------------------------------------------------------

  for (j in 1:mc){
    // integrate ODE
    for(i in 2:n){
      NCsim[i-1,1] = N2C0sim[j] * exp(-d*(t[i]-t[1])); // analytical solution
    }
    // likelihood
    N2C[j,1] ~ neg_binomial_2(N2C0sim[j]*f,tau[2]);
    for (i in 2:n){
      N2C[j,i] ~ neg_binomial_2(NCsim[i-1,1]*f,tau[2]);
    }
  }
```

#### 4.4 Model fitting

After defining the model code, we specify some parameters for the MCMC model fitting. Usually, 3–5 MCMC chains are used. These can be run in parallel on multicore processors, when options are specified accordingly. Further, we set the total number of iterations and number of warmup iterations for the sampling. The total number of posterior MCMC samples is `chains*(iter-warmup)`.

```
# stan options
chains = 3
rstan_options(auto_write = TRUE)
options(mc.cores = chains)
iter   =  4000
warmup =  2000
```

Then starting values for all free parameters are provided as a list (in case of multiple chains: a list of lists). For the time series intial states `N10sim`, `N20sim`, `N1C0sim`, `N2C0sim`, data in `t[1]` scaled up from the sampled fraction `f` to the total volume are a natural choice. Remaining parameters are given guesses. Stan usually converges quickly towards the parameter region featuring a high probability density. But with ODEs involved, good initial guesses that do not cause numerical issues in ODE integration are often required.

```
# initial values for sampling
init=rep(list(list(r=0.1,
                   K=2e4,
                   b=0.0003,
```

```
                    h=0.5,
                    e=0.1,
                    d=0.1,
                    N10sim=data$N1[, 1]/data$f,
                    N20sim=data$N2[, 1]/data$f,
                    N1C0sim=data$N1C[, 1]/data$f,
                    N2C0sim=data$N2C[, 1]/data$f,
                    tau=c(10,10)
))
,chains)
```

Finally, the model is compiled and sampling is started.

```
stanmodel = stan_model(model_code=stanmodelcode)

fit = sampling(stanmodel,
                data=data,
                iter=iter,
                warmup=warmup,
                chains=chains,
                init=init
)
```

#### 4.5 Model diagnostics

Convergence of the MCMC chains is checked, e.g. by looking at estimated model parameters. We check the effective sample size `n_eff` and the Gelman-Rubin diagnostics `Rhat` (should be <1.01), indicating that chains have mixed.

```
print(fit, digits=3, pars=c("r","K","b","h","e","d"), probs=c(0.025, 0.5, 0.975))
```

```
## Inference for Stan model: bdbd516be3b0bb71d901713e64a11e42.
## 3 chains, each with iter=4000; warmup=2000; thin=1;
## post-warmup draws per chain=2000, total post-warmup draws=6000.
##
##        mean se_mean      sd      2.5%       50%     97.5% n_eff  Rhat
## r     0.103   0.000   0.004     0.095     0.103     0.112  2718 1.001
## K 19862.174   7.059 490.433 18921.325 19852.281 20840.856  4827 1.000
## b     0.000   0.000   0.000     0.000     0.000     0.000  2890 1.001
## h     0.513   0.000   0.023     0.468     0.512     0.559  2895 1.001
## e     0.046   0.000   0.003     0.041     0.046     0.052  2430 1.002
## d     0.048   0.000   0.002     0.045     0.048     0.052  3277 1.000
##
## Samples were drawn using NUTS(diag_e) at Mon Apr  4 16:19:43 2022.
## For each parameter, n_eff is a crude measure of effective sample size,
## and Rhat is the potential scale reduction factor on split chains (at
## convergence, Rhat=1).
```

Additionally, chains are inspected visually via trace- and density plots.

```
library("coda")
samples=As.mcmc.list(fit)
plot(samples[, c("r","K","b","h","e","d")])
```

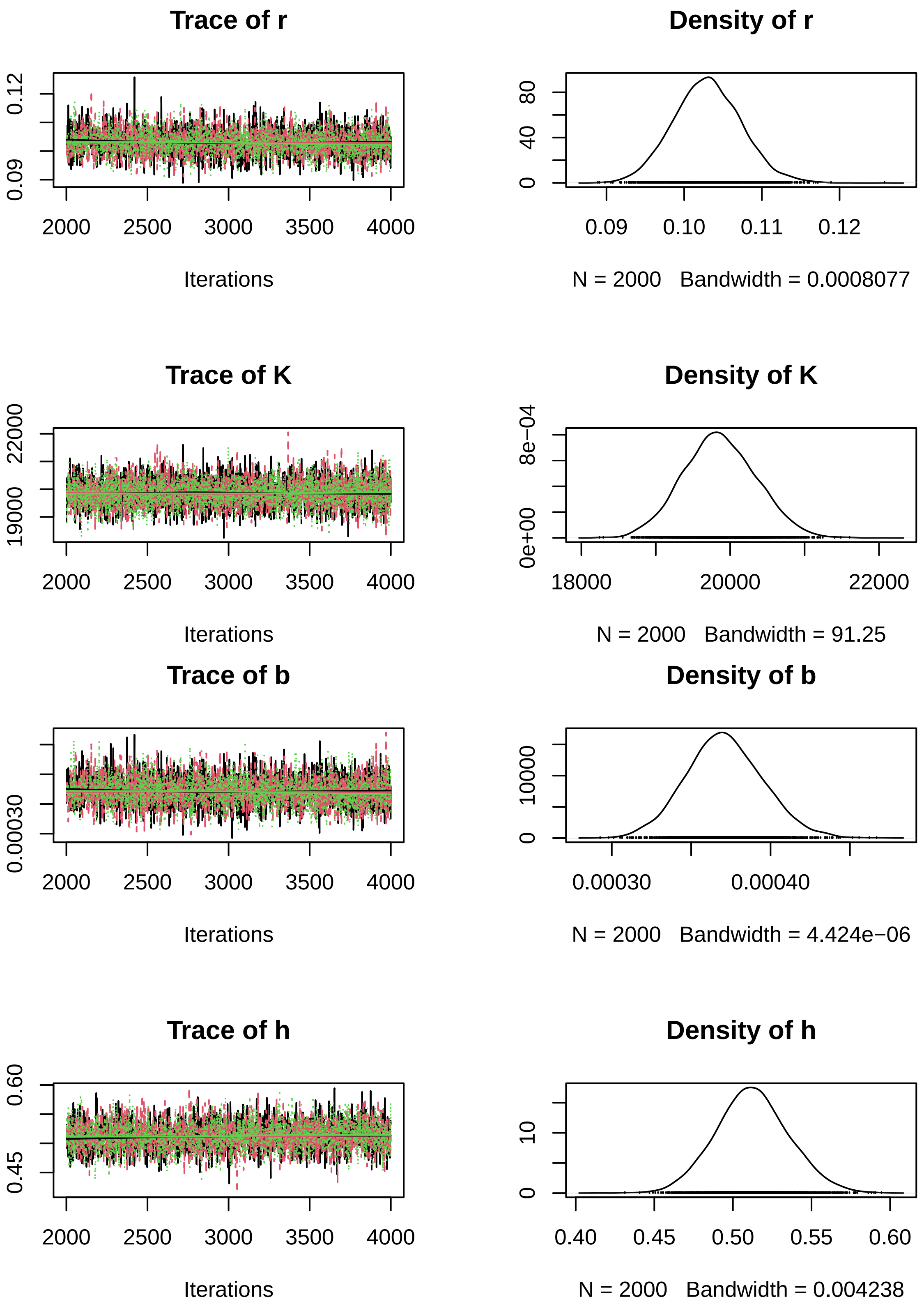

Trace of r
0.09
0.12
2000 2500 3000 3500 4000
Iterations
Density of r
0
40
80
0.09 0.10 0.11 0.12
N = 2000 Bandwidth = 0.0008077
Trace of K
19000
22000
2000 2500 3000 3500 4000
Iterations
Density of K
0e+00
8e−04
18000 20000 22000
N = 2000 Bandwidth = 91.25
Trace of b
0.00030
2000 2500 3000 3500 4000
Iterations
Density of b
0
10000
0.00030 0.00040
N = 2000 Bandwidth = 4.424e−06
Trace of h
0.45
0.60
2000 2500 3000 3500 4000
Iterations
Density of h
0
10
0.40 0.45 0.50 0.55 0.60
N = 2000 Bandwidth = 0.004238

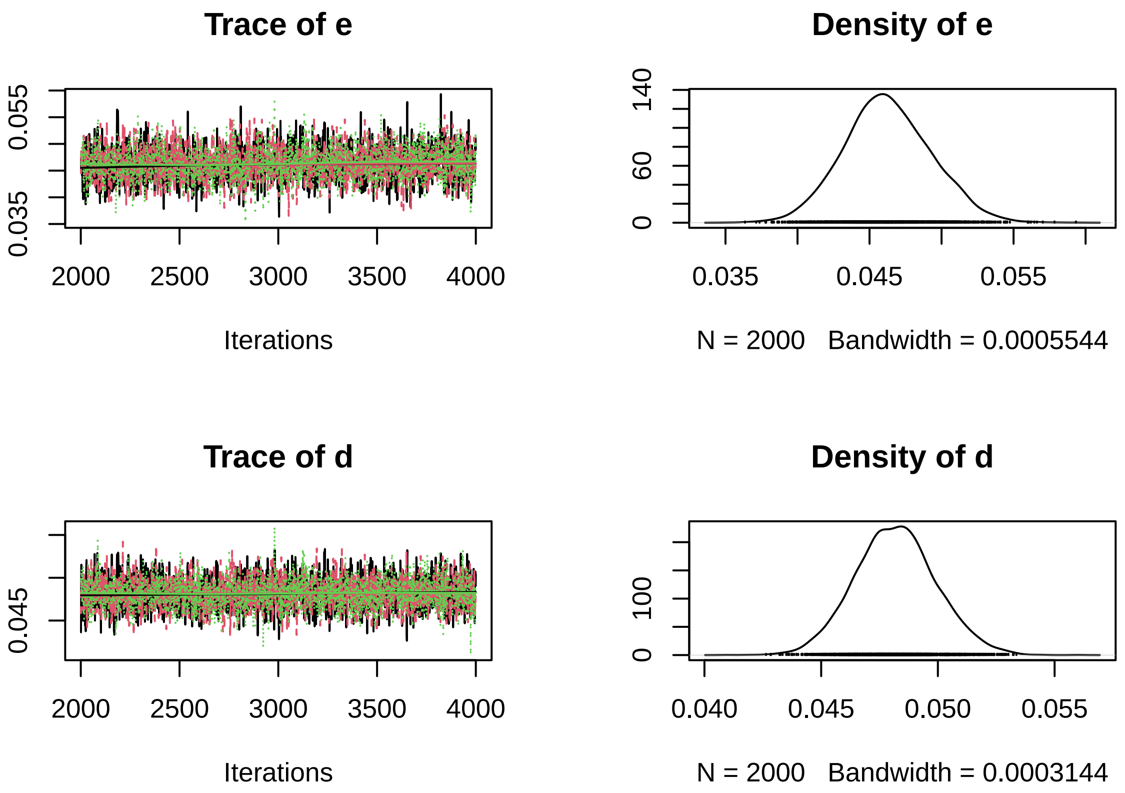


A pairs plot can detect correlations in model parameters. Almost perfect correlation would indicate non-identifiability.

```
pairs(fit, pars=c("r","K","b","h","e","d"))
```

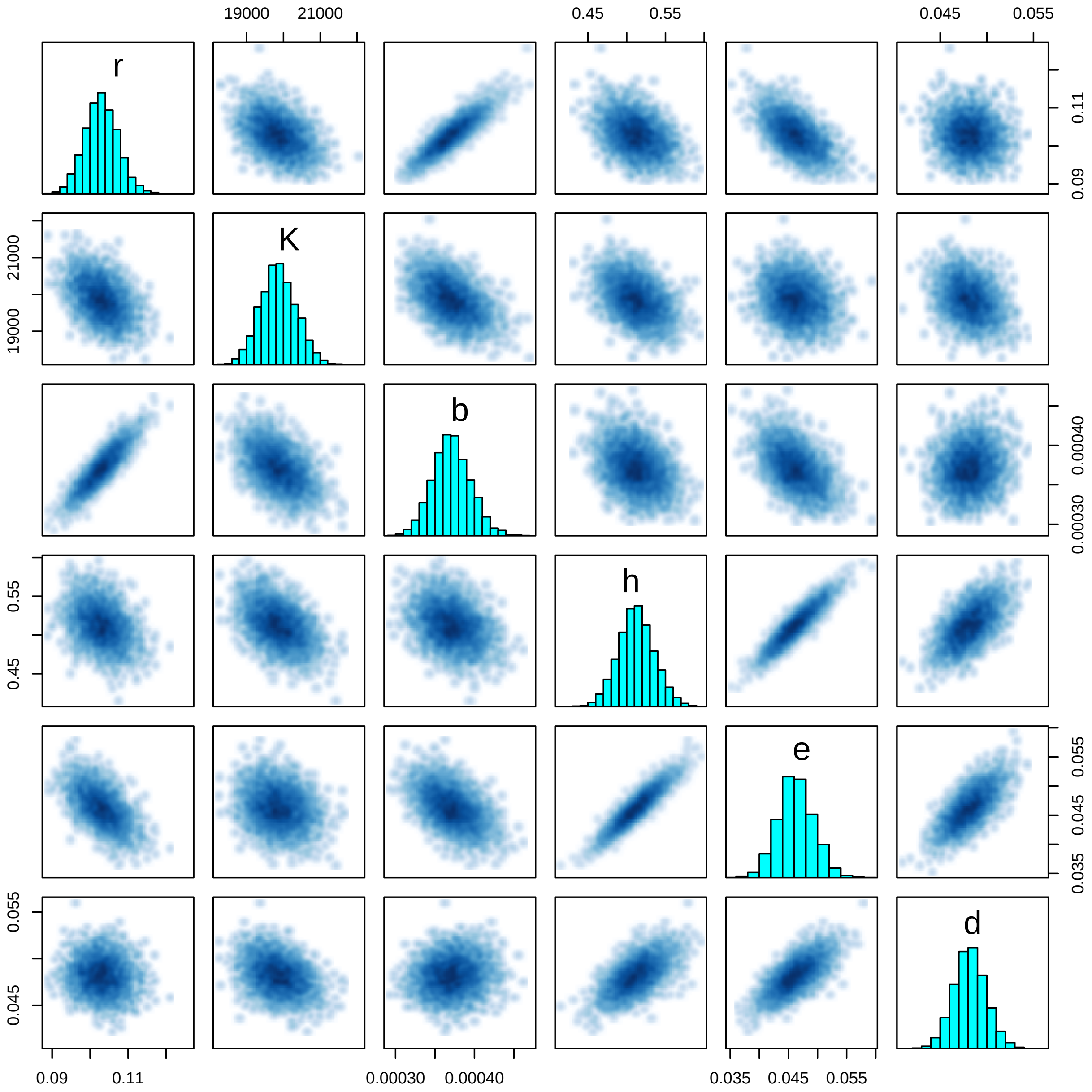


#### 4.6 Posterior predictions

Posterior predictions can generated from sampled posterior parameters. Alternatively, this can be done while sampling using `generated quantities{}` block in the model code. Here, we use the `deSolve` package for numerical integration to compute posterior predictions. First, we code an R function defining the population growth rate (ODE).

```
library("deSolve")
ode.model = function(t,N,p){
  dN1dt = p[1]*N[1]*(1.0-N[1]/p[2])-p[3]*N[1]*N[2]/(1.0+p[3]*p[4]*N[1])
  dN2dt = p[5]*p[3]*N[1]*N[2]/(1.0+p[3]*p[4]*N[1])-p[6]*N[2]
  return(list(c(dN1dt,dN2dt)))
}
```

Samples of the posterior distribution are converted to matrix. Each column contains all samples of a single parameter, while each row contains one sample from the multivariate posterior distribution.

```
post = as.matrix(fit)
# head(post)
```

`t.pred` includes times for which predictions are generated. The empty matrices `N1.pred` and `N2.pred` will store all predicted values for prey and predator: each row will contain one sample of a predicted timeseries.

```
n.post = 1000 # nrow(post)
t.pred = seq(from=min(data$t), to=max(data$t), by=1)
N1.pred = matrix(NA, nrow=n.post, ncol=length(t.pred))
N2.pred = matrix(NA, nrow=n.post, ncol=length(t.pred))
```

Predictions are generated for each time series replicate seperately (for-loop over `i`). For each sample from the posterior (row `j` in matrix `post[j, ]`), the bivariate trajectory is computed by numerically integrating the ODE with the model parameters, starting in initial state (`N10sim[i]`, `N20sim[i]`). Each column `k` of `N.pred[ ,k]` contains posterior predictions in `t.pred[k]`. We calculate quantiles of these predictions and plot them against the data.

Here, we only show the 2-species mixtures and omit the control time series.

Note that predicted time series differ only in their initial states (`N10sim[i]`, `N20sim[i]`), since model parameters are identical across time series replicates $i$.

```
par(mfrow=c(3,2), mar=c(3,3,0,0), oma=c(3,3,1,1))

for(i in 1:m){

  for(j in 1:n.post){
    Ns = as.data.frame(lsoda(y     = c(N=as.numeric(post[j, c(paste0("N10sim[",i,"]"),
                                                              paste0("N20sim[",i,"]"))])),
                             times = t.pred,
                             func  = ode.model,
                             parms = c(post[j, c("r","K","b","h","e","d")])
    ))[, 2:3]
    N1.pred[j, ]=Ns[, 1]
    N2.pred[j, ]=Ns[, 2]
  }

  plot(c(data$t,data$t), c(data$N1[i, ]/data$f,data$N2[i, ]/data$f),
       ylim=c(1e2,2e4), type="n", xlab="", ylab="", log="y")

  N1.pred.qs = apply(N1.pred, 2, function(x) quantile(x, probs=c(0.025,0.500,0.975), na.rm=TRUE))
  polygon(c(t.pred, rev(t.pred)), c(N1.pred.qs[1, ], rev(N1.pred.qs[3, ])),
          col = adjustcolor("red",alpha.f=0.25),  border = NA)
  lines(t.pred, N1.pred.qs[2, ], col="red", lwd=1.5)
  points(data$t, data$N1[i, ]/data$f)

  N2.pred.qs = apply(N2.pred, 2, function(x) quantile(x, probs=c(0.025,0.500,0.975), na.rm=TRUE))
  polygon(c(t.pred, rev(t.pred)), c(N2.pred.qs[1, ], rev(N2.pred.qs[3, ])),
          col = adjustcolor("blue",alpha.f=0.25),  border = NA)
  lines(t.pred, N2.pred.qs[2, ], col="blue", lwd=1.5)
  points(data$t, data$N2[i, ]/data$f)

  if(i==1) legend("topleft", c("res","con"), lwd=c(2,2), col=c("red","blue"))
}
mtext("Time [h]", side=1, outer=TRUE, line=1)
mtext("Abundance [ind], logscale", side=2, outer=TRUE, line=1)
```

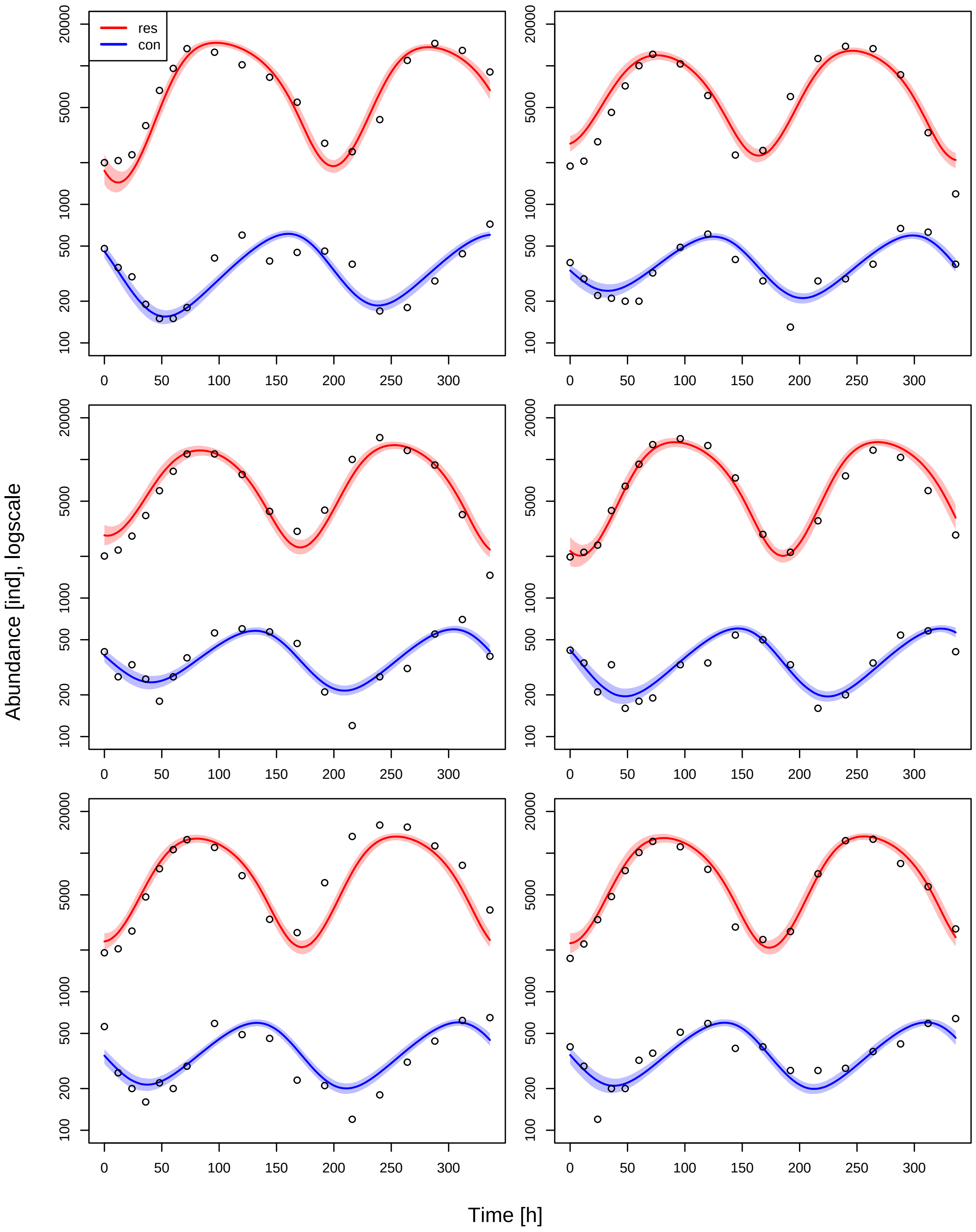
res
con
Abundance [ind], logscale
Time [h]
20000
5000
1000
500
200
100
0
50
100
150
200
250
300

### 5 Consumer resource - PROC model

#### 5.1 Model

The Rosenzweig-MacArthur predator-prey model with logistic growth and Type II functional response reads

$$\frac{dN_1}{dt} = r(1 - \frac{N_1}{K})N_1 - \frac{ahN_1}{1 + ahN_1}N_2$$
$$\frac{dN_2}{dt} = e\frac{ahN_1}{1 + ahN_1}N_2 - dN_2$$

Using one-step-ahead fitting (process error only), we will fit all datapoints $(N_{1,t}, N_{2,t})$ of $m$ 2-species timeseries replicates with predictions based on their previous observations $(N_{1,t-1}, N_{2,t-1})$. A joint set of model parameters $r, K, a, h, e, d$ ("complete pooling") is used.

Additionally, we fit trajectories for $m_c$ control (single-species) time series of prey ($N_2 = 0$, logistic growth)

$$\frac{dN_1}{dt} = r(1 - \frac{N_1}{K})N_1$$

and $m_c$ control time series of predator ($N_1 = 0$, exponential decline)

$$\frac{dN_2}{dt} = -dN_2$$

These control data will provide additional information on $r, K$ and $d$.

#### 5.2 Data preparation

The dataset `data_example_consumer_resource.csv` contains `m=6` timeseries with `n=18` observations of densities [ind] each, including their dates [h]. The fraction of the observed volume for sampling was `f=0.1`. Data preparation is identical to the OBS model manual.

```
rm(list=ls())
library("rstan")

data.in = read.csv("data_example_consumer_resource.csv", row.names=1)

head(data.in)
```

```
##        X1  X2  X3  X4  X5   X6   X7   X8   X9 X10 X11 X12  X13  X14  X15  X16
## time    0  12  24  36  48   60   72   96  120 144 168 192  216  240  264  288
## N1    200 207 228 370 664  958 1330 1253 1017 827 546 276  240  409 1094 1452
## N2    189 205 283 461 716 1002 1212 1032  610 227 245 600 1128 1383 1330  862
## N3    201 222 280 394 595  822 1097 1097  779 422 303 431 1001 1439 1161  912
## N4    198 214 241 428 641  924 1281 1413 1261 736 288 214  361  761 1168 1036
## N5    191 204 274 483 772 1060 1251 1100  688 333 267 612 1320 1595 1542 1128
##        X17 X18
## time   312 336
## N1    1291 903
## N2     329 119
## N3     400 146
## N4     596 285
## N5     817 389
```

```
n = 18
m = 6
mc = 3

times = as.numeric(data.in[1, ])
N1 = unname(as.matrix(data.in[2:(m+1), ]))
```

```
N2 = unname(as.matrix(data.in[(m+2):(2*m+1), ]))
N1C = unname(as.matrix(data.in[(2*m+2):(2*m+1+mc), ]))
N2C = unname(as.matrix(data.in[(2*m+2+mc):((2*m+1+2*mc)), ]))

data = list(n = n,
            m = m,
            mc = mc,
            t = times,
            N1 = N1,
            N2 = N2,
            N1C  = N1C,
            N2C  = N2C,
            f=0.1)

str(data)
```

```
## List of 9
##  $ n  : num 18
##  $ m  : num 6
##  $ mc : num 3
##  $ t  : num [1:18] 0 12 24 36 48 60 72 96 120 144 ...
##  $ N1 : int [1:6, 1:18] 200 189 201 198 191 174 207 205 222 214 ...
##  $ N2 : int [1:6, 1:18] 48 38 41 42 56 40 35 29 27 34 ...
##  $ N1C: int [1:3, 1:18] 40 51 51 96 144 117 259 370 411 637 ...
##  $ N2C: int [1:3, 1:18] 195 185 201 104 98 105 58 56 44 40 ...
##  $ f  : num 0.1
```

#### 5.3 Stan model

The Stan model is coded as a string, which is compiled later. A step by step walkthrough is given below.

```
stanmodelcode='
functions{
  real[] odemodel(real t, real[] N, real[] p, real[] x_r, int[] x_i){
    // p[1]=r, p[2]=K, p[3]=b, p[4]=h, p[5]=e, p[6]=d
    real dNdt[2];
    dNdt[1] = p[1]*N[1]*(1.0-N[1]/p[2])-p[3]*N[1]*N[2]/(1.0+p[3]*p[4]*N[1]);
    dNdt[2] = p[5]*p[3]*N[1]*N[2]/(1.0+p[3]*p[4]*N[1])-p[6]*N[2];
    return dNdt;
  }
}

data{
  int n; // observations
  int m; // replicates
  int mc; // replicates control
  real t[n];
  int N1[m,n];
  int N2[m,n];
  int N1C[mc,n];
  int N2C[mc,n];
  real f;
}

parameters{
  real<lower=0> r;
  real<lower=0> K;
  real<lower=0> b;
  real<lower=0> h;
```

```
  real<lower=0, upper=1> e;
  real<lower=0> d;
  real<lower=0> tau[2];
}

model{
  real p[6];
  real Nsim[1,2]; // simulated values, matrix. dim1 = 1 point in time, dim2 = dim_ODE = 2
  real NCsim[1,1]; // simulated values, matrix. dim1 = 1 point in time, dim2 = dim_ODE = 1

  // priors
  r ~ lognormal(-2,1);
  K ~ lognormal(9,1);
  b ~ gamma(2,1);
  h ~ gamma(2,1);
  e ~ uniform(0,1);
  d ~ gamma(2,1);
  tau ~ gamma(2,0.1);

  //---------------------------------------------------------------------------
  // 2-species data
  //---------------------------------------------------------------------------

  // parameters for integrator
  p[1] = r;
  p[2] = K;
  p[3] = b;
  p[4] = h;
  p[5] = e;
  p[6] = d;

  for(j in 1:m){
    for(i in 1:(n-1)){
      // integrate ODE
      Nsim = integrate_ode_rk45(odemodel, {N1[j,i]/f,N2[j,i]/f}, t[i], {t[i+1]}, p, rep_array(0.0,0)
      // likelihood
      if(N1[j,i]>0){
        N1[j,i+1] ~ neg_binomial_2(Nsim[1,1]*f,tau[1]);
      }
      if(N2[j,i]>0){
        N2[j,i+1] ~ neg_binomial_2(Nsim[1,2]*f,tau[2]);
      }
    }
  }

  //---------------------------------------------------------------------------
  // resource: control data
  //---------------------------------------------------------------------------

  for (j in 1:mc){
    for(i in 1:(n-1)){
      if(N1C[j,i]>0){
        // integrate ODE
        NCsim[1,1] = K/( 1.0+(K-N1C[j,i]/f)/(N1C[j,i]/f) * exp(-r*(t[i+1]-t[i]))); // analytical sol
        // likelihood
        N1C[j,i+1] ~ neg_binomial_2(NCsim[1,1]*f,tau[1]);
      }
    }
```

```
  }

  //---------------------------------------------------------------------------
  // consumer: control data
  //---------------------------------------------------------------------------

  for (j in 1:mc){
    for(i in 1:(n-1)){
      if(N2C[j,i]>0){
        // integrate ODE
        NCsim[1,1] = N2C[j,i]/f * exp(-d*(t[i+1]-t[i])); // analytical solution
        // likelihood
        N2C[j,i+1] ~ neg_binomial_2(NCsim[1,1]*f,tau[2]);
      }
    }
  }
}
'
```

##### 5.3.1 functions{ } block

This is identical to the OBS model manual.

```
real[] odemodel(real t, real[] N, real[] p, real[] x_r, int[] x_i){
  // p[1]=r, p[2]=K, p[3]=b, p[4]=h, p[5]=e, p[6]=d
  real dNdt[2];
  dNdt[1] = p[1]*N[1]*(1.0-N[1]/p[2])-p[3]*N[1]*N[2]/(1.0+p[3]*p[4]*N[1]);
  dNdt[2] = p[5]*p[3]*N[1]*N[2]/(1.0+p[3]*p[4]*N[1])-p[6]*N[2];
  return dNdt;
}
```

##### 5.3.2 data{ } block

This is identical to the OBS model manual.

```
int n; // observations
int m; // replicates
int mc; // replicates control
real t[n];
int N1[m,n];
int N2[m,n];
int N1C[mc,n];
int N2C[mc,n];
real f;
```

##### 5.3.3 parameters{ } block

Here, all free parameters are declared. This includes model parameters `r`,`K`,`b`,`h`,`e`,`d` and 2 parameters `tau` (for prey and predator observations, respectively) for the residual distribution (overdispersion of the negative binomial).

```
real<lower=0> r;
real<lower=0> K;
real<lower=0> b;
real<lower=0> h;
real<lower=0, upper=1> e;
real<lower=0> d;
real<lower=0> tau[2];
```

##### 5.3.4 model{ } block

This block defines how the posterior computed, using prior distributions and the likelihood function. First, some intermediate variables for the ODE simulation are declared. `p` is an array which stores all 6 model parameters. `Nsim` will contain the output of the numerical integration with dimensions `1` (predict only next state), and the second dimension is number of states (2). `NCsim` will contain predictions for single species control time series, so only 1 state.

```
real p[6];
real Nsim[1,2]; // simulated values, matrix. dim1 = 1 point in time, dim2 = dim_ODE = 2
real NCsim[1,1]; // simulated values, matrix. dim1 = 1 point in time, dim2 = dim_ODE = 1
```

Priors: This is identical to the OBS model manual.

```
// priors
r ~ lognormal(-2,1);
K ~ lognormal(9,1);
b ~ gamma(2,1);
h ~ gamma(2,1);
e ~ uniform(0,1);
d ~ gamma(2,1);
tau ~ gamma(2,0.1);
```

After the priors are defined, predictions are computed. We start with the 2-species mixtures, and then single species time series for prey and predator (control data).

For the 2-species time series, predictions are computed via numerically integrating the ODE, which requires all model parameters are stored in one array `p`.

```
//--------------------------------------------------------------------------
// 2-species data
//--------------------------------------------------------------------------

// parameters for integrator
p[1] = r;
p[2] = K;
p[3] = b;
p[4] = h;
p[5] = e;
p[6] = d;
```

In a loop over all `m` time series replicates and `n-1` timepoints, numerical integration is performed, to predict the next state (`N1[j,i+1]`,`N2[j,i+1]`) from the current state (`N1[j,i]`,`N2[j,i]`). Observed data is abundance in sampling fraction `f`, so the initial values are scaled up to the total volume (`N1[j,i]/f`,`N2[j,i]/f`) for numerical integration, and predictions are scaled down later to the sampling fraction (`Nsim[,]*f`) for calculating residuals. The integration routine `integrate_ode_rk45()` is must be called in a specific format. Arguments are the function `odemodel` as defined in the `functions{}` block. The initial state is converted to an array using `{ }`. Integration is performed from `t[i]` to `t[i+1]`. `p` contains the model parameters. For additional variables `x_r` and `x_i` (mandatory but not used here), empty arrays are generated.

After computing the predicted state `Nsim` (in `t[i+1]`), it is confronted with the data (`N1[j,i+1]`,`N2[j,i+1]`): : a negative binomial distribution defines likelihood values. Since the data was sampled in a fraction `f=0.1`, the distribution's mean is `Nsim[,]*f`, with some overdispersion `tau`.

The prey prediction is only added to the likelihood if `N1[j,i]>0`. `N1[j,i]=0` would predict a 0 in `t[i+1]` irrespective of the model parameters, so it wouldn't contribute any information to the likelihood function. Same applies for predator osbervations `N2[j,i]` and their predictions.

```
for(j in 1:m){
  for(i in 1:(n-1)){
    // integrate ODE
    Nsim = integrate_ode_rk45(odemodel, {N1[j,i]/f,N2[j,i]/f}, t[i], {t[i+1]}, p, rep_array(0.0,0)
    // likelihood
```

```
      if(N1[j,i]>0){
        N1[j,i+1] ~ neg_binomial_2(Nsim[1,1]*f,tau[1]);
      }
      if(N2[j,i]>0){
        N2[j,i+1] ~ neg_binomial_2(Nsim[1,2]*f,tau[2]);
      }
    }
  }
```

Next, we compute one-step-ahead predictions for the single species prey time series (control data). They follow logistic growth, so we can compute them analytically, i.e. with a direct formula instead of numerical integration:

$$\widehat{N}_1(t_{i+1}) = \frac{K}{1 + \frac{K-N_{1,i}}{N_{1,i}} e^{-r(t_{i+1}-t_i)}}$$

This is generally faster than performing numerical integration. If no analytical formulation is available, numerical integration has to be performed by either using the same ODE function as above with initial values `{N1C[j,i], 0}`, or by coding a separate function for the single-species growth model.

```
//---------------------------------------------------------------------------
// resource: control data
//---------------------------------------------------------------------------

for (j in 1:mc){
  for(i in 1:(n-1)){
    if(N1C[j,i]>0){
      // integrate ODE
      NCsim[1,1] = K/( 1.0+(K-N1C[j,i]/f)/(N1C[j,i]/f) * exp(-r*(t[i+1]-t[i]))); // analytical sol
      // likelihood
      N1C[j,i+1] ~ neg_binomial_2(NCsim[1,1]*f,tau[1]);
    }
  }
}
```

Finally, we do the same for the single species predator time series (control data). They follow exponential decline with an analytical solution

$$\widehat{N}_2(t_{i+1}) = N_{2,i} e^{-d(t_{i+1}-t_i)}$$

Again, if no analytical formulation is available, numerical integration has to be performed by either using the same ODE function as above with initial values `{0, N2C[j,i]}`, or by coding a separate function for the single-species predator model.

```
//---------------------------------------------------------------------------
// consumer: control data
//---------------------------------------------------------------------------

for (j in 1:mc){
  for(i in 1:(n-1)){
    if(N2C[j,i]>0){
      // integrate ODE
      NCsim[1,1] = N2C[j,i]/f * exp(-d*(t[i+1]-t[i])); // analytical solution
      // likelihood
      N2C[j,i+1] ~ neg_binomial_2(NCsim[1,1]*f,tau[2]);
    }
  }
}
```

#### 5.4 Model fitting

This is identical to the OBS model manual.

```
# stan options
chains = 3
rstan_options(auto_write = TRUE)
options(mc.cores = chains)
iter   =  4000
warmup =  2000
```

Then starting values for all free parameters are provided as a list (in case of multiple chains: a list of lists). Remaining parameters are given guesses. Stan usually converges quickly towards the parameter region featuring a high probability density. But with ODEs involved, good initial guesses that do not cause numerical issues in ODE integration are often required.

```
# initial values for sampling
init=rep(list(list(r=0.1,
                   K=2e4,
                   b=0.0003,
                   h=0.5,
                   e=0.1,
                   d=0.1,
                   tau=c(10,10)
))
,chains)
```

Finally, the model is compiled and sampling is started.

```
stanmodel = stan_model(model_code=stanmodelcode)

fit = sampling(stanmodel,
               data=data,
               iter=iter,
               warmup=warmup,
               chains=chains,
               init=init
)
```

#### 5.5 Model diagnostics

Convergence of the MCMC chains is checked, e.g. by looking at estimated model parameters. We check the effective sample size `n_eff` and the Gelman-Rubin diagnostics `Rhat` (should be <1.01), indicating that chains have mixed.

```
print(fit, digits=3, pars=c("r","K","b","h","e","d"), probs=c(0.025, 0.5, 0.975))
```

```
## Inference for Stan model: 79a585f5976930ce93327af35b64294e.
## 3 chains, each with iter=4000; warmup=2000; thin=1;
## post-warmup draws per chain=2000, total post-warmup draws=6000.
##
##        mean se_mean      sd      2.5%       50%     97.5% n_eff  Rhat
## r     0.091   0.000   0.003     0.085     0.091     0.098  3221 1.000
## K 19903.440   8.783 520.758 18946.143 19892.982 20961.752  3516 1.001
## b     0.000   0.000   0.000     0.000     0.000     0.000  2619 1.001
## h     0.555   0.001   0.041     0.475     0.554     0.636  2292 1.003
## e     0.052   0.000   0.005     0.043     0.052     0.062  2127 1.002
## d     0.047   0.000   0.004     0.040     0.047     0.054  2461 1.001
##
## Samples were drawn using NUTS(diag_e) at Sat Apr  2 08:49:03 2022.
## For each parameter, n_eff is a crude measure of effective sample size,
```

```
## and Rhat is the potential scale reduction factor on split chains (at
## convergence, Rhat=1).
```

Additionally, chains are inspected visually via trace- and density plots.

```
library("coda")
samples=As.mcmc.list(fit)
plot(samples[, c("r","K","b","h","e","d")])
```

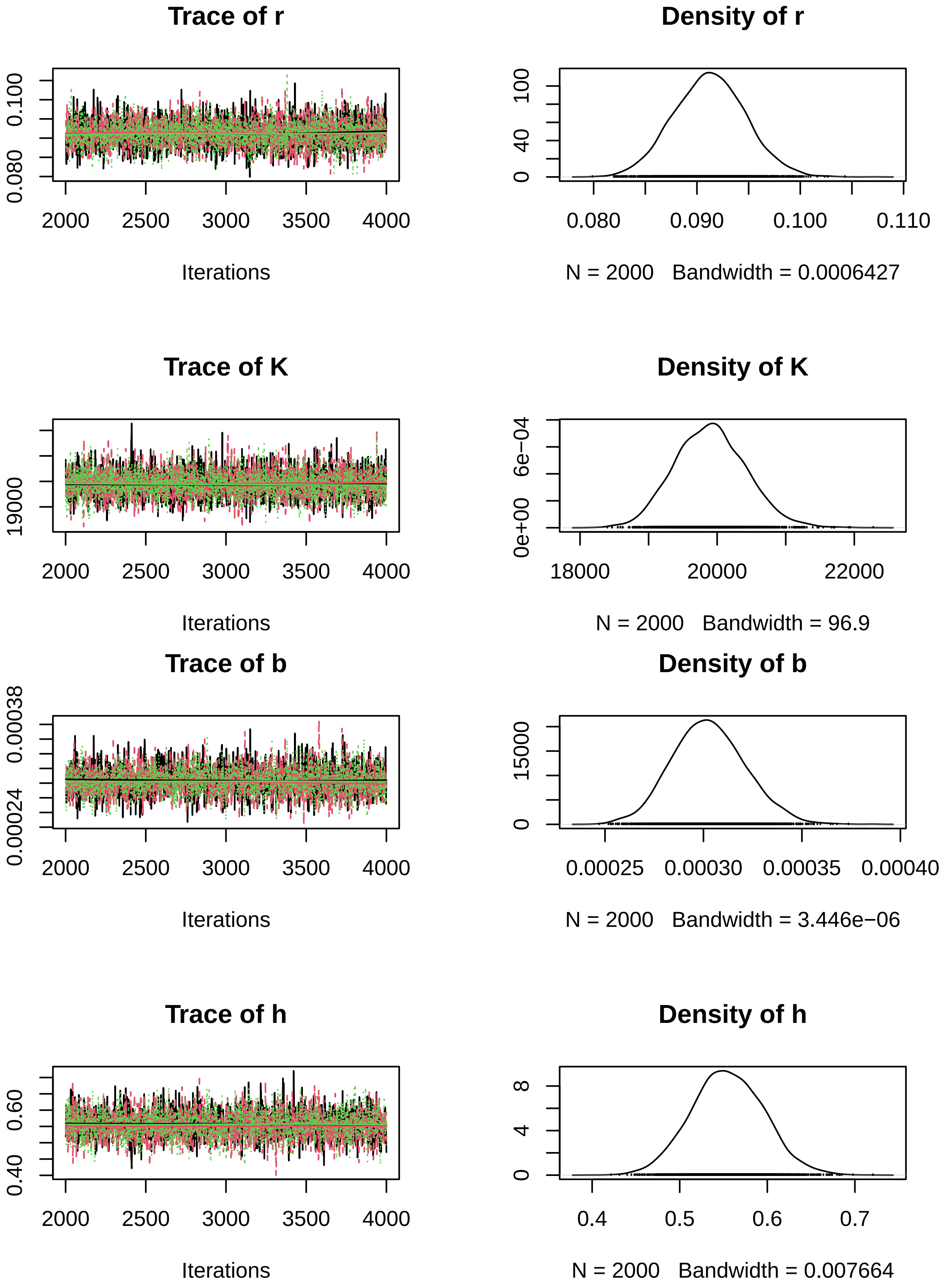

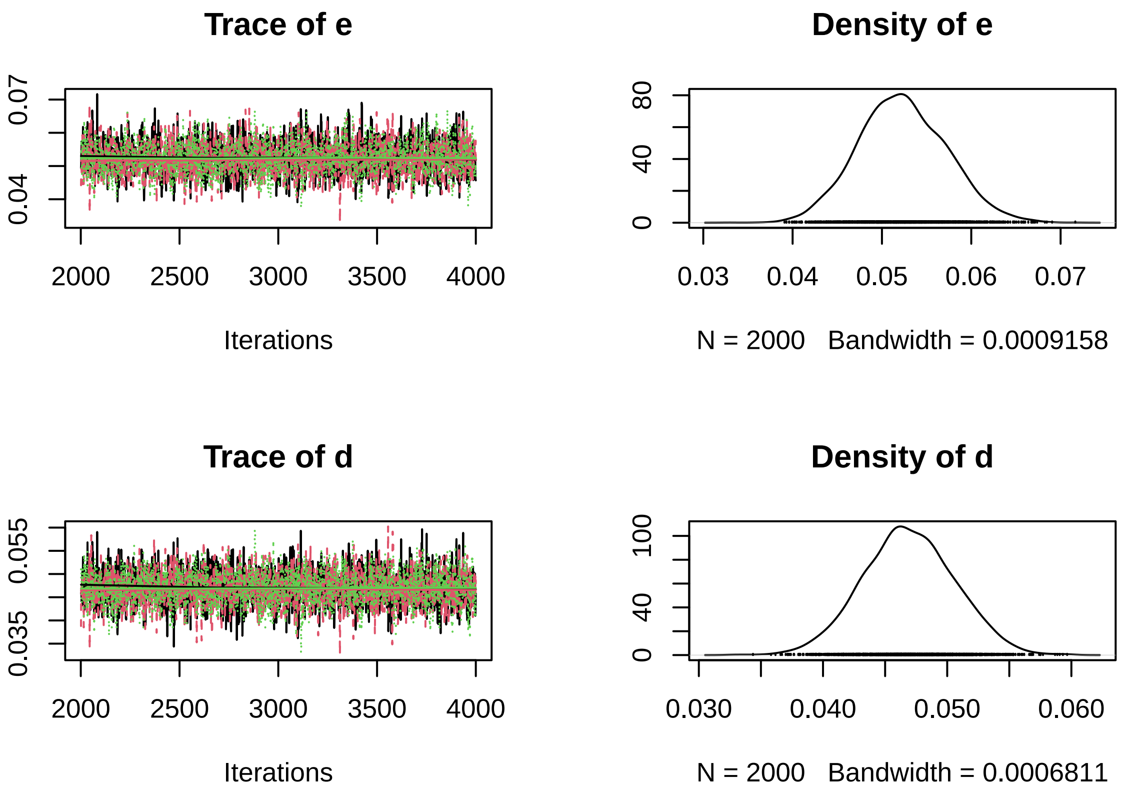


A pairs plot can detect correlations in model parameters. Almost perfect correlation would indicate non-identifiability.

```
pairs(fit, pars=c("r","K","b","h","e","d"))
```

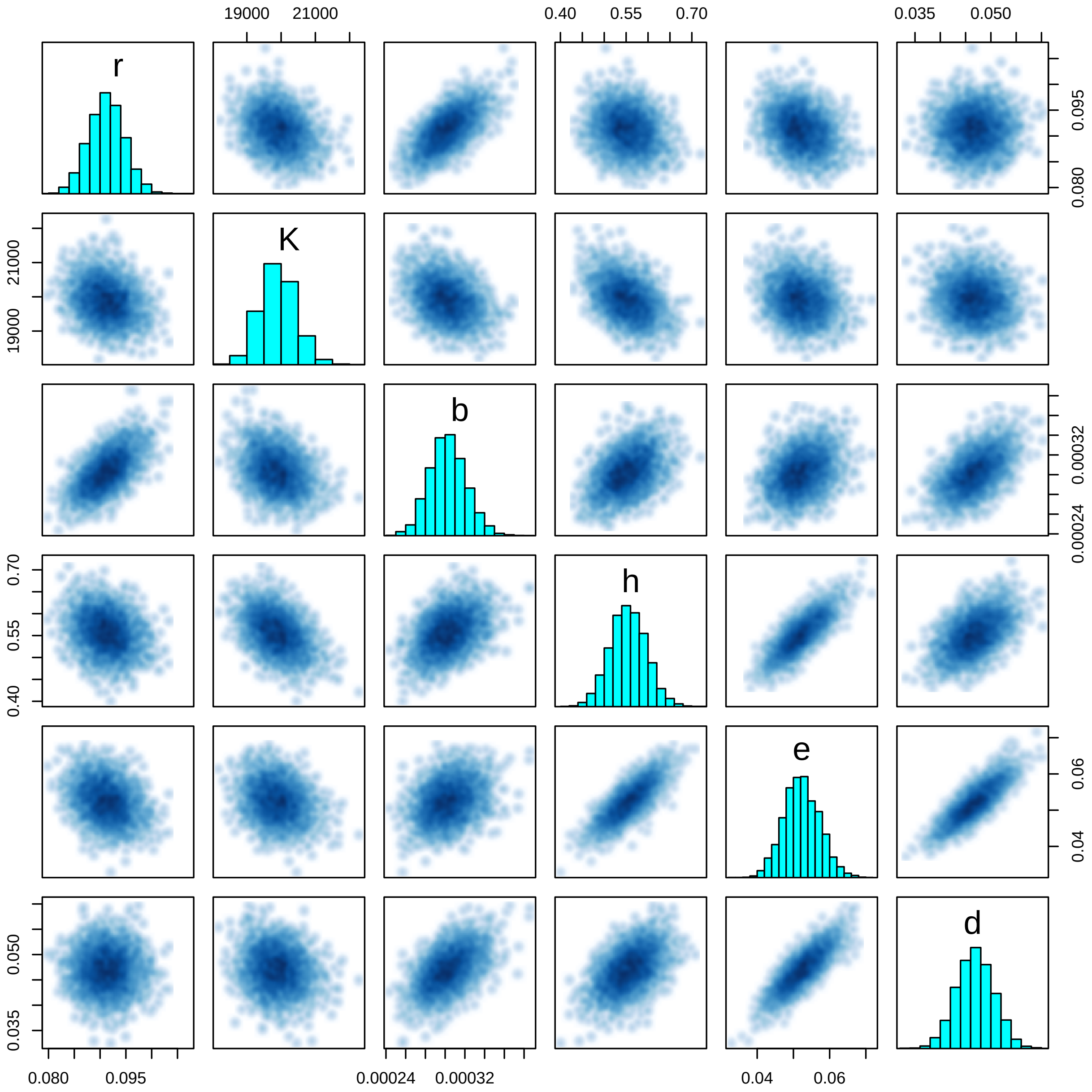


#### 5.6 Posterior predictions

Posterior predictions are generated one-step-ahead from sampled posterior parameters. Alternatively, this can be done while sampling using `generated quantities{}` block in the model code. Here, we use the `deSolve` package for numerical integration to compute posterior predictions. First, we code an R function defining the population growth rate (ODE).

```
library("deSolve")
ode.model = function(t,N,p){
  dN1dt = p[1]*N[1]*(1.0-N[1]/p[2])-p[3]*N[1]*N[2]/(1.0+p[3]*p[4]*N[1])
  dN2dt = p[5]*p[3]*N[1]*N[2]/(1.0+p[3]*p[4]*N[1])-p[6]*N[2]
  return(list(c(dN1dt,dN2dt)))
}
```

Samples of the posterior distribution are converted to matrix. Each column contains all samples of a single parameter, while each row contains one sample from the multivariate posterior distribution.

```
post = as.matrix(fit)
# head(post)
n.post = 1000 # nrow(post)
```

The empty matrices N1.pred and N2.pred(prey and predator) will store all predicted values: each row will contain one sample of a predicted timeseries. Predictions are computed for each time series replicate (for-loop over i) and each timepoint (for-loop over k) separately. For each sample from the posterior (row j in matrix post[j, ]), each prediction is computed by numerically integrating the ODE, starting in the last observation (data$N1[i,k]/data$f,data$N1[i,k]/data$f, scaled up to the total volume). Each column k of N1.pred[ ,k]/N2.pred[ ,k] contains posterior predictions in t[k+1]. We calculate quantiles of these predictions and plot them against the data.

Here, we only show the 2-species mixtures and omit the control time series.

Since these are one-step-ahead predictions from the last observation, we plot them pointwise in time, rather than a trajectory as in the OBS model.

```
par(mfrow=c(3,2), mar=c(3,3,0,0), oma=c(3,3,1,1))

for(i in 1:6){

  N1.pred = matrix(NA, nrow=n.post, ncol=length(data$t)-1)
  N2.pred = matrix(NA, nrow=n.post, ncol=length(data$t)-1)

  for(j in 1:n.post){
    for(k in 1:(length(data$t)-1)){
      Ns = as.data.frame(lsoda(y     = c( N = c(data$N1[i,k]/data$f,data$N2[i,k]/data$f) ),
                               times = data$t[c(k,k+1)],
                               func  = ode.model,
                               parms = c(post[j, c("r","K","b","h","e","d")])
      ))[2, 2:3]
      N1.pred[j,k] = as.numeric(Ns[1])
      N2.pred[j,k] = as.numeric(Ns[2])
    }
  }

  N1.pred.qs = apply(N1.pred, 2, function(x) quantile(x, probs=c(0.025,0.500,0.975), na.rm=TRUE))
  N2.pred.qs = apply(N2.pred, 2, function(x) quantile(x, probs=c(0.025,0.500,0.975), na.rm=TRUE))

  plot(c(data$t,data$t), c(data$N1[i, ]/data$f,data$N2[i, ]/data$f),
       ylim=c(1e2,2e4), type="n", xlab="", ylab="", log="y")

  for(k in 1:(n-1)){
    lines(c(data$t[k+1],data$t[k+1]), c(N1.pred.qs[1,k],N1.pred.qs[3,k]), col="red", lwd=1.5)
  }
  points((data$t[2:n]), N1.pred.qs[2, ], col="red", pch="-", cex=1.5)
  points((data$t[2:n]), (data$N1[i,2:n]/data$f), type="p")

  for(k in 1:(n-1)){
    lines(c(data$t[k+1],data$t[k+1]), c(N2.pred.qs[1,k],N2.pred.qs[3,k]), col="blue", lwd=1.5)
  }
  points(as.numeric(data$t[2:n]), N2.pred.qs[2, ], col="blue", pch="-", cex=1.5)
  points((data$t[2:n]), (data$N2[i,2:n]/data$f), type="p")

  if(i==1) legend("topleft", c("res","con"), lwd=c(2,2), col=c("red","blue"))
}
mtext("Time [h]", side=1, outer=TRUE, line=1)
mtext("Abundance [ind], logscale", side=2, outer=TRUE, line=1)
```

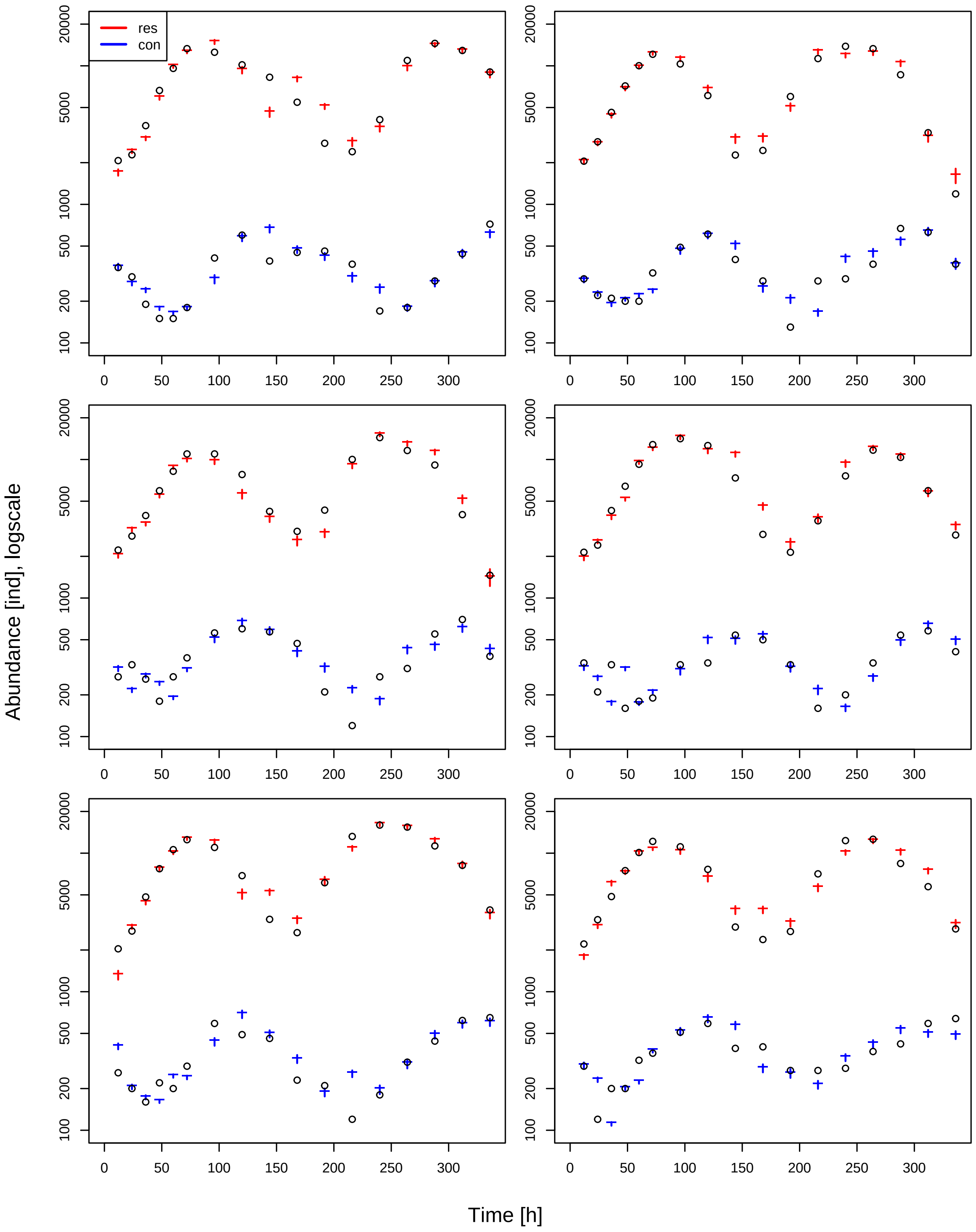
res
con
Abundance [ind], logscale
Time [h]
20000
5000
1000
500
200
100
0
50
100
150
200
250
300

### 6 Consumer resource - State-space model

#### 6.1 Model

The Rosenzweig-MacArthur predator-prey model with logistic growth and Type II functional response reads

$$\frac{dN_1}{dt} = r(1 - \frac{N_1}{K})N_1 - \frac{ahN_1}{1 + ahN_1}N_2$$
$$\frac{dN_2}{dt} = e\frac{ahN_1}{1 + ahN_1}N_2 - dN_2$$

Using a state-space model (including observation and process errors), we will fit all datapoints $(N_{1,t}, N_{2,t})$ of $m$ time series replicates with a joint set of parameters $r, K, a, h, e, d$ ("complete pooling"). For each observation $N_{1,i,j}, N_{2,i,j}$ in $t_i$ of time series $j$, underlying true states `Ntrue1[j,i]`, `Ntrue2[j,i]` are free parameters, which are estimated too.

Additionally, we fit time series for $m_c$ control (single-species) time series of prey ($N_2 = 0$, logistic growth)

$$\frac{dN_1}{dt} = r(1 - \frac{N_1}{K})N_1$$

and $m_c$ control time series of predator ($N_1 = 0$, exponential decline)

$$\frac{dN_2}{dt} = -dN_2$$

These control data will provide additional information on $r, K$ and $d$. Underlying true states `N1Ctrue[j,i]` for the prey and `N2Ctrue[j,i]` for the predator time series, respectively, are free parameters too.

#### 6.2 Data preparation

The dataset `data_example_consumer_resource.csv` contains `m=6` timeseries with `n=18` observations of densities [ind] each, including their dates [h]. The fraction of the observed volume for sampling was `f=0.1`. Data preparation is identical to the OBS model manual.

```
rm(list=ls())
library("rstan")

data.in = read.csv("data_example_consumer_resource.csv", row.names=1)

head(data.in)
```

```
##        X1  X2  X3  X4  X5   X6   X7   X8   X9 X10 X11 X12  X13  X14  X15  X16
## time    0  12  24  36  48   60   72   96  120 144 168 192  216  240  264  288
## N1    200 207 228 370 664  958 1330 1253 1017 827 546 276  240  409 1094 1452
## N2    189 205 283 461 716 1002 1212 1032  610 227 245 600 1128 1383 1330  862
## N3    201 222 280 394 595  822 1097 1097  779 422 303 431 1001 1439 1161  912
## N4    198 214 241 428 641  924 1281 1413 1261 736 288 214  361  761 1168 1036
## N5    191 204 274 483 772 1060 1251 1100  688 333 267 612 1320 1595 1542 1128
##       X17 X18
## time  312 336
## N1   1291 903
## N2    329 119
## N3    400 146
## N4    596 285
## N5    817 389
```

```
n = 18
m = 6
mc = 3
```

```
times = as.numeric(data.in[1, ])
N1 = unname(as.matrix(data.in[2:(m+1), ]))
N2 = unname(as.matrix(data.in[(m+2):(2*m+1), ]))
N1C = unname(as.matrix(data.in[(2*m+2):(2*m+1+mc), ]))
N2C = unname(as.matrix(data.in[(2*m+2+mc):((2*m+1+2*mc)), ]))

data = list(n = n,
            m = m,
            mc = mc,
            t = times,
            N1 = N1,
            N2 = N2,
            N1C  = N1C,
            N2C  = N2C,
            f = 0.1)

str(data)
```

```
## List of 9
##  $ n  : num 18
##  $ m  : num 6
##  $ mc : num 3
##  $ t  : num [1:18] 0 12 24 36 48 60 72 96 120 144 ...
##  $ N1 : int [1:6, 1:18] 200 189 201 198 191 174 207 205 222 214 ...
##  $ N2 : int [1:6, 1:18] 48 38 41 42 56 40 35 29 27 34 ...
##  $ N1C: int [1:3, 1:18] 40 51 51 96 144 117 259 370 411 637 ...
##  $ N2C: int [1:3, 1:18] 195 185 201 104 98 105 58 56 44 40 ...
##  $ f  : num 0.1
```

#### 6.3 Stan model

The Stan model is coded as a string, which is compiled later. A step by step walkthrough is given below.

```
stanmodelcode='
functions{
  real[] odemodel(real t, real[] N, real[] p, real[] x_r, int[] x_i){
    // p[1]=r, p[2]=K, p[3]=b, p[4]=h, p[5]=e, p[6]=d
    real dNdt[2];
    dNdt[1] = p[1]*N[1]*(1.0-N[1]/p[2])-p[3]*N[1]*N[2]/(1.0+p[3]*p[4]*N[1]);
    dNdt[2] = p[5]*p[3]*N[1]*N[2]/(1.0+p[3]*p[4]*N[1])-p[6]*N[2];
    return dNdt;
  }
}

data{
  int n; // observations
  int m; // replicates
  int mc; // replicates control
  real t[n];
  int N1[m,n];
  int N2[m,n];
  int N1C[mc,n];
  int N2C[mc,n];
  real f;
}

parameters{
  real<lower=0> r;
  real<lower=0> K;
```

```
  real<lower=0> b;
  real<lower=0> h;
  real<lower=0, upper=1> e;
  real<lower=0> d;
  real<lower=0> sigma[2];
  real<lower=0> tau[2];
  real<lower=0> N1true[m,n]; // states in times t
  real<lower=0> N2true[m,n]; // states in times t
  real<lower=0> N1Ctrue[mc,n]; // states in times t
  real<lower=0> N2Ctrue[mc,10]; // states in times t
}

model{
  real p[6];
  real Nsim[1,2]; // simulated values, matrix. dim1 = 1 point in time, dim2 = dim_ODE = 2
  real NCsim[1,1]; // simulated values, matrix. dim1 = 1 point in time, dim2 = dim_ODE = 1

  // priors
  r ~ lognormal(-2,1);
  K ~ lognormal(9,1);
  b ~ gamma(2,1);
  h ~ gamma(2,1);
  e ~ uniform(0,1);
  d ~ gamma(2,1);
  sigma ~ gamma(2,1);
  tau ~ gamma(2,0.1);

  for(i in 1:m){
    N1true[i,1] ~ normal(0,2000);
    N2true[i,1] ~ normal(0,2000);
  }
  for(i in 1:mc){
    N1Ctrue[i,1] ~ normal(0,2000);
    N2Ctrue[i,1] ~ normal(0,2000);
  }

  //---------------------------------------------------------------------------
  // 2-species data
  //---------------------------------------------------------------------------

  // parameters for integrator
  p[1] = r;
  p[2] = K;
  p[3] = b;
  p[4] = h;
  p[5] = e;
  p[6] = d;

  // process equations
  for(j in 1:m){
    for(i in 1:(n-1)){
      // integrate ODE
      Nsim = integrate_ode_rk45(odemodel, {N1true[j,i],N2true[j,i]}, t[i], {t[i+1]}, p, rep_array(0.
      // process error
      N1true[j,i+1] ~ lognormal(log(Nsim[1,1]), sigma[1]);
      N2true[j,i+1] ~ lognormal(log(Nsim[1,2]), sigma[2]);
    }
  }
```

```
  // observation equations
  for(j in 1:m){
    for(i in 1:n){
      N1[j,i] ~ neg_binomial_2(N1true[j,i]*f, tau[1]);
      N2[j,i] ~ neg_binomial_2(N2true[j,i]*f, tau[2]);
    }
  }

  //----------------------------------------------------------------------------
  // resource: control data
  //----------------------------------------------------------------------------

  // process equations
  for(j in 1:mc){
    for(i in 1:(n-1)){
      // integrate ODE
      NCsim[1,1] = K/( 1.0+(K-N1Ctrue[j,i])/N1Ctrue[j,i] * exp(-r*(t[i+1]-t[i]))); // analytical sol
      // process error
      N1Ctrue[j,i+1] ~ lognormal(log(NCsim[1,1]), sigma[1]);
    }
  }

  // observation equations
  for(j in 1:mc){
    for(i in 1:n){
      N1C[j,i] ~ neg_binomial_2(N1Ctrue[j,i]*f, tau[1]);
    }
  }

  //----------------------------------------------------------------------------
  // consumer: control data
  //----------------------------------------------------------------------------

  // process equations
  for(j in 1:mc){
    for(i in 1:(10-1)){ // only first 10 observations
      // integrate ODE
      NCsim[1,1] = N2Ctrue[j,i] * exp(-d*(t[i+1]-t[i])); // analytical solution
      // process error
      N2Ctrue[j,i+1] ~ lognormal(log(NCsim[1,1]), sigma[2]);
    }

  // observation equations
  for(j in 1:mc){
    for(i in 1:10){ // only first 10 observations
      N2C[j,i] ~ neg_binomial_2(N2Ctrue[j,i]*f, tau[2]); // sdev_obs now overdispersion parameter
    }
  }

}
'
```

##### 6.3.1 functions{ } block

This is identical to the OBS model manual.

```
  real[] odemodel(real t, real[] N, real[] p, real[] x_r, int[] x_i){
    // p[1]=r, p[2]=K, p[3]=b, p[4]=h, p[5]=e, p[6]=d
```

```
    real dNdt[2];
    dNdt[1] = p[1]*N[1]*(1.0-N[1]/p[2])-p[3]*N[1]*N[2]/(1.0+p[3]*p[4]*N[1]);
    dNdt[2] = p[5]*p[3]*N[1]*N[2]/(1.0+p[3]*p[4]*N[1])-p[6]*N[2];
    return dNdt;
  }
```

##### 6.3.2 data{ } block

This is identical to the OBS model manual.

```
int n; // observations
int m; // replicates
int mc; // replicates control
real t[n];
int N1[m,n];
int N2[m,n];
int N1C[mc,n];
int N2C[mc,n];
real f;
```

##### 6.3.3 parameters{ } block

Here, all free parameters are declared. This includes model parameters `r`,`K`,`b`,`h`,`e`,`d`, 2 parameters `sigma` for the process residual distribution (scale parameter of lognormal) and `tau` for the observation residual distribution (overdispersion of the negative binomial). Additionally, for all `m x n` observations, the underlying true states `Ntrue1` (prey) and `Ntrue2` (predator) are estimated. Same for control timeseries `N2Ctrue`, `N2Ctrue`. All parameters are required to be positive.

For the control data of the predator, we choose to use only the first 10 observations. The observed populations are extinct at that point. Predicting true states `N2Ctrue[i,j]` after that (`i>10`) would drive these estimates arbitrarily close to zero and potentially lead to numerical issues.

```
real<lower=0> r;
real<lower=0> K;
real<lower=0> b;
real<lower=0> h;
real<lower=0, upper=1> e;
real<lower=0> d;
real<lower=0> sigma[2];
real<lower=0> tau[2];
real<lower=0> N1true[m,n]; // states in times t
real<lower=0> N2true[m,n]; // states in times t
real<lower=0> N1Ctrue[mc,n]; // states in times t
real<lower=0> N2Ctrue[mc,10]; // states in times t
```

##### 6.3.4 model{ } block

This block defines how the posterior computed, using prior distributions and the likelihood function. First, some intermediate variables for the ODE simulation are declared. `p` is an array which stores all 6 model parameters. `Nsim` will contain the output of the numerical integration with dimensions `1` (predict only next state), and the second dimension is number of states (2). `NCsim` will contain predictions for single species control time series, so only 1 state.

```
real p[6];
real Nsim[1,2]; // simulated values, matrix. dim1 = 1 point in time, dim2 = dim_ODE = 2
real NCsim[1,1]; // simulated values, matrix. dim1 = 1 point in time, dim2 = dim_ODE = 1
```

Priors: This is identical to the OBS and PROC model manual, initial densities parameters `N...true[i,1]` are assigned a weak prior, too.

```
// priors
r ~ lognormal(-2,1);
```

```
K ~ lognormal(9,1);
b ~ gamma(2,1);
h ~ gamma(2,1);
e ~ uniform(0,1);
d ~ gamma(2,1);
sigma ~ gamma(2,1);
tau ~ gamma(2,0.1);

for(i in 1:m){
  N1true[i,1] ~ normal(0,2000);
  N2true[i,1] ~ normal(0,2000);
}
for(i in 1:mc){
  N1Ctrue[i,1] ~ normal(0,2000);
  N2Ctrue[i,1] ~ normal(0,2000);
}
```

After the priors are defined, predictions are computed. We start with the 2-species mixtures, and then single species time series for prey and predator (control data).

For the 2-species time series, predictions are computed via numerically integrating the ODE, which requires all model parameters are stored in one array `p`.

```
//-----------------------------------------------------------------------
// 2-species data
//-----------------------------------------------------------------------

// parameters for integrator
p[1] = r;
p[2] = K;
p[3] = b;
p[4] = h;
p[5] = e;
p[6] = d;
```

The likelihood consists of two parts: the process equations and the observation equations.

In a first loop over all `m` time series replicates and `n-1` timepoints, numerical integration is performed, to predict the next true state (`N1true[j,i+1]`,`N2true[j,i+1]`) from the current true state (`N1true[j,i]`,`N2true[j,i]`). The integration routine `integrate_ode_rk45()` is must be called in a specific format. Arguments are the function `odemodel` as defined in the `functions{}` block. The initial state is converted to an array using `{ }`. Integration is performed from `t[i]` to `t[i+1]`. `p` contains the model parameters. For additional variables `x_r` and `x_i` (mandatory but not used here), empty arrays are generated.

After computing the predicted state `Nsim` (in `t[i+1]`), it is confronted with estimate of (`N1true[j,i+1]`,`N2true[j,i+1]`): we assume a lognormal distribution, `sigma` denoting the process errors.

```
// process equations
for(j in 1:m){
  for(i in 1:(n-1)){
    // integrate ODE
    Nsim = integrate_ode_rk45(odemodel, {N1true[j,i],N2true[j,i]}, t[i], {t[i+1]}, p, rep_array(0.
    // process error
    N1true[j,i+1] ~ lognormal(log(Nsim[1,1]), sigma[1]);
    N2true[j,i+1] ~ lognormal(log(Nsim[1,2]), sigma[2]);
  }
}
```

Second, the true states estimates (`N1true[j,i]`, `N2true[j,i]`) are confronted with the data (`N1[j,i]`,`N2[j,i]`): a negative binomial distribution defines likelihood values. Since the data was sampled in a fraction `f=0.1`, the distribution's mean is `N..true[,]*f`, with some overdispersion `tau`.

```
// observation equations
for(j in 1:m){
  for(i in 1:n){
    N1[j,i] ~ neg_binomial_2(N1true[j,i], tau[1]);
    N2[j,i] ~ neg_binomial_2(N2true[j,i], tau[2]);
  }
}
```

Next, we compute one-step-ahead predictions for the single species prey time series (control data). They follow logistic growth, so we can compute them analytically, i.e. with a direct formula instead of numerical integration:

$$\widehat{N}_1(t_{i+1}) = \frac{K}{1 + \frac{K - N_{1,i}}{N_{1,i}} e^{-r(t_{i+1} - t_i)}}$$

This is generally faster than performing numerical integration. If no analytical formulation is available, numerical integration has to be performed by either using the same ODE function as above with initial values `{N1C[j,i], 0}`, or by coding a separate function for the single-species growth model.

Again, we have to code both process equations and observation equations.

```
//-------------------------------------------------------------------------
// resource: control data
//-------------------------------------------------------------------------

// process equations
for(j in 1:mc){
  for(i in 1:(n-1)){
    // integrate ODE
    NCsim[1,1] = K/( 1.0+(K-N1Ctrue[j,i])/N1Ctrue[j,i] * exp(-r*(t[i+1]-t[i]))); // analytical sol
    // process error
    N1Ctrue[j,i+1] ~ lognormal(log(NCsim[1,1]), sigma[1]);
  }
}

// observation equations
for(j in 1:mc){
  for(i in 1:n){
    N1C[j,i] ~ neg_binomial_2(N1Ctrue[j,i]*f, tau[1]);
  }
}
```

Finally, we do the same for the single species predator time series (control data). They follow exponential decline with an analytical solution

$$\widehat{N}_2(t_{i+1}) = N_{2,i} e^{-d(t_{i+1} - t_i)}$$

Again, if no analytical formulation is available, numerical integration has to be performed by either using the same ODE function as above with initial values `{0, N2C[j,i]}`, or by coding a separate function for the single-species predator model.

As described above, only the first 10 observations are considered (observed populations extinct after that).

Again, we have to code both process equations and observation equations.

```
//-------------------------------------------------------------------------
// consumer: control data
//-------------------------------------------------------------------------

// process equations
```

```
  for(j in 1:mc){
    for(i in 1:(10-1)){ // only first 10 observations
      // integrate ODE
      NCsim[1,1] = N2Ctrue[j,i] * exp(-d*(t[i+1]-t[i])); // analytical solution
      // process error
      N2Ctrue[j,i+1] ~ lognormal(log(NCsim[1,1]), sigma[2]);
    }

  // observation equations
  for(j in 1:mc){
    for(i in 1:10){ // only first 10 observations
      N2C[j,i] ~ neg_binomial_2(N2Ctrue[j,i]*f, tau[2]);
    }
  }
```

#### 6.4 Model fitting

This is identical to the OBS model manual.

```
# stan options
chains = 3
rstan_options(auto_write = TRUE)
options(mc.cores = chains)
iter   =  4000
warmup =  2000
```

Then starting values for all free parameters are provided as a list (in case of multiple chains: a list of lists). For the time series true states estimates `Ntrue`, observed data scaled up from the sampled fraction `f` to the total volume are a natural choice (we add 1 in case of zeros in the data to ensure positivity). Remaining parameters are given guesses. Stan usually converges quickly towards the parameter region featuring a high probability density. But with ODEs involved, good initial guesses that do not cause numerical issues in ODE integration are often required.

```
# initial values for sampling
init=rep(list(list(r=0.1,
                   K=2e4,
                   b=0.0003,
                   h=0.5,
                   e=0.1,
                   d=0.1,
                   sigma=c(0.1,0.1),
                   tau=c(10,10),
                   N1true=data$N1+1,
                   N2true=data$N2+1,
                   N1Ctrue=data$N1C+1,
                   N2Ctrue=data$N2C[, 1:10]+1
))
,chains)
```

Finally, the model is compiled and sampling is started. For this model and this dataset, some Stan control parameters are adapted to avoid divergent transitions while sampling. See https://mc-stan.org/misc/warnings.html for more information.

```
stanmodel = stan_model(model_code=stanmodelcode)

fit = sampling(stanmodel,
               data=data,
               iter=iter,
               warmup=warmup,
               chains=chains,
```

```
            init=init,
            control=list(adapt_delta=0.99, max_treedepth=12)
)
```

#### 6.5 Model diagnostics

Convergence of the MCMC chains is checked, e.g. by looking at estimated model parameters. We check the effective sample size `n_eff` and the Gelman-Rubin diagnostics `Rhat` (should be <1.01), indicating that chains have mixed.

```
print(fit, digits=3, pars=c("r","K","b","h","e","d"), probs=c(0.025, 0.5, 0.975))
```

```
## Inference for Stan model: 0f701afc0aa8a066f915085ec1770ad2.
## 3 chains, each with iter=4000; warmup=2000; thin=1;
## post-warmup draws per chain=2000, total post-warmup draws=6000.
##
##        mean se_mean      sd      2.5%       50%     97.5% n_eff  Rhat
## r     0.098   0.000   0.002     0.094     0.098     0.102  2076 1.003
## K 20082.003   6.223 210.055 19663.801 20079.646 20499.129  1139 1.003
## b     0.000   0.000   0.000     0.000     0.000     0.000  2026 1.001
## h     0.509   0.001   0.025     0.459     0.509     0.559  1924 1.001
## e     0.048   0.000   0.003     0.042     0.048     0.054  1827 1.001
## d     0.047   0.000   0.002     0.043     0.047     0.051  2793 1.001
##
## Samples were drawn using NUTS(diag_e) at Thu Apr  7 23:20:48 2022.
## For each parameter, n_eff is a crude measure of effective sample size,
## and Rhat is the potential scale reduction factor on split chains (at
## convergence, Rhat=1).
```

Additionally, chains are inspected visually via trace- and density plots.

```
library("coda")
samples=As.mcmc.list(fit)
plot(samples[, c("r","K","b","h","e","d")])
```

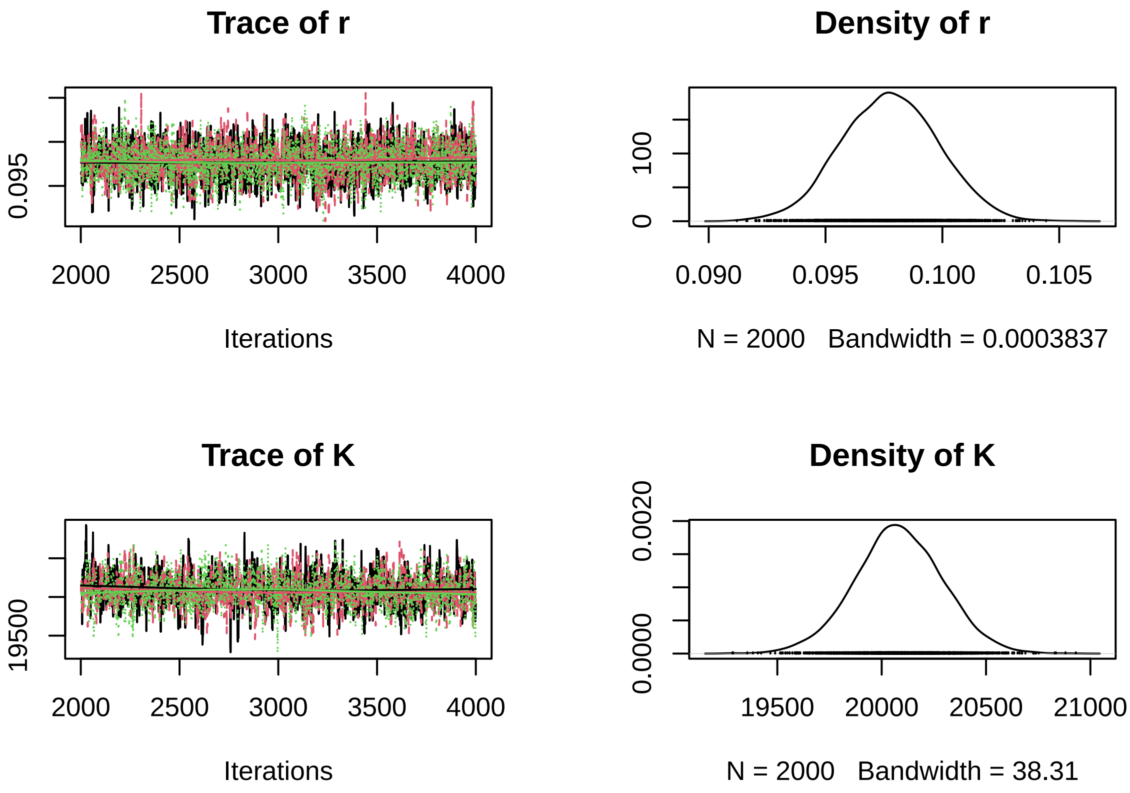

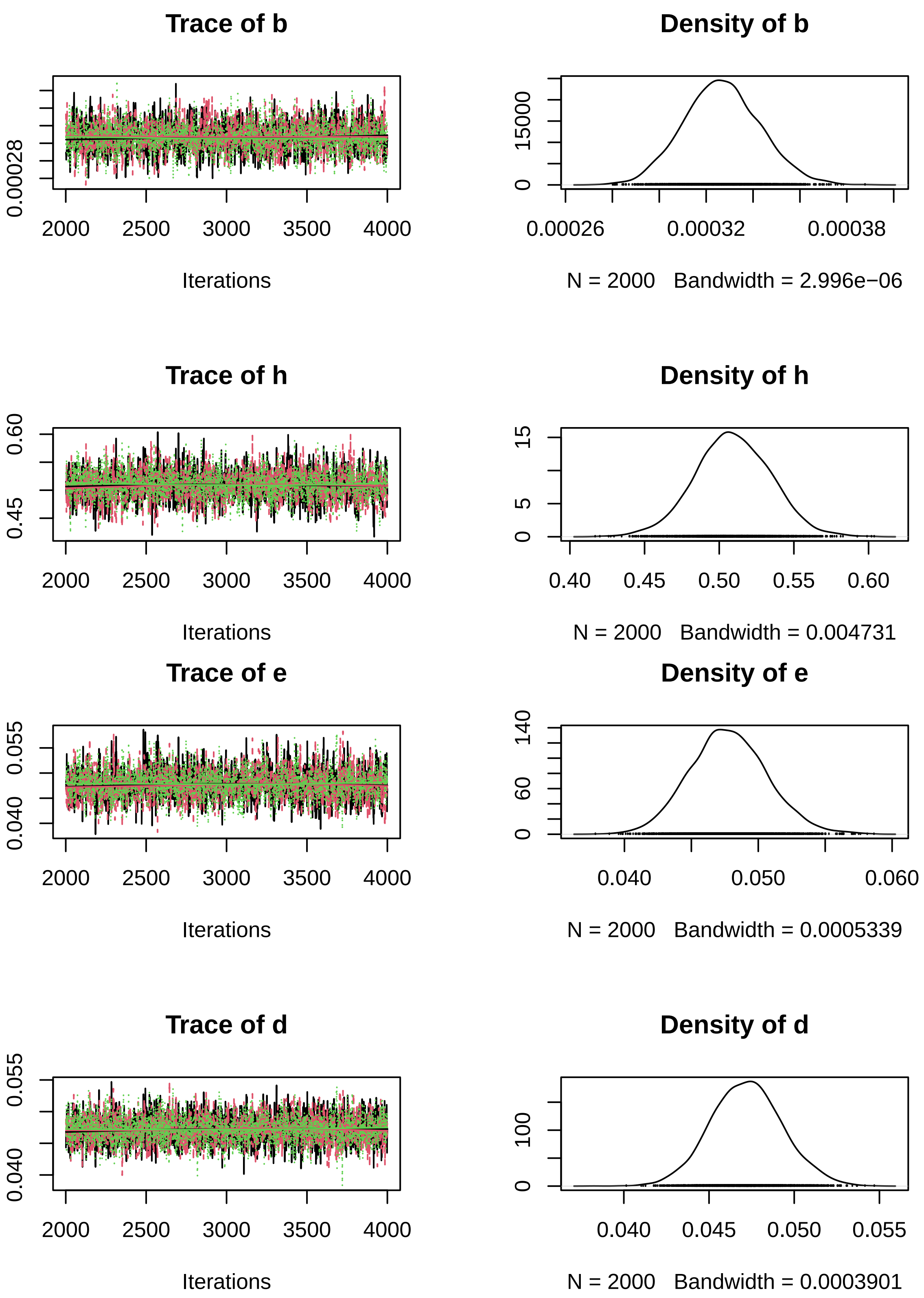


A pairs plot can detect correlations in model parameters. Almost perfect correlation would indicate non-identifiability.

```
pairs(fit, pars=c("r","K","b","h","e","d"))
```

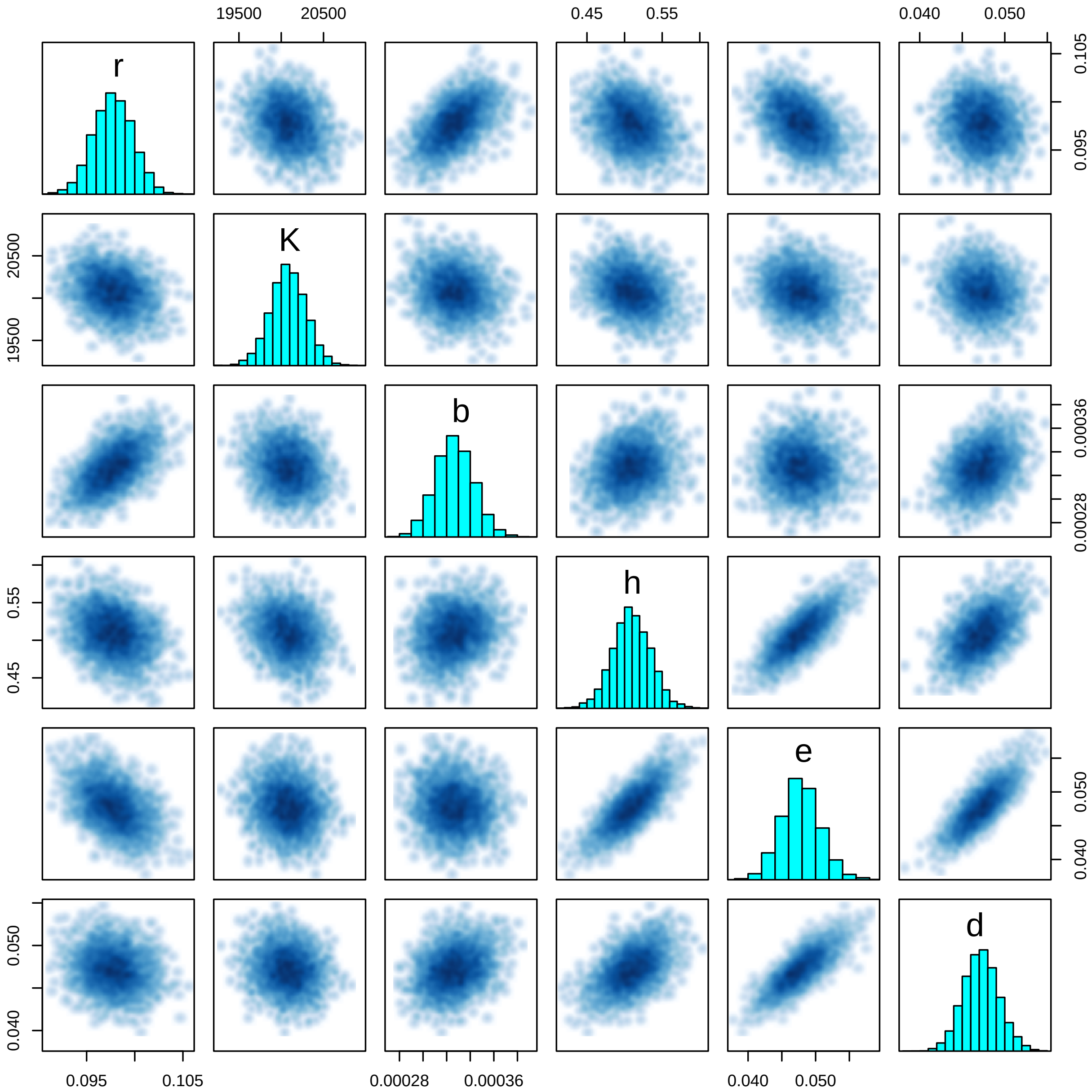


#### 6.6 Posterior predictions

Since underlying true states (`N1true[i,j]`,`N2true[i,j]`) for all observations (`N1[i,j]`,`N2[i,j]`) are already estimated, we can directly pull their means, medians or CIs from the summary table. We plot them against the data, for simplicity as piecewise linear interpolations between observations.

Here, we only show the 2-species mixtures and omit the control time series.

```
summary = summary(fit)$summary

head(summary)
```

```
##              mean      se_mean           sd         2.5%          25%          50%
## r 9.776301e-02 4.525458e-05 2.062021e-03 9.376568e-02 9.634572e-02 9.774709e-02
## K 2.008200e+04 6.222803e+00 2.100550e+02 1.966380e+04 1.994609e+04 2.007965e+04
## b 3.267130e-04 3.608593e-07 1.624295e-05 2.959818e-04 3.156921e-04 3.263333e-04
## h 5.093018e-01 5.796770e-04 2.542771e-02 4.589295e-01 4.922962e-01 5.088279e-01
## e 4.779115e-02 6.786151e-05 2.900815e-03 4.232692e-02 4.587146e-02 4.771600e-02
## d 4.721738e-02 3.976372e-05 2.101364e-03 4.314712e-02 4.577906e-02 4.720542e-02
##               75%        97.5%    n_eff     Rhat
## r 9.917559e-02 1.017926e-01 2076.162 1.002516
```

```
## K 2.022197e+04 2.049913e+04 1139.447 1.002623
## b 3.372672e-04 3.591763e-04 2026.069 1.000942
## h 5.265050e-01 5.585075e-01 1924.165 1.001105
## e 4.971652e-02 5.376865e-02 1827.230 1.001152
## d 4.858845e-02 5.148482e-02 2792.728 1.000623
```

```
N1.pred.qs = matrix(NA, nrow=3, ncol=length(data$t))
N2.pred.qs = matrix(NA, nrow=3, ncol=length(data$t))

par(mfrow=c(3,2), mar=c(3,3,0,0), oma=c(3,3,1,1))

for(i in 1:m){
  for(j in 1:length(times)){
    N1.pred.qs[, j] = summary[paste0("N1true[",i,",",j,"]"), c("2.5%", "50%", "97.5%") ]
    N2.pred.qs[, j] = summary[paste0("N2true[",i,",",j,"]"), c("2.5%", "50%", "97.5%") ]
  }

  plot(data$t, data$N1[i, ]/data$f,
       ylim=c(1e2,2e4), type="n", xlab="", ylab="", log="y")

  polygon(c(data$t, rev(data$t)), c(N1.pred.qs[1, ], rev(N1.pred.qs[3, ])),
          col = adjustcolor("red",alpha.f=0.25),  border = NA)
  lines(data$t, N1.pred.qs[2, ], col="red", lwd=1.5)
  points(data$t, data$N1[i, ]/data$f)

  polygon(c(data$t, rev(data$t)), c(N2.pred.qs[1, ], rev(N2.pred.qs[3, ])),
          col = adjustcolor("blue",alpha.f=0.25),  border = NA)
  lines(data$t, N2.pred.qs[2, ], col="blue", lwd=1.5)
  points(data$t, data$N2[i, ]/data$f)

  if(i==1) legend("topleft", c("res","con"), lwd=c(2,2), col=c("red","blue"))

}
mtext("Time [h]", side=1, outer=TRUE, line=1)
mtext("Abundance [ind], logscale", side=2, outer=TRUE, line=1)
```

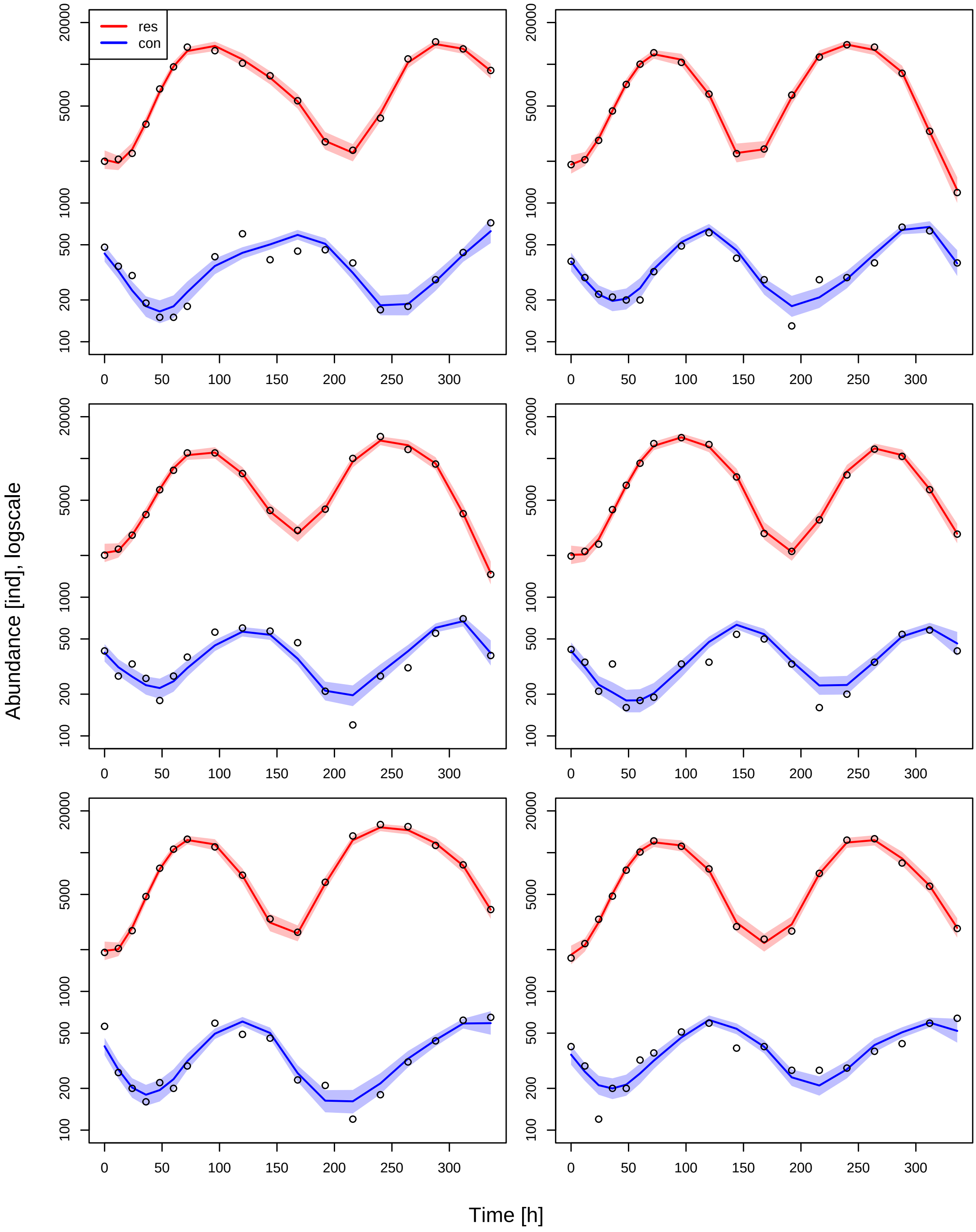
res
con
Abundance [ind], logscale
Time [h]
100
200
500
1000
5000
20000
0
50
100
150
200
250
300
